# Variable efficiency of nonsense-mediated mRNA decay across human tissues, tumors and individuals

**DOI:** 10.1101/2024.02.29.582778

**Authors:** Guillermo Palou-Márquez, Fran Supek

## Abstract

Nonsense-mediated mRNA decay (NMD) is a quality-control pathway that degrades mRNA bearing premature termination codons (PTCs) resulting from mutation or mis-splicing, and that additionally participates in gene regulation of unmutated transcripts. We analyzed ∼10,000 exomes and ∼27,000 transcriptomes from human tumors and healthy tissues to quantify individual-level NMD efficiency, and assess its variability between tissues and between individuals. This was done by monitoring allele-specific expression of germline PTCs, and independently supported by mRNA levels of endogenous NMD target transcripts. Nervous system and reproductive system tissues have lower NMD efficiency than other tissues such as the digestive tract. Next, there is considerable systematic inter-individual variability in NMD efficiency, and we identify two underlying mechanisms. First, in cancers there are somatic copy number alterations that robustly associate with NMD efficiency, prominently the commonly-occurring gain at chromosome 1q that encompasses two core NMD genes: *SMG5* and *SMG7* and additional functionally interacting genes such as *PMF1* and *GON4L*. Second, loss-of-function germline variants in various genes such as the *KDM6B* chromatin modifier can associate with higher or lower NMD efficiency in individuals, affecting different tissues thereof. Variable NMD efficiency should have clinical implications as it modulates positive selection upon somatic nonsense mutations in tumor suppressor genes, and is associated with survival of cancer patients, with relevance to predicting immunotherapy responses across cancer types.

## 1. Introduction

Nonsense-mediated mRNA decay (NMD) is a quality-control mechanism that degrades mRNAs containing premature termination codons (PTCs), which can arise from nonsense mutations, frameshift mutations, or aberrant splicing events^1–3^. Traditionally recognized for its role in preventing the translation of defective transcripts that could lead to truncated proteins with potentially detrimental functions, NMD also plays a common role in the regulation of non-mutated, naturally occurring transcripts. It is estimated that NMD regulates approximately 10% of the normal transcriptome, impacting physiological gene expression^4–6^. This broader regulatory role is attributed to the presence of NMD-triggering features in many endogenous transcripts, even in the absence of PTCs^7^. An additional regulatory role of NMD stems from being coupled with alternative splicing that retains introns or skips exons and thus introduces NMD-triggering features as a means to regulate gene expression^7^.

In mammals, the NMD pathway primarily distinguishes between PTCs and normal stop codons through a mechanism linked to the splicing process and translation. The ribosome removes the exon-junction complex (EJC) proteins deposited onto splice sites as elongation progresses. The residual EJCs downstream of a PTC signal the initiation of NMD; thus, the position of the PTC along the mRNA determines NMD efficiency upon that variant. Based on various targeted experiments that yielded this mechanistic understanding, as well as on large-scale genomic analyses of PTCs and transcriptomes, key genomic rules for NMD evasion were identified ^8–10^. Most prominently, the PTCs located in the last exon of a transcript escape NMD, as well as PTCs within the final ∼50 nt of the penultimate exon. Start-proximal PTCs are also less likely to initiate NMD, perhaps in part through reinitiation of translation, and PTCs in long exons also partially evade NMD through less clear mechanisms^8,11^. These rules notwithstanding, the majority of PTCs trigger NMD to some extent.

Furthermore, non-mutated endogenous NMD targets exhibit specific features that can trigger NMD^5,6,12–14^. For example, an upstream open reading frame (uORF) in the 5’ untranslated region (UTR) can cause NMD activation^15,16^, mimicking a PTC with a downstream EJC deposited, and similarly so an intron within the 3’ UTR^12,17^. High GC content in 3’UTR may also be associated with increased NMD efficiency^18^.

However, the predictive accuracy of NMD features is not perfect: around half of PTCs in NMD-triggering regions exhibit a lower NMD degradation^11^, suggesting additional rules for NMD evasion. Similarly, many endogenous (i.e. not mutated) transcripts known to be NMD targets lack known NMD-triggering features^12,13^. This indicates that there are still more genomic rules to discover, however these yet-to-be-discovered rules of NMD efficiency acting upon particular PTCs likely have quantitatively minor effects, compared to known rules^8^. Instead, it appears that additional features contribute to NMD variation, and these might not pertain only to different PTCs or transcripts due to their genomic context, but were also proposed to vary among individuals^19,20^. Particular examples underlying this hypothesized inter-individual NMD variability put forth on links with germline variation, somatic alterations or changes in expression of NMD factors^20–22^. Furthermore, though less extensively studied, variations in NMD efficiency have been observed between different tissues and cell lines in mice^23,24^ and in human cells^25,26^. For example, considering the same PTCs across various human tissues, they showed that 18% of variants exhibited heterogeneous effects across tissues and 8% were tissue specific^27^. These variations in NMD efficiency between tissues or across individuals can lead to distinct pathologies in genetic diseases and affect therapeutic outcomes. This is illustrated in the case of two Duchenne muscular dystrophy patients with identical PTCs but differing disease severities^28^ associated with NMD efficiency. Another case involved two fetuses harboring the same PTC for Roberts syndrome^29^, thus particular examples suggest individuals can differ in NMD efficiency but it is not clear what is the magnitude of this variation and its extent across populations, which are the common mechanisms nor the implications to genetic disease.

One genetic disease with many established links with NMD is cancer. NMD may exhibit either pro-tumor and tumor suppressor functions, as a function of cell type, and possibly the genetic context and tumor microenvironment^30^. For example, a reduction in NMD activity, as suggested by downregulation of the key factor *UPF1*, has been reported in various cancers^31–37^, in comparison with matched normal tissue; however in other studies of cancerous tissues an overexpression of *UPF1* was reported instead^38,39^. A notable case was found in colorectal adenocarcinoma cancer (CRAD), where patients with microsatellite-instability (MSI) hypermutation exhibited overexpression of essential NMD factors such as *UPF1/2* and *SMG1/6/7*, in contrast to microsatellite-stable (MSS) primary CRAD^40^. Moreover, inhibiting NMD impaired cell proliferation and tumor growth in MSI but not MSS^40^. The above collectively suggests the importance of modulating NMD activity during tumor evolution to promote growth.

Approximately 30% of inherited diseases, as well as the majority of cancers with mutations in tumor suppressor genes (TSGs), are caused by PTCs resulting from frameshift mutations or nonsense variants^41,42^, and thus may be modulated by NMD. NMD has been recognized as a significant factor influencing disease severity and clinical phenotype^11,28,43–45^, and a population genomics analysis suggests that it can either exacerbate or alleviate disease phenotypes, depending on the disease, with the former case being the more common^11^.

Additionally, NMD was considered in therapeutic contexts: it is a determinant in the efficacy of cancer immunotherapy, where only those frameshift mutations that evade NMD predict a positive immunotherapy response^11,44^. Mouse cancer models of genetic NMD targeting show favorable responses^46^. In the context of genetic diseases, it has been shown that NMD inhibition markedly increases the levels of mutated mRNAs^47^, conceivably ameliorating the clinical phenotype severity, especially in combination with stop codon read-through therapy drugs^48–50^.

This underlines the importance of considering NMD in the clinic, in cancer and potentially in other genetic diseases, as well as investigating NMD inhibition as supplementary therapy (reviewed by Supek et. al^1^). If there were variability in NMD efficiency across individuals and/or tissues, it would impact the disease phenotype and potentially options for treatments, further motivating large-scale studies of differences in NMD efficiency.

Despite reports of individual cases, a systematic study of NMD efficiency across human individuals and tissues is lacking, as well as the understanding of its mechanistic basis. Here, we address this gap by assessing NMD-triggering and evading features genome-wide, for which we deployed two methodologies for estimating NMD efficiency, measured globally for each individual and compared across individuals or tissues (see schematic from Fig. 1). The first method draws on allele-specific expression (ASE) of PTC-bearing transcripts, and the second method assesses expression of endogenous NMD target transcripts. We report significant variation in NMD efficiency between individuals, considering both the normal tissues, as well as various cancer types arising in these individuals. We have identified somatic copy number alterations (CNAs) and germline variants that explain some of this variability in NMD efficiency. We further suggest significant differences between somatic tissues, with lower NMD efficiency in the nervous and reproductive system tissues. Furthermore, we explored the impact of individual NMD efficiency on the somatic evolution processes acting on tumor suppressor genes, as well as survival outcomes and immunotherapy response in different cancer types, suggesting the clinical relevance of NMD in cancer.

**Fig. 1.**
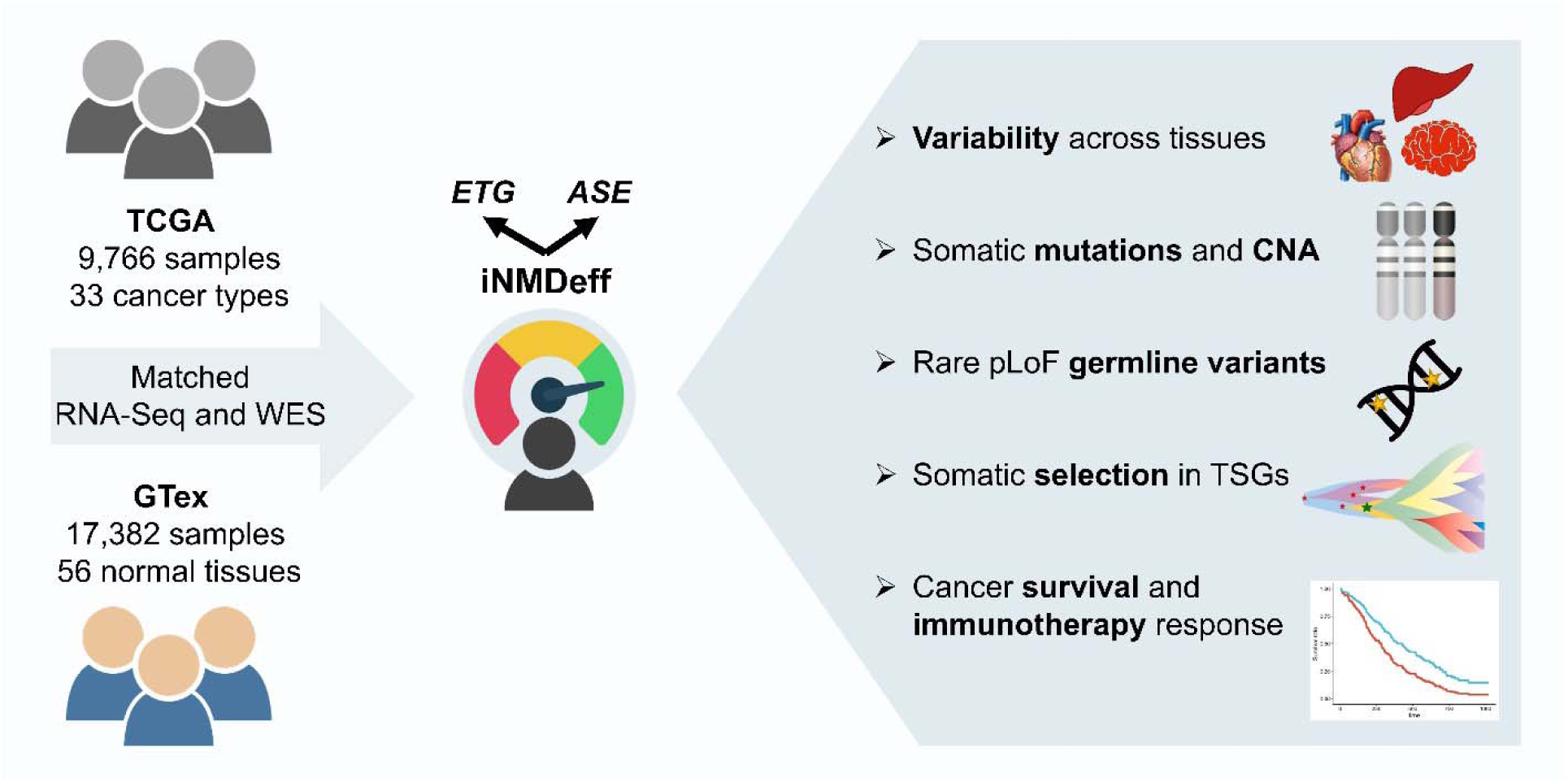
Exploring variability and identifying determinants of NMD efficiency across tissues, tumors, and individuals.

## 2. Results

### Individual-level quantification of NMD efficiency

We quantified the individual-level NMD efficiency (iNMDeff) by utilizing matched RNA-seq and whole-exome sequencing (WES) data from two cohorts: i) the tumor tissues in The Cancer Genome Atlas (TCGA), comprising 9,766 tumor samples from 33 different cancer types, and ii) the normal tissues from the Genotype-Tissue expression (GTex) consortium, consisting of 979 individuals across 56 tissues, totaling 17,382 samples (with a median of 19 tissues per donor).

We devised two orthogonal methodologies to estimate iNMDeff in each tumor or normal sample (Supp. Fig. S1A). Both are implemented using a negative binomial regression (see Methods) to model either transcript counts or allele expression counts, comparing two groups: NMD targets or controls. Each method was subjected to rigorous filters to ensure that the set of transcripts/variants and individuals used were less confounded by other sources of variability (see Methods).

#### Endogenous target gene (ETG) method

We started with an initial set of genes that had been experimentally identified as endogenous NMD target genes from various studies that perturbed NMD activity in human cell lines^5,6,13,14,51^ (see Methods). For each gene, we categorized its transcripts into NMD targets and controls based on computationally predicted NMD-triggering features (see Methods and Supp. Fig. S1B). Here, we relied on two known features: i) uORF at the 5’ UTR, and ii) at least one splice site (or EJC) within the 3’ UTR. Next, for each NMD target gene, we selected two transcripts, one NMD target and the other as a NMD non-target, serving as an internal control. Our iNMDeff is the coefficient of the indicator variable stating whether the pooled transcripts within an individual are either NMD target (1) or control (0), from a negative binomial regression (see Methods); the interpretation is that it is a negative log (base e) ratio of the raw expression levels of the NMD target transcripts divided by the control transcripts.

We calculated the iNMDeff for the different NMD gene sets from each study separately (see Methods): “Karousis”^13^, “Colombo”^5^, “Tani”^6^, and “Courtney”^51^, and we further tagged the transcripts as “nonsense_mediated_decay” from the Ensembl gene annotation as an additional “Ensembl’’ set. Additionally, genes found in at least 2 independent studies constituted a “NMD Consensus” gene set, and the union of genes across all studies was the “NMD All” gene set. For our control groups, we selected two sets of genes at random (excluding genes reported as NMD targets in all of the experimental studies mentioned above), selecting from genes with and without NMD-triggering features in their transcripts: “RandomGenes with NMD features”, and “RandomGenes without NMD features”. The former should behave similarly as the experimental NMD gene sets but is based only on computational prediction, while the latter is a negative control gene set.

#### Allele-specific expression (ASE) method

We used exonic coding germline variants (population minor allele frequency, MAF, < 20%) to define three variant sets: i) “NMD-triggering PTCs” resulting from nonsense variants and from frameshifting indels; ii) “NMD-evading PTCs” resulting from nonsense and indels; iii) synonymous variants. The latter two were used as negative control sets. For each variant, the mutated allele counts in RNA-seq reads were used as NMD targets and the wild-type allele counts at the same locus were used as controls. We calculated the iNMDeff for each individual using the three aforementioned NMD variant sets separately. Similarly as in ETG, all variants within the individual were pooled to estimate the iNMDeff, obtained from the regression coefficient of the indicator variable stating whether the variants are either mutated (1) or wild-type (0) (see Methods). The interpretation is that it is a negative log (base e) ratio of the raw RNA-seq allele counts of the mutant vs wild-type variants.

In summary, we computed the NMD efficiency for each individual multiple times, varying the gene/variant set used, in TCGA and GTex cohorts: including the negative controls this amounted to 11 NMD gene sets for the ETG method, and 3 NMD variant sets for the ASE method. Henceforth, the ETG gene sets and the ASE variant sets will be collectively referred to as “NMD sets” for brevity.

#### Robustness of the individual NMD efficiency estimates

Given the high correlation between ETG iNMDeff across different NMD gene sets (Supp. Fig. S3E-F) -- except, encouragingly, for the negative control lacking NMD-triggering features -- we opted to focus on the stringent “NMD Consensus” and the permissive “NMD All”, while the iNMDeff for the rest of the NMD gene sets are detailed in Supp. Fig. S2A-C. In TCGA, the iNMDeff estimates from the ETG method (Fig. 2A) and from the ASE method (Fig. 2B) show various trends that imply they are reliable. For instance, ETG estimates of iNMDeff are similarly efficient across the “Consensus” and “All” NMD gene sets, and as anticipated the higher-confidence “NMD Consensus” gene set displaying a readout of slightly higher efficiency than the more permissive “NMD All”, which presumably contains weaker NMD target genes (p < 2e-16, Mann-Whitney *U* test). Encouragingly, the negative control gene set “Random Genes without NMD features” shows negligible efficiency (p < 2e-16, Mann-Whitney *U* test, when compared to “NMD Consensus”), with the median close to zero (Fig. 2A). Underscoring the reliability of the features we used to identify NMD-targeted transcripts for the ETG method, the gene set “Random Genes with NMD features” exhibits efficiency similar to the experimentally identified NMD gene sets, with a very slight albeit significant difference (median iNMDeff = 2.10 vs 2.11, p = 4.8e-2, Mann-Whitney *U* test, when comparing to “NMD Consensus”). This result also suggests the presence of many additional genes with at least one NMD-targeted transcript not identified experimentally in studies considered here^5,6,13,51^. In GTex, we saw similar results as described above (Supp. Fig. S2B-C).

**Fig. 2.**
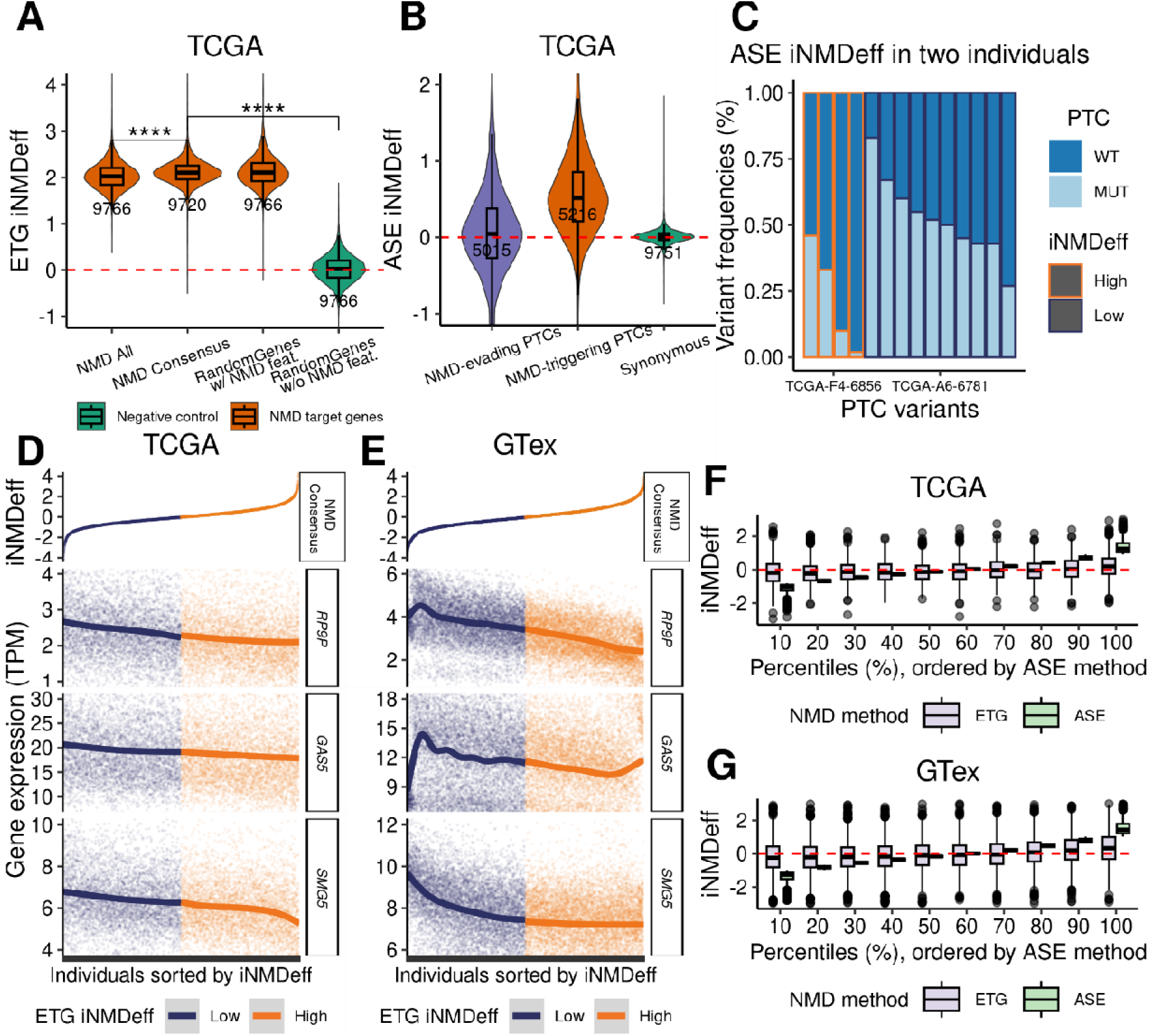
Individual-level quantification of NMD efficiency. **A**, Estimation of individual-level NMD efficiency, iNMDeff (Y-axis), using the ETG method across 9,766 TCGA tumor samples, showcasing three NMD gene sets, one being a random gene set with NMD-triggering features, and a random gene set without NMD features as control. **B**, iNMDeff estimations using the ASE method for two NMD variant sets, alongside a non-NMD variant set as a control. ****p□<□0.0001, by two-sided Mann–Whitney U test for A and B. **C**, Variability in ASE iNMDeff between two example individuals from TCGA is depicted through the frequencies of wild-type (WT) and mutated (MUT) variants (Y-axis) for different PTCs (X-axis). The individual with high iNMDeff (orange) has lower variant frequency of the MUT allele, and vice versa for the low iNMDeff individual (magenta). **D-E**, Gene expression levels (TPM) of two NMD target genes, RP9P and GAS5, and an NMD factor, SMG5, compared against the ETG iNMDeff (using the “NMD Consensus” gene set) sorted from lowest to highest along the X-axis in TCGA (D) and GTex (E, n = 17,382). Samples are stratified by the median ETG iNMDeff in high and low. **F-G**, Stratification of iNMDeff (Y-axis) into ten percentiles (X-axis) for both ASE and ETG methods, ordered by the ASE method, for TCGA (F) and GTex (G).

To further support the concept of ETG method for estimating iNMDeff, we compared the iNMDeff estimates (from “NMD Consensus”) with the gene expression of two well-known NMD target genes, *RP9P* and *GAS5*^5^; of note, we excluded them from our methods to avoid circularity in this analysis. In this context, individuals with high iNMDeff typically show lower gene expression of both *RP9P* and *GAS5* in tumor samples from TCGA (Fig. 2D) and healthy tissue samples from GTex (Fig. 2E). Additionally, the gene expression of key NMD factors were analyzed. For NMD genes *UPF1*, *UPF2*, *UPF3A*, *UPF3B*, *SMG1*, *SMG5*, *SMG6*, *SMG7* and *SMG9*, but not *SMG8*, a similar pattern to *RP9P* and *GAS5* was observed in both cohorts (Supp. Fig. S2D-E). This is consistent with the negative feedback loop and cell-type-specific inherent in NMD factors, where inhibition of NMD by siRNA caused an upregulation of the mRNAs of all key NMD genes except for *UPF3A*, *SMG8* and *SMG9* (these last two were not tested)^52^. The autoregulation is maintained through internal NMD-triggering features in the various NMD factors^52^. This correlation was less notable or even absent for EJC genes *MAGOH, RBM8A, EIF4A3* and *CASC3* (Supp. Fig. S2F-G).

Next, as a test of reliability of the ASE iNMDeff estimates, we consider the differences in distribution of efficiency across the 3 sets of variants. While ASE-derived individual level NMD efficiency is observable when calculated using “NMD-triggering PTCs”, when these estimates are derived from negative control variant sets, such as “NMD-evading PTCs” or “Synonymous”, the efficiency drops to almost negligible levels (p < 2e-16 for both variant sets, Mann-Whitney *U* test, when comparing to “NMD-triggering PTCs”).

To illustrate the ASE iNMDeff estimation, we present a representative example of raw allele-specific expression counts from PTC variants in two COAD_MSI individuals from TCGA (Fig. 2C). For individual “TCGA-F4-6856”, who was labeled as having a high iNMDeff, we observe four PTC variants with notably low RNA-seq counts of their alternative alleles, relative to the reference allele. This is consistent with the rapid degradation characteristic of an actively engaged NMD pathway. In contrast, the individual “TCGA-A6-6781”, who exhibits a low iNMDeff, is characterized by ten PTC variants where the counts of reference and alternative alleles are proportionally balanced, consistent with what is typically seen in a heterozygous variant. This pattern of a consistent lack of ASE across 10 PTCs implies a global reduction in the activity of the NMD pathway in that individual (at least in the sampled tissue), affecting all variants therein.

#### Agreement between the two methods to estimate NMD efficiency

If both ETG and ASE methods are reflecting the true NMD activity of the individual, then a correlation between the iNMDeff estimates would be anticipated. We proceeded to estimate pan-cancer and pan-tissue correlations between iNMDeff for ASE (using “NMD-triggering PTCs” variant set) and ETG (using “NMD Consensus” set) (Supp. Fig. S3A-B), revealing a statistically significant correlation (Pearson p < 2-16 in both cases). The same trend was observed when correlations were calculated for each tissue or cancer stratified into subtypes, with positive correlations in 76 out of the 101 tested tissues/cancers (Supp. Fig. S3C-D). That ETG and ASE method agree well is evident upon stratifying the various samples by bins of iNMDeff. The top decile (samples with lowest ASE iNMDeff) of TCGA patients has a median scaled ETG iNMDeff of −0.19, while the bottom decile (samples with highest ASE iNMDeff) has a median scaled ETG iNMDeff of 0.20 (difference at p = 4.12e-13 by Mann-Whitney *U* test, Fig. 2F). Similarly, in GTex the top ASE decile has a median ETG iNMDeff of −0.25 while the bottom ASE decile has ETG iNMDeff = 0.34 (p < 2e-16, Fig. 2G).

We also tested correlations between ASE iNMDeff and ETG iNMDeff and other biological covariates or technical variables: age, sex, tumor mutation burden (TMB), tumor indel burden (TIB), tumor nonsense burden (TNB), purity, MSI score, RNA-seq sample library size, leukocyte fraction, CNA burden, and number of NMD targets used to estimate iNMDeff (either PTCs for ASE, or transcripts for ETG) (Supp. Fig. S3E-F). The tested covariates did not show significant correlations with any of our iNMDeff in either TCGA or GTex. The “NMD Consensus” gene set correlates the highest to ASE iNMDeff (Supp. Fig. S3A-B and S3E-F), and thus we used it as the default gene set for the ETG method.

In summary, we developed two genomic methodologies for quantifying individual-level NMD efficiency - the ASE and ETG methods - which rely on RNA-seq signal in germline “NMD-triggering PTCs” variants, and in the “NMD Consensus” experimentally determined set of NMD target genes for ASE and ETG, respectively. We demonstrated the robustness of these methods by various approaches, allowing us to use the ASE and ETG method to investigate potential variation in iNMDeff across different human tissues and cancer types.

### Significant variability in NMD efficiency across human tissues and individuals

#### Inter-tissue variability of NMD efficiency

Next, we applied our data set of iNMDeff estimates for 27,148 different tumoral and normal tissue samples to rigorously test the hypothesis that NMD efficiency varies across human tissues. We grouped the iNMDeff estimates by cancer types in TCGA (Supp. Fig. S4A), and by type of normal tissue in GTex (Supp. Fig. S4B), and tested for differences between the tissues using a randomization test (see Methods for details and schematic from Fig. 3A). The test statistic we used, termed “Inter-Tissue iNMDeff Variability Deviation” (ITNVD), is a measure of how much the iNMDeff differences between tissue medians deviate from expectation if the iNMDeff values were randomly distributed across samples from different tissues. If there is no tissue-specific variability in iNMDeff, the value should be close to 0.

**Fig. 3.**
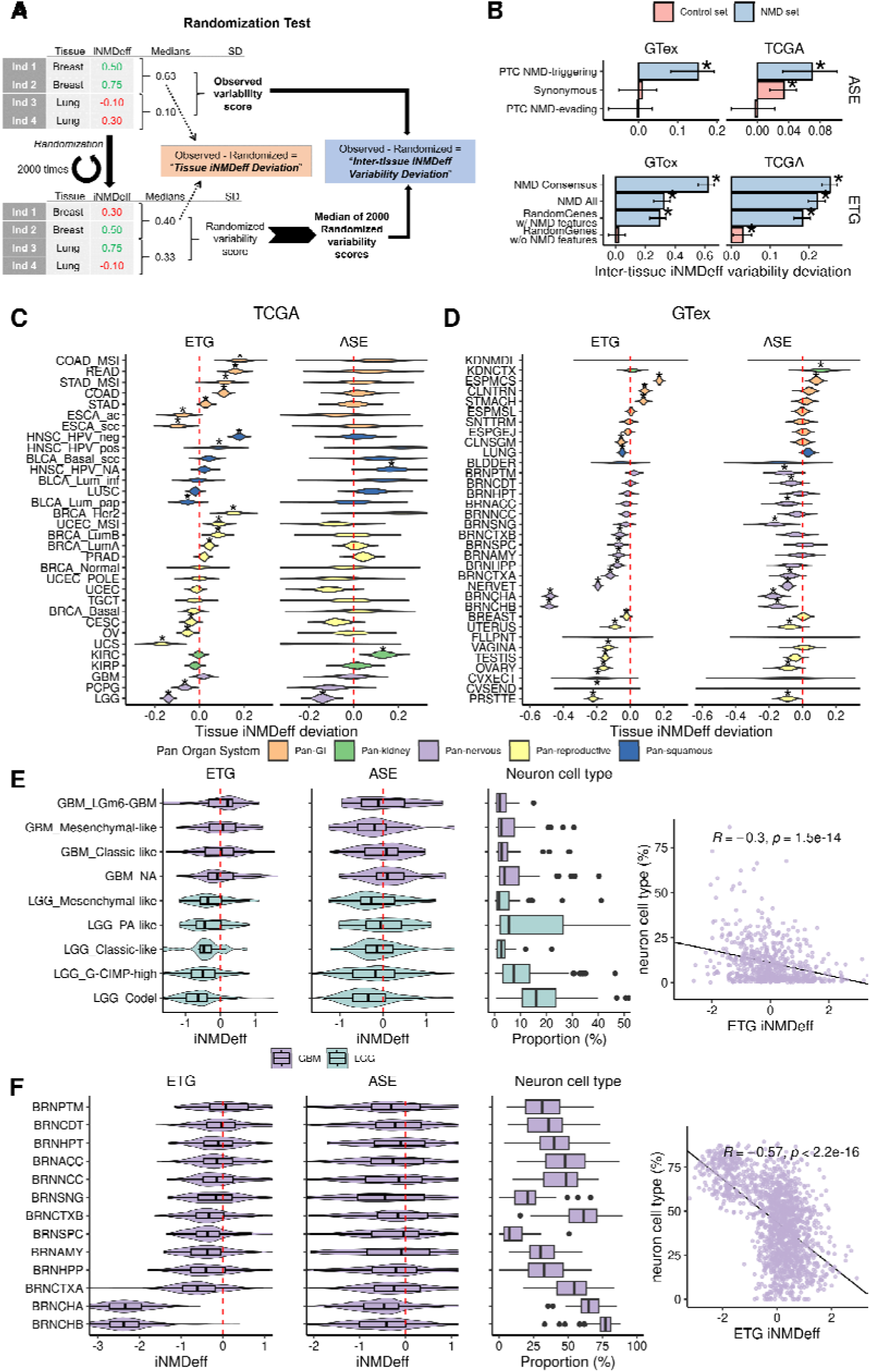
Significant variability in NMD efficiency across human tumors, normal tissues and individuals. **A**, Schematic of the randomization tests, illustrating the methodologies for assessing NMD efficiency variability: Tissue iNMDeff Deviation (TND) and Inter-tissue iNMDeff Variability Deviation (ITNVD). **B**, Displays ITNVD test scores (X-axis) for each NMD gene set in the ETG method (bottom panel) and NMD variant set in the ASE method (top panel). Scores for non-NMD negative control sets are also shown for comparison. Data are presented for both GTex (left) and TCGA (right) cohorts. Positive scores indicate variability in iNMDeff across tissues or cancer types, with stars (*) denoting statistical significance for tissue specificity at p ≤ 0.05, by randomization test. **C-D**, Shows TND test scores (X-axis) for each cancer type within TCGA (C) and each normal tissue within GTex (D), providing insight into the specific variations of NMD efficiency within tissues. Tissues are grouped based on cell-of-origin: Nervous system-related tissues (Pan-nervous), Kidney-related tissues (Pan-kidney), Reproductive system tissues (Pan-reproductive), Gastrointestinal tissues (Pan-GI), and those originating from Squamous cells (Pan-squamous). The groups of tissues were ordered based on the median TND scores, arranging them from the highest to the lowest median scores, top to bottom. Complete names of tissue acronyms can be found in Supp. Table S2. **E**, Analyzes scaled ETG and ASE iNMDeff (X-axis) in low grade glioma (LGG) and glioblastoma (GBM) from TCGA, categorized by genetic or histological subtypes (first two panels). Also displays neuron cell type proportions derived from bulk RNA-seq deconvolution (third panel) and their Pearson correlation (r) with ETG iNMDeff (fourth panel), indicating the relationship between cellular composition and NMD efficiency. **F,** Mirrors the analysis in E but focuses on GTex brain tissues, stratified by different subregions, showcasing the variability of NMD efficiency in normal brain tissue contexts and its correlation with neuron cell type proportions.

The test was applied across both ASE and ETG estimated iNMDeff in the TCGA and GTex cohorts. We observed a significant, albeit modest, variability deviation across both cancerous tissues in TCGA, and across normal tissues in GTex (Fig. 3B and Supp. Fig. S4C). Notably, the NMD gene/variant sets exhibited a much higher variability deviation than the negative control gene/variant sets, with a mean ITNVD of 0.27 versus 0.01 (p = 3.5e-3, t-test). Specifically, the ETG “NMD Consensus” set demonstrated the largest deviation (ITNVD = 0.62 in GTex and 0.26 in TCGA, both with p = 5e-4), and this was replicated in the ASE “NMD-triggering PTCs” variant set (ITNVD = 0.15 in GTex with p = 3e-3, and ITNVD = 0.07 in TCGA with p = 5.3e-3). In contrast, the variability deviation in the negative control sets was negligible: the ASE of “Synonymous” variant set showed a ITNVD of 0.01 in GTex and 0.03 in TCGA, and the ETG “Random Genes without NMD-triggering features” set had a ITNVD of 0.02 in GTex and 0.03 in TCGA. These results suggest there is an important non-random iNMDeff variability associated with tissue and/or cell type identity, observed across both cancers and normal tissues.

A recent study suggested no differences in NMD efficiency between human tissues^53^. To reconcile this with our results we compared our ranking of tissues, ordered by the tissue median iNMDeff, with the ranking published by Teran et al. (2021)^53^, who used ASE on PTCs in GTex data. We found a strong correlation between their tissue rankings and our results in both our ASE (R = 0.75, p = 7e-10) and our ETG estimates (R = 0.72, p = 1.62e-8). Our analysis demonstrates this differential NMD efficiency across tissues is significant.

#### Intra-tissue variability of NMD efficiency

To identify which tissues and cancer types have the most and least efficient NMD (Supp. Fig. S4A-B), we focused on the variation in iNMDeff within each specific tissue, and devised a statistic of Tissue iNMDeff Deviation (TND), with a baseline derived from a randomization test. (see Methods and schematic in Fig. 3A). The resulting distribution of deviations (Fig. 3C-D) suggests the tissue variability in NMD efficiency, where positive TND values indicate that a tissue exhibits higher iNMDeff than what would be expected by chance, and vice versa for negative values. Notably, 38 out of 54 tissues, and 19 out of 32 cancer types show a significant TND (*p* < 0.05 by randomization test) by the ETG method (for brevity, a selection is shown in Fig. 3C-D, and the complete list is in Supp. Fig. S5A-B). The test for tissue-specificity of NMD efficiency replicates for 43% of these tissues in the ASE method as well. Next, we have categorized these into five broad primary groups for ease of analysis and interpretation: Nervous system-related tissues (Pan-nervous), Kidney-related tissues (Pan-kidney), Reproductive system tissues (Pan-reproductive), Gastrointestinal tissues (Pan-GI), and those originating from Squamous cells (Pan-squamous), as denoted in a previous study^54^.

The Pan-GI group of tissues displayed the highest iNMDeff in comparison to randomized values, i.e. positive TND, across both the ETG and ASE NMD methods in both TCGA and GTex cohorts. Specifically in TCGA (Fig. 3C), colon, stomach, and rectum adenocarcinomas (COAD, STAD, READ, respectively) exhibited significantly higher ETG iNMDeff than random (TND varying 0.03 - 0.16 for the 3 GI tissues, p = 3.47e-2 - 1.23e-3). A similar pattern was observed in the GTex cohort (Fig. 3D) for normal colon (TND = 0.09, p = 9.71e-4) and stomach (TND = 0.08, p = 9.71e-4) tissues. Notably, MSI cancer subtypes, such as COAD_MSI and STAD_MSI, demonstrated higher iNMDeff compared to their non-MSI counterparts (Supp. Fig. S5C-D), placing them among the cancers with the highest ETG iNMDeff (TND = 0.18, p = 1.23e-3 and TND = 0.12, p = 2.29e-3, respectively). Esophagus mucosa (ESPMCS) normal tissue displayed significantly higher iNMDeff in both ETG (TND = 0.17, p = 9.71e-4) and ASE methods (TND = 0.08, p = 2.75e-3), however, esophageal cancer showed a lower iNMDeff for both squamous cell carcinoma and adenocarcinoma (ESCA_scc and ESCA_ac, respectively; Fig. 3C) types, suggesting that some cancer types might undergo a change in NMD efficiency during transformation from normal tissue.

In contrast to the Pan-GI, tissues classified as Pan-reproductive generally exhibited lower iNMDeff (Supp. Fig. S4A-B) than expected based on randomization, i.e. negative TND. This trend was consistent across both TCGA and GTex cohorts and observed in both ETG and ASE methods. For example, in GTex’s Pan-reproductive normal tissues: ovary, cervix ectocervix (CVXECT), and cervix endocervix (CVSEND) all showed lower iNMDeff in ETG (TND = −0.16, - 0.19 and −0.20; p = 9.7e-4, 8.7e-3 and 8.7e-3, respectively) and in ASE this replicated for ovary (TND = −0.09, p = 2.2e-2). A similar trend was observed in TCGA, with ovarian serous cystadenocarcinoma (OV, ETG TND = −0.05, p = 1.23e-3, ASE TND = −0.02, p = 4.7e-1) and cervical squamous cell carcinoma and endocervical adenocarcinoma also exhibiting lower iNMDeff in both methods (CESC, ETG TND = −0.04, p = 1.4e-2, ASE TND = −0.08, p = 0.12).

Other normal Pan-reproductive tissues like vagina, testis, uterus, and prostate (PRSTTE) showed strong significant differences in iNMDeff in ETG and ASE methods in GTex (Fig. 3D). These differences were not significant in their respective cancer types in TCGA, though they generally followed the same downward trend, with exception of prostate. Interestingly, uterine corpus endometrial carcinoma, MSI subtype (UCEC_MSI) showed significantly higher ETG iNMDeff (TND = 0.09, p = 1.23e-3), higher than the MSS subtype (Supp. Fig. S5C-D), paralleling the MSI trends observed previously in COAD and STAD.

For breast normal tissue, a slightly lower trend towards lower iNMDeff was observed in ETG (non-significant in ASE), aligning with trends in TCGA for breast invasive carcinoma in the basal subtype and the normal-like subtype (Fig. 3C), but not in the other subtypes (BRCA_LumA, BRCA_LumB, BRCA_Her2). This suggests that cancer types with very diverse subtypes, such as breast cancer, may display differences in NMD efficiency between subtypes.

Analysis of Pan-squamous and Pan-kidney tissues overall did not yield significant and/or consistent results across the two NMD methods, in either TCGA tumors or GTex tissues (Fig. 3C-D). Considering lung cancer in specific (LUSC in Fig. 3C and lung adenocarcinoma, LUAD, in Supp. Fig. S5A), there is no consistency between ETG and ASE NMD estimation methods. Of other tissues/cancers, we note a trend of higher iNMDeff for skin in both GTex and TCGA (melanoma), and lower iNMDeff in the thyroid in both GTex and TCGA (Supp. Fig. S5A-B).

As an additional approach to assess if NMD efficiency varies across tissues, we created a linear model predicting iNMDeff, including tissue or cancer type, and other covariates (see Methods). In this model, the removal of the cancer type/subtype variable significantly reduced the explained variability (R²), more so than removal of any of the other variables (Supp. Table S1). This was evident in both ETG and ASE methods, where the full model explained 17.1% and 58.7% of variability in TCGA and GTEx cohorts, respectively, dropping to 6.3% and 13%, respectively, upon the tissue variable removal.

#### Lower NMD efficiency in the tissues of the nervous system

Notably, the Pan-nervous group of tissues exhibited lower observed iNMDeff (Supp. Fig. S4A-B) compared to randomized expectations (Fig. 3D). In the GTex cohort, almost all brain subregions showed a significant reduction in iNMDeff, many with large effect sizes, as observed in the TND statistic in one or both NMD methods. Specific examples replicating in both iNMDeff methods include the cerebellum (BRNCHA, ETG TND = −0.48, p = 9.71e-4; ASE TND = −0.17, p = 2.8e-3), the cerebellar hemisphere (BRNCHB, −0.49, 9.7e-4; −0.15, 2.8e-3), along with the cortex (BRNCTXA, −0.12, 9.7e-4; −0.08, 1.4e-2), and further the frontal cortex BA9, hippocampus, and the substantia nigra show significance in one of the NMD estimation methods (ASE or ETG) and a consistent trend in the other method. Additionally, nervous-related tissues such as the pituitary (PTTARY) and nerve/tibial (NERVET) displayed reduced iNMDeff in both methods (Fig. 3D). In TCGA data, two brain cancer types that arise from glial cells are available for comparison, and the same significantly reduced iNMDeff was evident in low grade glioma (LGG, −0.14, p = 1.2e-3, −0.14, p = 8e-3). This was further seen in the nervous system-associated pheochromocytoma and paraganglioma tumors (PCPG, −0.06, p = 1.2e-3, −0.11, p = 0.12), for both ETG and ASE methods.

Next, we hypothesized that different cell types within the nervous system might have different NMD efficiencies. To determine if the strong reduction in iNMDeff observed in normal and cancerous brain tissues is specifically linked to neuronal cells or to glial cell types, we utilized cellular composition deconvolutions from GTex tissues’ bulk RNA-seq data, as estimated by Donovan M. K. R. *et al.*^55^. Analyzing the neuron cell type proportions in each brain tissue type, ranked by the median ETG iNMDeff (Fig. 3F), revealed that tissues with lower ETG iNMDeff had a higher proportion of neurons. Notably, the two brain samples with the lowest iNMDeff, the cerebellum regions BRNCHA and BRNCHB, had the highest neuron proportions (medians of 65% and 77%, respectively). We correlated iNMDeff with neuron proportions within each brain sample (Fig. 3F), finding that most correlations between neuron proportion and ETG iNMDeff across subregions were negative (R = −0.34).

In TCGA tumor samples, we performed a deconvolution using the *UCDBase* method^56^ and analyzed glioblastoma (GBM) and LGG tumor samples (see Methods). We stratified these cancer types into subtypes based on cell of origin or mutation type information^57^ (Fig. 3E). The cancer subtype with the highest estimated neuron cell type proportion (median of 16%), LGG_Codel (characterized by the co-deletion of chr 1p/19q), exhibited the lowest iNMDeff (median ETG −0.64), while at the other end, GBM_LGm6 (characterized by the DNA methylation cluster LGm6) had a median neuron proportion of 2% and a median ETG iNMDeff of 0.21. The ETG iNMDeff correlations with estimated fractions of neurons across these cancer subtypes were negative (Fig. 3E; median R of −0.19 in ETG), supporting that neural cells, rather than glia or other cells infiltrating these tumor samples, are the cell types with lowered NMD activity (Supp. Fig. S6A-B). This provides an explanation why between the two major brain cancer types in TCGA, the LGG has lower iNMDeff compared to the GBM (median neuron proportion of 10% vs 3%, and median ETG iNMDeff of −0.52 vs 0.04).

In sum, our analysis underscores the significant variability of NMD efficiency across tissues, with high iNMDeff in digestive tract tissues, and low iNMDeff in the reproductive and nervous system tissues, the latter being associated with neuron content of the tissue.

#### Extensive inter-individual variation of NMD efficiency

Having established there exist significant differences in NMD efficiency between human tissues, we next asked whether NMD efficiency is variable between different individuals, compared to a baseline variation between different variants within the same individual. For this, we derived the measure of NMD efficiency for a set of ASE PTCs (termed pNMDeff for PTC NMD efficiency, as detailed in Methods) in the TCGA separately for somatic and germline variants, and additionally for GTex germline variants. Next, we compared the intra-individual variation (different PTC in the same individual) with a randomized baseline, and then the inter-individual variation (same PTC across individuals) with a control randomization (Supp Fig. S6C).

The randomization controls display higher variances than either inter-individual pNMDeff (excess variance = 0.83 - 1.01 across the three datasets, p = 2.6e-7 - 1.3e-83) or intra-individual pNMDeff (excess variance = 0.41 - 0.78, p = 1.2e-1 - 1.7e-36). The two tests for non-random variation suggest that each PTC has some inherent property determining its pNMDeff, and that this property can be shared across different PTCs within an individual but different between individuals. The former is not unexpected given the various known rules of NMD efficiency depending on PTC placement^1,8^, however we note that here we selected those PTCs that should trigger NMD by the known rules, yet there is still systematic variation, therefore suggesting that additional NMD rules exist. The latter means that the PTCs within an individual are more consistent (less variability) in pNMDeff than expected at chance, suggesting mechanisms governing individual-level NMD efficiency. The reduced variance over random is considerable in both tests, suggesting that there is considerable between-PTC as well as, more interestingly for the matter at hand, inter-individual variation in NMD efficiency.

As an additional statistic on this data, when examining the Pearson correlations of pNMDeff between pairs of PTCs, the different PTCs across individuals had a notably higher correlation than randomized controls, and similarly so for the same PTC allele within individuals (p <= 2.2e-16 for both, Supp. Fig. S6D). It bears mentioning that, despite the systematic variability in NMD efficiency both across PTCs and across individuals, much of the NMD efficiency across different contexts is preserved, compatible with that NMD pathway is a conserved RNA surveillance mechanism. These findings motivated us to explore mechanisms that might underpin the inter-individual variability in NMD efficiency.

### Somatic pan-cancer CNA signatures are associated with NMD efficiency

#### Association analysis of somatic mutations

We hypothesized that genetic variants might generate the inter-individual variation in NMD efficiency. We initiated our investigation by considering TCGA tumors, and testing for genetic associations between our iNMDeff estimates and somatic mutations. We adjusted for various covariates, and carried out the analysis for each gene and each cancer type individually, including additionally a pan-cancer analysis (as detailed in Methods). Our focus was on: i) a set of 727 recognized cancer driver genes from the Cancer Gene Census (CGC)^58^; ii) 112 genes related to the NMD pathway sourced from two experimental studies^59,60^ and additional literature^3,7,61^ (see Methods); iii) the remaining 18,780 genes were considered as a baseline (“random genes”) for our analysis. We then tested these genes against different sets of somatic variants: firstly the point mutations -- synonymous, truncating (which includes nonsense, indels, and splicing variants), missense -- and secondly the CNAs.

For both analysis, in total we systematically conducted 6,493,740 tests across 19,619 genes and 33 cancer types (here, not stratified by subtype, to prevent increasing number of tests and reducing sample sizes), where each test utilized one NMD method (ASE or ETG) for discovery and the other for validation. To ensure the robustness of our association results, we calculated the lambda (inflation factor) for each cancer and type of somatic variant, excluding the few tests with lambda >1.5 (Supp. Fig. S7A-B). All significant hits using one NMD estimation method in one particular cancer type were re-tested in the same cancer type with the other NMD method, adjusting by FDR within those hits. With this criteria, two replicated significant associations emerged at a FDR threshold of 5%: a missense variant in the *TLX1* gene in LUAD and a truncating variant in the *CDH1* gene in BRCA, both associated with iNMDeff with opposite directions (Supp. Fig. S7C-D). Further exploration of the effect sizes of these two associations in other cancer types revealed that, for the *CDH1* association (mean ASE-ETG effect size = - 0.15), the directionality was consistent, but not significant, in pan-cancer (effect size = −0.1) and UCEC (effect size = −0.09), both being negatively associated with iNMDeff (Supp. Fig. S7E). For the *TLX1* association (effect size = 1.1), a consistent positive direction was observed in LUSC (0.34), Bladder urothelial carcinoma, BLCA (0.23), and Thyroid carcinoma, THCA (0.1), with a modest agreement also in pan-cancer (0.03), with associations in the opposite direction for other tumor types (Supp. Fig. S7E). This supports the association of somatic mutations in the *TLX1* transcription factor with reduced NMD efficiency.

#### Association analysis of somatic pan-cancer CNA signatures

We next proceeded to test associations of iNMDeff with somatic CNAs. Given that CNA often manifests as broader alterations affecting many genes up to an entire chromosome arm, we devised a custom method for somatic CNA association analysis. Instead of testing the gene-level CNA associations, which displayed inflation in our Q-Q plots (Supp. Fig. S8A-B), possibly because of extensive genetic linkage between CNA of neighboring genes, our method considers recurrent large-scale, multi-gene CNAs as monolithic units.

To do so, we performed a sparse principal component analysis (sparse-PCA) on the pan-cancer gene-level CNA data (see Methods, Supp. Fig. S9A), identifying 86 principal components, termed CNA principal component signatures (CNA-PCs). Notably, the first (higher variance explained) 47 CNA-PCs represented large-scale alterations, often spanning approximately arm-level changes, while the subsequent CNA-PCs pinpointed more localized events (Supp. Fig. S9B). Thereafter, we tested associations between the individual’s weights of these CNA-PC signatures and our iNMDeff, at the pan-cancer and cancer type level (see Methods). Utilizing ASE method for discovery and ETG for validation, we identified 3 pan-cancer CNA-PCs significantly associated with iNMDeff at FDR < 10%, replicating in both methods with the same direction of NMD effect (Fig. 4A): CNA-PC 3 (ASE p = 1.3e-2; ETG FDR = 1.2e-37), CNA-PC 52 (2.2e-2; 3.1e-9), and CNA-PC 86 (8.7e-2; 8.6e-37). Encouragingly, these recurrent CNA signatures exhibited consistent directionality in their associations across various cancer types (Supp. Fig. S10A). Of note, the sparse CNA-PCs 3 and 86 exhibited similar genome-wide patterns of CNA gains, with a Pearson’s correlation coefficient of 0.79.

**Fig. 4.**
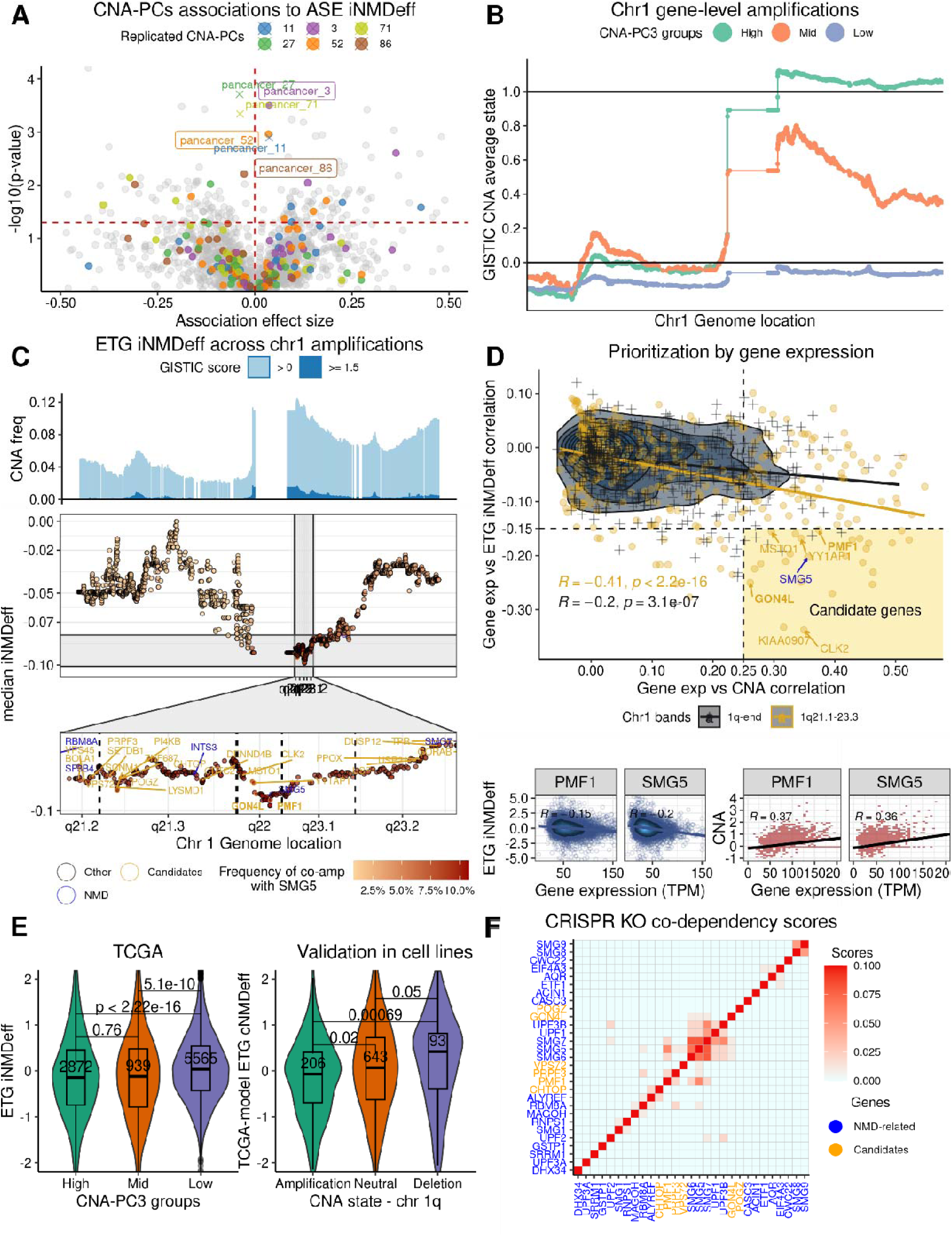
Somatic pan-cancer CNA signatures are associated with NMD efficiency. **A,** Pan-cancer CNA principal component (CNA-PCs) associations with ASE iNMDeff. Effect sizes are beta coefficients from linear models. The vertical dashed red line indicates a coefficient of zero, and the horizontal dashed red line marks the significance threshold at 10% FDR. Six significant pan-cancer CNA-PCs are color-highlighted, with three and replicated CNA-PCs (in ASE and ETG methods) PC3, 52, and 86 are marked with dots. **B**, CNA-PC3 reflects gene amplifications across chromosome 1, ordered by genome location, plotted along the X-axis, with amplifications assessed by averaging GISTIC CNA scores for each gene across TCGA participants (Y-axis). Samples are categorized into “High”, “Mid” and “Low” bins based on scores from pan-cancer CNA-PC3. **C**, Gene-level iNMDeff scores are calculated as the median iNMDeff across individuals with focal CNAs for each gene on chromosome 1, arranged by genomic location (X-axis). The top panel shows pan-cancer CNA frequencies per gene in TCGA. The middle panel displays gene-wise iNMDeff scores, with red intensity indicating co-amplification frequency with NMD factor gene SMG5. The bottom panel zooms into the 1q21.1-23.1 region, highlighting a reduction in iNMDeff when amplified. “Candidate” causal genes, “NMD” factors or related genes, and the rest of the genes (“Other”) are color-highlighted. **D**, Candidate gene prioritization from chromosome 1q using two scoring criteria: Correlation between gene expression and ETG iNMDeff (Y-axis) or CNA amplification (X-axis). Genes in the 1q21.1-23.1 region are in orange, NMD-related genes are in blue, remaining genes are in black. Candidates are expected in the bottom-right quadrant, indicating correlation between expression and iNMDeff (negative) and between expression and CNA amplification (positive). Vertical and horizontal dashed black lines denote the thresholds for selecting candidates. Example candidate gene PMF1 and known NMD-factor SMG5 are displayed in the bottom panels, showing gene expression vs. ETG iNMDeff (bottom left) or CNA amplification (bottom right). **E**, Stratification of ETG iNMDeff by CNA-PC3. The left panel presents a scaled ETG iNMDeff (Y-axis) of individuals classified by CNA-PC3 groups as “High”, “Mid”, and “Low”. The right panel provides a validation in cell lines from the Achilles project, categorized by 1q CNA status: gain, neutral, and deletion; NMD was assessed with the proxy model for ETG cNMDeff. P-values are obtained through a two-sided Mann-Whitney U test. **F**, Clustering heatmap of CRISPR co-dependency scores for 30 candidate genes and 23 NMD-related genes from chromosome 1q. The analysis focuses on the 1q21.1-23.1 region, revealing a distinct cluster of candidate genes alongside NMD factor genes. Co-dependency scores have been normalized using the RPCO onion method, where generally, higher scores denote stronger gene functional associations.

#### Somatic chromosome 1q gain associates with reduced NMD efficiency

To further examine these three recurrent CNA signatures, we categorized individuals based on their scores for each CNA-PC, ranging from lower to higher. For CNA-PC3, individuals in group “High” predominantly exhibited an arm-level gain of chromosome 1q (Fig. 4B). In contrast, group “Mid”, with intermediate scores, displayed a more localized gain around the 1q21-23.1 region. The group “Low”, with the lowest or ∼ 0 scores in the sparse PCA, lacked notable CNA alterations in 1q. Interestingly, CNA-PC86 mirrored the pattern observed in CNA-PC3 but with a more narrow focal gain on 1q21-23.1 (Supp. Fig. S11A), thus both recurrent CNA signatures reflect chr 1q gains, albeit at different scales. The individuals in groups “High” and “Mid” who exhibit chr 1q gains, demonstrate a decrease in both ASE and ETG-estimated iNMDeff (Fig. 4E and Supp. Fig. S11C-D) compared to those in group “Low”, who lack any notable CNA alterations (median ETG iNMDeff = 2.07 vs 2.12, two-sided Mann-Whitney *U* test p = 5.1e-10, comparing “High” against “Low”; median ASE iNMDeff = 0.49 vs 0.53, two-sided Mann-Whitney *U* test p = 4.7e-2, comparing “High” against “Low”).

To validate this 1q somatic CNA gain association with NMD efficiency in an additional data set, the GTex is not suitable since it does not contain somatic alterations. Instead, we used the transcriptomic and CNA data of 1450 human cell lines from the Cancer Cell Line Encyclopedia (CCLE)^62^. We built proxy models of our ETG iNMDeff (see Methods) using global gene-level expression data to predict the NMD efficiency for every cell line (cNMDeff). The models were trained in either TCGA or GTex cohorts (cross validation R² = 0.4 and 0.82 for TCGA and GTex, respectively, Supp. Fig. S12A). We replicated our earlier findings (Fig. 4E and Supp. Fig. S12B); cell lines with 1q gains exhibited a decreased cNMDeff in comparison to diploid cells (p = 0.02 and non-significant, for TCGA-based and GTex-based models of cNMDeff, respectively). This reduction was even more pronounced when comparing cells with 1q gains to those with 1q deletions (p = 6.9e-4 and p = 0.02, for TCGA and GTex models respectively).

We next asked which cancer types were most influenced by the iNMDeff changes linked with chromosome 1q gain. To study this, we analyzed the distribution of iNMDeff in individuals across the Low/Mid/High groups, for the three identified CNA-PC signatures, considering each cancer type separately (Supp. Fig. S13A). BRCA, liver hepatocellular carcinoma (LIHC), cholangiocarcinoma (CHOL), and LUAD had over 50% of their samples falling into group “High” for CNA-PC3, indicating that 1q gains are common in breast, liver and lung cancers. This is also reflected in iNMDeff. For instance, for 61.8% of LUAD samples with 1q gain, 67% of those also had a lower ETG iNMDeff than the median ETG iNMDeff across the LUAD samples with no gain. These 1q gains were proposed to be selected because they increase dosage of the *MDM4* oncogene^63^, whose product downregulates the p53 protein post-translationally^64^, hence phenocopying the loss of the tumor suppressor *TP53*^65^. However, there are other oncogenes in chr 1q that might be driving this gain.

When examining the more localized 1q21-23.1 amplification represented by CNA-PC86, 10-13% of the tumors from BRCA, LUAD and LIHC were found in Group “High” (Supp. Fig. S13B), and associated with reduced iNMDeff. Furthermore, other cancers, including uterine carcinosarcoma (UCS), LUSC, OV, and skin cutaneous melanoma (SKCM), also exhibited high incidences of both these recurrent 1q gain CNA-PC3 and PC86 (Supp. Fig. S13A-B). We note this latter segment does not include the *MDM4* oncogene, which is located at 1q32.1, thus other genes must be driving this recurrent change in CNA-PC86, with possible candidates being known cancer genes *BCL9* (1q21.2), *ARNT* (1q21.3), *SETDB1* (1q21.3) or *SF3B4* (1q21.2).

#### Identifying NMD-associated genes via focal CNA analysis

While CNA gains can affect numerous dosage-sensitive genes in one event, only a subset or a single gene within the CNA segment might be directly implicated in the observed phenotype, here the NMD deficiency. Within the 1q arm reside three known NMD factors and two NMD-related genes identified in experimental and computational studies^59,60^: *RBM8A* (located at q21.1), a core component of the EJC; *SMG5* (q22) and *SMG7* (q25.3), both core NMD factors; *SF3B4* (q21.2), a known splicing factor and oncogene; and *INTS3* (q21.3), involved in snRNA transcription. To prioritize candidate genes responsible for the NMD deficiency associated with 1q gains, we computed the median iNMDeff of individuals having a focal copy number amplification for each gene separately; considering only focal CNA events for this analysis helps distinguish between the linked genes. Mapping these gene-wise scores along chr 1, we observed that tumor samples with CNA gains in genes proximal to the 1q21-23.1 region (Fig. 4C, top panel) have a pronounced reduction in ETG iNMDeff (Fig. 4C, bottom panel). A similar local trend was evident in the ASE iNMDeff scores (Supp. Fig. S14A). Notably, the NMD-related genes mentioned above, except for *SMG7*, are situated within this region.

To narrow down potential candidate NMD-modulating genes within this gene-dense region, we employed two scoring criteria. First, we calculated the correlation between gene expression -- presumably the downstream effect of the CNA relevant to the phenotype in question -- and ETG iNMDeff. Second, as a supporting test for prioritization, we assessed the correlation between gene expression and CNA. The correlation between these two scores (Fig. 4D and Supp. Fig. S15A for ASE) is stronger within our target region (R = −0.41) than outside (R = −0.2), supporting that this segment harbors causal genes. We established thresholds for selecting candidates based on the two criteria (Fig. 4D), and among the 30 selected candidate genes, *SMG5* was the only known NMD core gene. Parsimoniously, the *SMG5* gain might generate the effect on NMD efficiency, however we considered the hypothesis that other genes within this segment might affect NMD efficiency.

To pinpoint the potential causal gene from the 30 identified candidates, we considered genetic interactions inferred from gene-level CRISPR screening data obtained from the Achilles Project^66^, allowing to infer functional links between genes by studies of codependency profiles across cell lines. Here, we used the de-biased data (*onion* method, RPCO), suggested for its power for inferring gene function^67^ (see Methods). Our hypothesis posits that if these candidate genes 1q21-23.1 are indeed implicated in NMD efficiency, their CRISPR knockout fitness effect should correlate with established NMD factor genes like *UPF1*, *UPF2*, *UPF3B*, *SMG1*, and others; the codependency implies a functional link.

Therefore, we clustered and visualized these codependency CRISPR scores involving the 30 candidate genes, 23 NMD-related genes, and 18 random negative control genes on chr 1q but outside the 1q21.1-23.1 region (see Methods). This revealed a prominent cluster of functional interactions (Fig. 4F) comprising NMD factor genes such as *SMG5, SMG6, SMG7, UPF1*, and *UPF3B*, suggesting that the RPCO scores derived from CRISPR screens^67^ are powerful for identifying NMD factors. Within this cluster were two candidate genes residing within 1q21.1-23.1, *PMF1* and *GON4L*, showing notable codependency scores with *SMG5* (0.06 and 0.0062, for *PMF1* and *GON4L*, respectively)*, SMG6* (0.02 and 0.006), and *SMG7* (0.019 and 0.012). For comparison, the codependency score between *SMG5* and *SMG7* is 0.077, and the majority of scores with control genes are 0. Furthermore, *VPS72* and *PRPF3* exhibited codependency with *SMG5* (0.08 and 0.011, respectively), and so might have relevance. In a more detailed quantitative analysis, we examined all 274 genes within 1q21.1-23.1. For each, we computed the mean CRISPR codependency scores with 10 core NMD factor genes and compared these with the mean CRISPR score for the same gene with the previous 18 random negative control genes on 1q. We identified 14 genes with FDR < 5% (one-sided Mann-Whitney *U* test). Notably and as expected, 3 of these genes are directly involved in NMD (*SMG5*, *RBM8A*, and *SF3B4*) (Supp. Fig. S15B). The 4 genes identified above by a visual inspection of the clustering – *PRPF3* (FDR = 4.6%, one-side Mann-Whitney *U* test), *GON4L* (4.6%), *VPS72* (3.1%), and *PMF1* (2.5%) – also emerged as significant in this analysis. Additionally, 7 genes not considered as candidates due to not meeting our initial threshold criteria (correlations with gene expression) were significant here. Interestingly, 10 of these 14 genes are related to a RNA metabolic process (see Discussion).

Our analyses underscore that the 1q21.1-23.1 genomic region harbors multiple genes with potential to modulate NMD activity upon CNA gains and overexpression, some of which have not been previously identified as NMD factors, with prime candidates being *PMF1* and *GON4L*.

#### Somatic chromosome 2q gain also associates with reduced NMD efficiency

Next, we analyzed the CNA-PC52 signature, negatively associated with iNMDeff. Individuals with higher scores in the CNA-PC52 signature exhibited CNA gain peaks at chromosome 2p and 2q, (Supp. Fig. S11E), however, of relatively lower intensity (average GISTIC score around 0.2), suggesting this CNA-PC captures low-level gains. A decrease in iNMDeff was observed in individuals with the 2q gain peak spanning approximately the 2q31.1-2q36.3 region (Supp. Fig. S14B-C). This CNA signature was prevalent in testicular germ cell tumors (TGCT, 97% of samples), adrenocortical carcinoma (ACC, 69%), OV (39%), LUSC (38%), and kidney renal papillary cell carcinoma (KIRP, 37%) (Supp. Fig. S13C). Within this region, four NMD-related genes are located: *CWC22* (2q31.3), *SF3B1* (2q33.1), *NOP58* (2q33.1), and *FARSB* (2q36.1). Further investigation by our iNMDeff-gene expression and CNA-gene expression correlations (see above) identified 18 candidate genes, including *CWC22*, *SF3B1* and *NOP58* (Supp. Fig. S15C). Additionally, we analyzed the CRISPR KO codependency scores to assess whether any of the 303 genes within the region had genetic interactions to 10 well-known NMD factor genes, compared to 383 control genes outside the region (Supp. Fig. S15D-E). In this CRISPR analysis, *NOP58* (one of the NMD-related genes, FDR = 7.2%), *BARD1* (9.4%), *SF3B1* (FDR = 16%), *CWC22* (12%), and *FARSB* (16%) were significant at FDR < 25% (one-sided Mann-Whitney *U* test), plus 8 additional genes not known to be NMD-associated (Supp. Fig. S15D), two of them overlapping with our set of 18 candidates (*PRKRA* and *CTDSP1*). Overall, the somatic CNA gains in the 2q31.1-2q36.3 region may also alter the expression of various RNA processing genes, associated with alterations in NMD efficiency.

### Rare deleterious germline variants are associated with NMD efficiency

#### Rare variant association analysis identifies 10 genes associated with NMD efficiency

We hypothesized that an additional cause for inter-individual variation in NMD efficiency could be germline genetic variation across individuals, affecting NMD-relevant genes. To test this, we conducted a gene-based rare variant association study (RVAS, see schematic from Fig. 5A), relying on the state-of-the-art SKAT-O test^68^ (see Methods).

**Fig. 5.**
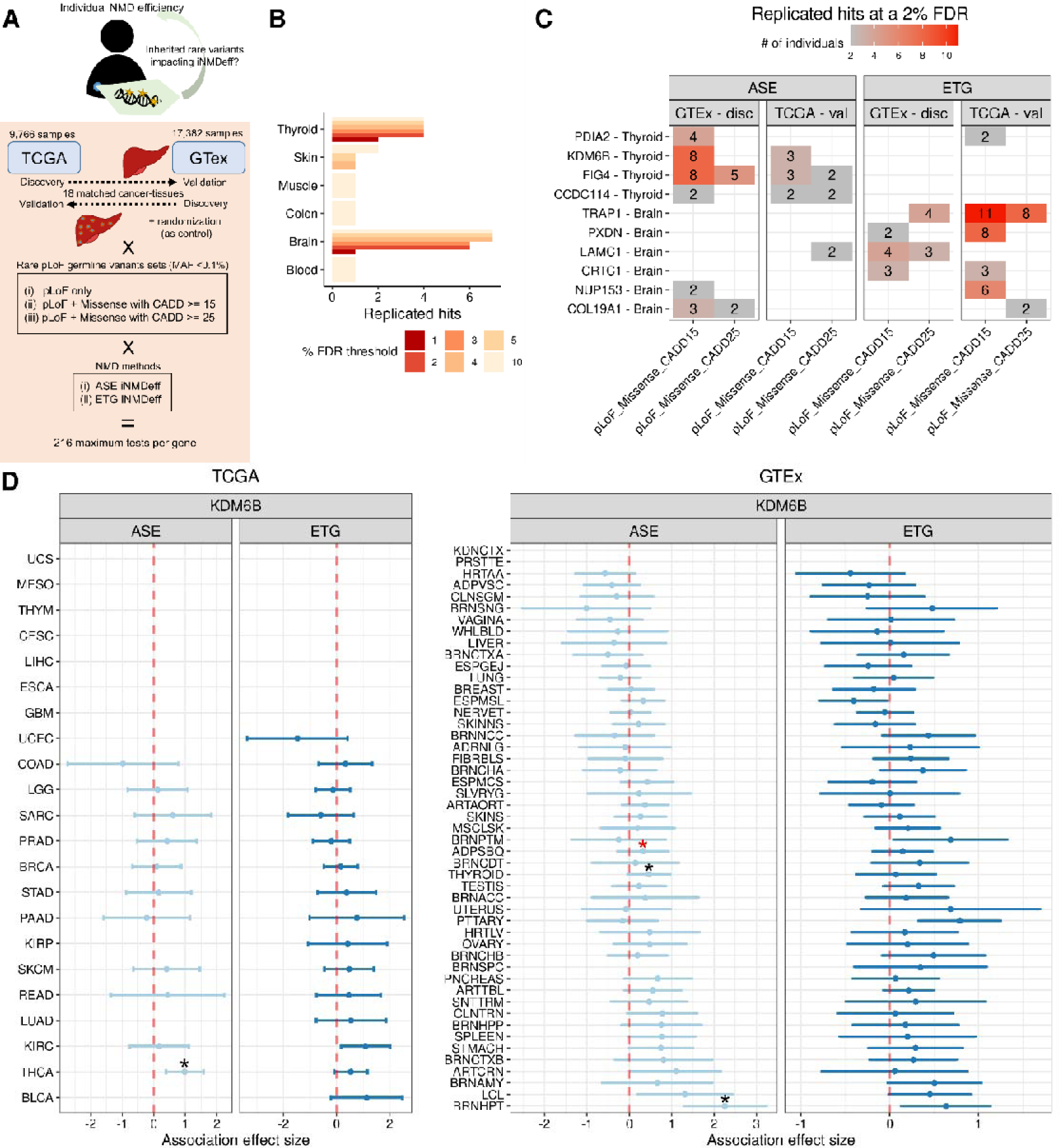
Rare deleterious germline variants are associated with NMD efficiency. **A**, Overview of RVAS analysis. Illustrates the SKAT-O analysis framework for identifying associations between rare germline putative loss-of-function (pLoF) variants and NMD efficiency. Utilizes either TCGA or GTex as discovery cohorts with the other serving as validation, and vice versa. The analysis spans 18 matched cancer-normal tissues, three sets of pLoF variants, and employs two distinct NMD methods for gene-level associations. iNMDeff values were randomized as a control measure. **B**, Replication of gene-tissue associations. Depicts the number of replicated gene-tissue pairs across various tissues (Y-axis) against the range of FDR thresholds from 1% to 10% (X-axis). **C**, Detailed view of replicated associations at 2% FDR. Shows replicated gene-tissue pairs (Y-axis) at a 2% FDR threshold, categorized by pLoF variant set (X-axis), NMD method, and the cohort where the association was identified (either discovery or validation). The values indicate the total count of individuals harboring a rare pLoF variant within the successfully replicated genes. **D**, Gene burden test for KDM6B associations. Analyzes the association between rare pLoFs within the KDM6B gene and iNMDeff, represented by the effect size (beta coefficient) from linear model associations (X-axis) against cancer types (TCGA, left panel) or normal tissues (GTex, right panel), segmented by NMD method. Displays 95% confidence intervals to indicate the precision of the effect size estimates. An asterisk signifies FDR < 5%, highlighting statistically significant associations. Tissues with no values did not have samples with rare pLoFs in the gene.

We defined three stringency thresholds for determining putative loss-of-function (pLoF) for the rare germline variants i.e. those with a MAF < 0.1%. One threshold involved only NMD-triggering PTC variants, and the other two additionally included the predicted deleterious missense variants using CADD scores^69^ at two different cutoffs: >= 25 (more stringent) and >= 15 (permissive).

We tested all genes in which at least 2 individuals carried a rare pLoF variant, for the three pLoF stringency levels, using the two iNMDeff methods, ASE and ETG, and correcting by potential confounder variables (see Methods). Testing was performed in each cohort, TCGA and GTex, contributing cancer and normal tissue, respectively. Significant associations identified in one cohort (TCGA or GTex) were subsequently re-tested requiring replication in the other cohort (GTex or TCGA, respectively), relying on pairing 18 TCGA tumor types to corresponding GTex normal tissues (Supp. Table S2).

A total of 850,721 tests to associate iNMDeff with rare germline variants were conducted in GTex and 221,613 in TCGA, after excluding those dataset-tissue pairs with a lambda >= 1.5. The number of tests in GTex was higher due to multiple tissue samples analyzed per individual, allowing separate testing of the same gene across various tissues. Across the three variant sets, two NMD methods, and two cohorts, considering all directions of validation, we identified 48 tissue-specific significant associations that were successfully replicated, at a nominal FDR threshold of 2%. This is in contrast to the 24 significant associations found by randomizing the iNMDeff estimates (Supp. Fig. S16A). Out of these replicated associations, 27 were discovered in GTex and validated in TCGA. These hits span 10 unique individual gene-tissue type pairs clustered in 2 matched-tissues: thyroid (4) and brain (6) (Fig. 5B-C). Expanding the nominal FDR threshold to a more permissive 10% revealed additional replicated genes across four more tissues: blood (1), colon (1), muscle (1), and skin (2), suggesting that future analyses with larger sample sizes may identify additional NMD-modulating genes, and that NMD-modulating variants may manifest in various tissues.

We further calibrated these FDR estimates by comparing against a distribution of FDRs obtained from running SKAT-O on a randomization of the iNMDeff estimates. This resulted in stringent estimates of empirical FDR (Supp. Fig. S16A), where the nominal FDR of 2% corresponded to a conservatively estimated empirical FDR threshold of approximately 30%, calculated as 3 replicated hits from randomized iNMDeff data, versus 10 hits obtained from our observed iNMDeff values. The 10 gene-tissue pair associations that met our criteria are detailed in Figure 5C, categorized by pLoF stringency variant set and NMD estimation method. The same table for an FDR of 5% is also available (Supp. Fig. S16B).

To assess the effect size of the associations for 10 replicated genes and determine their direction of effect on iNMDeff, we utilized a gene-burden test through general linear regression (see Methods) to complement the SKAT-O results. We conducted this estimation separately for each NMD method in both TCGA and GTex, across all cancers or tissues, respectively (Supp. Fig. S17A-B). Among the genes studied, the chromatin modifier *KDM6B* stood out in that it demonstrated a consistently significant positive effect size in the thyroid (0.46, SKAT-O FDR = 2%, in ASE), our GTex-discovery tissue, and in THCA (0.98, SKAT-O FDR = 0.5%, in ASE), our TCGA-validation matched cancer type. The effect sizes in the ETG method were consistent with the ASE method (0.07 in GTex thyroid, 0.98 in TCGA THCA). The coefficient direction is positive, indicating that individuals with a pLoF variant in this gene have a higher iNMDeff than those without (Supp. Fig. S18). Moreover, a positive effect size is also observed in the hypothalamus brain region, (BRNHPT in GTex: 2.25 in ASE; 0.63 in ETG) and in the adipose subcutaneous tissue (ADPSBQ: 0.32 in ASE; 0.14 in ETG), with the ASE SKAT-O FDR = 0.55% and 17%, for brain and adipose respectively. The other tissues and cancer types, while not significant with SKAT-O, usually exhibited a similar positive trend (Supp. Fig. S17), suggesting this *KDM6B* association with NMD efficiency may not be a thyroid-specific association but likely impacts other tissues as well.

From the 6 brain-identified replicated genes (Fig. 5C), the consistency of effect sizes (pLoF burden) across the two cohorts (GTex tissues and TCGA tumors) was observed only for *LAMC1*, (Supp. Fig. S17 and S18). For *NUP153*, consistent effects were observed across various brain tissue samples within GTex but did not extend to TCGA brain tumors, possibly due to normal-cancer tissue differences and/or due to that glioma brain cancers originate from glia rather than neurons (Supp. Fig. S17 and S18). Overall, our analysis provides evidence that deleterious germline variants in diverse genes may alter NMD efficiency in individuals, with the histone demethylase *KDM6B* being a prime candidate for follow-up.

### NMD efficiency modulates selection of somatic nonsense mutations

Previous research^11,45^ reported that somatic nonsense mutations (producing PTCs) are positively selected specifically in NMD-triggering regions of tumor suppressor genes (TSGs) and they may be negatively selected in oncogenes (OGs). This led us to ask whether the “global” NMD efficiency of an individual, in addition to the NMD efficiency of specific PTCs in the individual’s genome, provides context for modeling the selection upon somatic PTCs. In specific, we tested whether there is stronger positive selection for NMD-triggering PTCs among individuals/tumors with high NMD efficiency, compared to individuals/tumors with low NMD efficiency, plausible due to the tumors’ ability to utilize NMD for ablating the activity of TSGs. Serving as a negative control, the NMD-evading PTCs would not be anticipated to show a notable difference in positive selection between individuals with higher versus lower iNMDeff. Regarding oncogenes, we do not expect positive selection on somatic PTC except in rare exceptions, while there might be modest negative selection instead ^11,70^.

To test our hypothesis, we employed the standard dNdS tool^71^ that compares observed mutation counts in a gene to an expectation modeled from covariates (e.g. gene expression) and from synonymous mutation counts. Here we focussed on nonsense somatic mutations (splitting by NMD-triggering and NMD-evading PTCs) and missense somatic mutations (serving as negative controls since they should not directly trigger NMD), within the whole TCGA cohort of tumor exomes (see Methods). A dN/dS ratio above 1 for a gene indicates positive selection, while a ratio below 1 signifies negative selection. The analysis involved stratifying patients based on their iNMDeff into high and low groups and comparing dN/dS, i.e. estimating conditional selection on the cancer genes depending on iNMDeff. Moreover, we studied 50 TSGs and 52 OGs (see Methods). Briefly, we started with the 727 cancer genes from the CGC list^58^, and intersected them with the genes with experimental evidence they act as TSGs or OGs (“STOP” and “GO” genes^72^), and with the significant genes (q-value < 0.01) from the Mutpanning set^73^. This led to a core set of 102 higher-confidence cancer genes.

For TSGs, we first checked missense mutations, which normally do not trigger NMD. There was no significant difference in selection between high and low iNMDeff individual groups, defined either by the ETG method (Fig. 6A, right panel) or the ASE method (Supp. Fig. S19A, right panel) for iNMDeff (missense mutations located in NMD-triggering regions ETG dNdS = 1.19 vs 1.20, p = 0.53, and NMD-evading regions dNdS= 1.03 vs 1.10, p = 0.61, one-sided Wilcoxon paired tests). However, in the case of nonsense (PTCs) mutations, we observed a difference with contrasting directions: genes with NMD-triggering PTC variants exhibited a trend towards higher positive selection in tumor samples with high ETG iNMDeff, whereas tumors with low ETG iNMDeff demonstrated no positive selection (dNdS = 1.64 vs 1.33, p = 0.12), thereby suggesting that individual-level, global NMD efficiency can indeed shape selection on cancer genes (Fig. 6A and Supp. Fig. S19A, left panels). Further, when examining the specific genes driving this higher positive selection in TSGs (Fig. 6B and Supp. Fig. S19B), we identified the top genes with dNdS differences (> 1) between high and low iNMDeff groups: *SMAD4* (8.4), *TP53* (8.36), *APC* (3.10), *XPA* (2.46), *NRG1* (1.25), *FAT1* (1.09), and *EP300* (1.06); for these genes, there is evidence that global NMD efficiency boosts positive selection. In contrast, genes with the most negative dNdS differences (< −1) were *ARID1A* (−1.29), *BCOR* (−1.89), *RB1* (−2.53), *CDKN1B* (−3.41), *PTEN* (−4.56), and *SMAD2* (−6.19), and in those genes there is not evidence that iNMDeff affects positive selection. In the case of OGs (Supp. Fig. S19C-D), for nonsense mutations, NMD-triggering PTCs trended towards negative selection overall, but no significant differences were noted between high and low ETG iNMDeff groups (dNdS = 0.77 vs 0.66, p = 0.42). In a cancer-type specific analysis, no significant differences were found after FDR adjustment. Nonetheless, trends of higher positive selection on TSG in the ETG high iNMDeff samples (p < 0.25) were seen in STAD_MSI (p = 7.6e-2), UCEC_POLE (p = 0.13), lung cancers LUAD (p = 0.17) and LUSC (p = 0.19), GBM (p = 0.17) and COAD_MSI (p = 0.22); many of these cancer types typically have high mutation burdens. In summary, our data suggests that higher individual-level NMD efficiency enables more effective positive selection of PTCs in TSG, including, prominently, the master tumor suppressor gene *TP53*.

**Fig 6.**
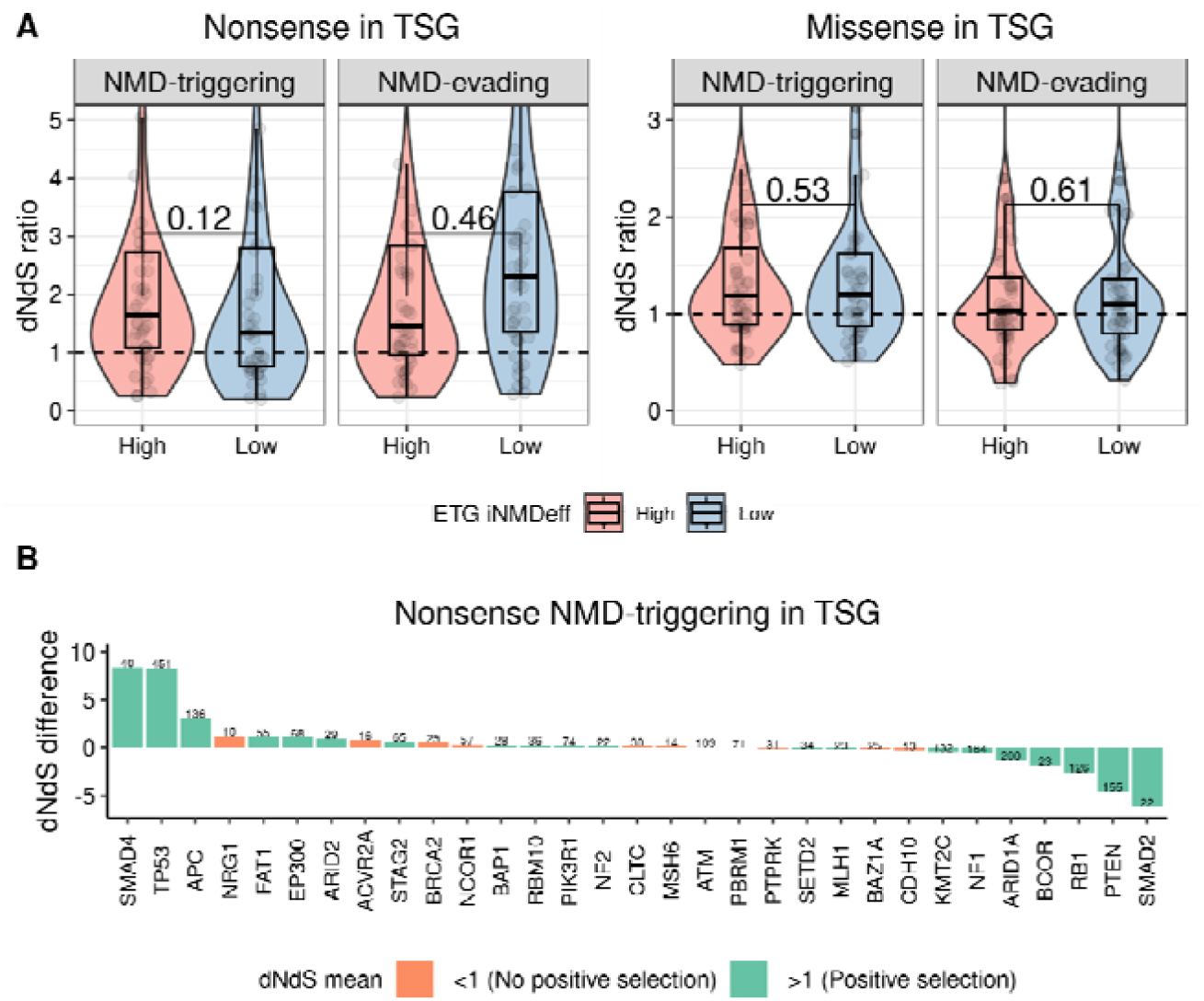
NMD efficiency modulates somatic selection in tumor suppressor genes. **A**, The dN/dS ratios (Y-axis) of tumor suppressor genes (TSGs) for both NMD-triggering and NMD-evading nonsense (left panel) and missense (right panel) somatic mutations are compared between two groups of individuals with high and low iNMDeff, as determined by the median ETG iNMDeff. Statistical significance is assessed using one-sided Wilcoxon tests on paired data. **B**, The differences in dN/dS (Y-axis) between high and low iNMDeff groups are plotted for each gene (X-axis), specifically for NMD-triggering nonsense mutations within TSGs. Positive values indicate a stronger selection pressure in the high iNMDeff group compared to the low group, and vice versa. The number above each bar denotes the total count of nonsense mutations contributing to the dN/dS calculation for both groups. Genes with less than 10 counts were removed. The barplot colors correspond to the mean dNdS ratios of the two iNMDeff groups (high and low): green indicates a mean ratio greater than 1, suggesting positive selection, while orange signifies a mean ratio less than 1, indicative of lack of positive selection.

### NMD efficiency impacts survival in chemotherapy and immunotherapy treated patients

#### NMD efficiency is associated with overall survival (OS) of cancer patients

In light of our previous findings that PTC-level NMD efficiency can modulate phenotype severity of genetic diseases^11^, we explored, here focusing on cancer, whether individual-level NMD efficiency influences cancer survival outcomes. Our analysis utilized Kaplan-Meier (KM) curves and Cox proportional hazards regression models with overall survival (OS) as our outcome, binarized into “High” and “Low” groups at varying thresholds, across 47 various TCGA cancer types stratified into subtypes, including pan-cancer, and employing randomized data as a negative control (see Methods).

After applying FDR correction to all 960 tests (480 FDR adjusted meta-p combining ETG and ASE estimations of NMD efficiency), we identified 17 significant tests at a nominal FDR of 5%, compared to 23 in the 9600 randomized tests, yielding an empirical FDR of 16% (results for individual percentiles given in Supp. Table S3). The significant results spanned four cancer types: SKCM (median FDR meta-p = 0.78%, median hazard ratio (HR) = 1.61), ESCA_ac (FDR = 3.2%, HR = 2.22), PRAD (FDR = 2.8%, HR = 0.11) and LUAD (FDR = 0.75%, HR = 3.49). Of note, in ESCA_ac and LUAD, we were unable to calculate the Cox model for ASE at due to missing data; thus, the reported meta-p reflects only the ETG method. In PRAD, higher iNMDeff correlates with increased survival (Supp. Fig. S20B and F). Roles of NMD in carcinogenesis are recognized to be two-fold, where NMD can boost some tumor-promoting but also some tumor-suppressive mechanisms^30,74^. We suggest that the balance tips towards the latter option for prostate cancer, where a more active NMD implies a less aggressive tumor. In contrast to the lowly mutated PRAD, in the often highly mutated SKCM (Supp. Fig. S20A and E), and also ESCA_ac (Supp. Fig. S20C and G) and LUAD (Supp. Fig. S20D and H), higher iNMDeff is associated with reduced survival. This finding aligns with proposed approaches to potentiate immune response to tumors by NMD modulation^1,46,75,76^, as a high number of SKCM samples in TCGA were from patients who received immunotherapy (73 out of 469). In contrast, only 5 LUAD samples in TCGA, and none from ESCA_ac and PRAD, were treated with immunotherapy. We hypothesize that patients with lower iNMDeff might experience better treatment outcomes, as higher NMD efficiency could result in the loss of truncated peptides (via PTC) that act as neoantigens, essential for the efficacy of immunotherapy.

#### NMD efficiency predicts progression-free survival (PFS) in treated patients

As another test of the impact of NMD efficiency on patient outcomes, we stratified patients based on the type of treatment and focused on progression-free survival (PFS), considering treatments separately and comparing the top 20% of individuals with the highest iNMDeff against the lowest 20%. For SKCM patients lacking available treatment data (Fig. 7A and Supp. Fig. S21A for ASE), KM curves showed no significant difference (log-rank test) in PFS between the high and low iNMDeff groups (ETG p = 0.73, ASE p = 0.77).

**Fig. 7.**
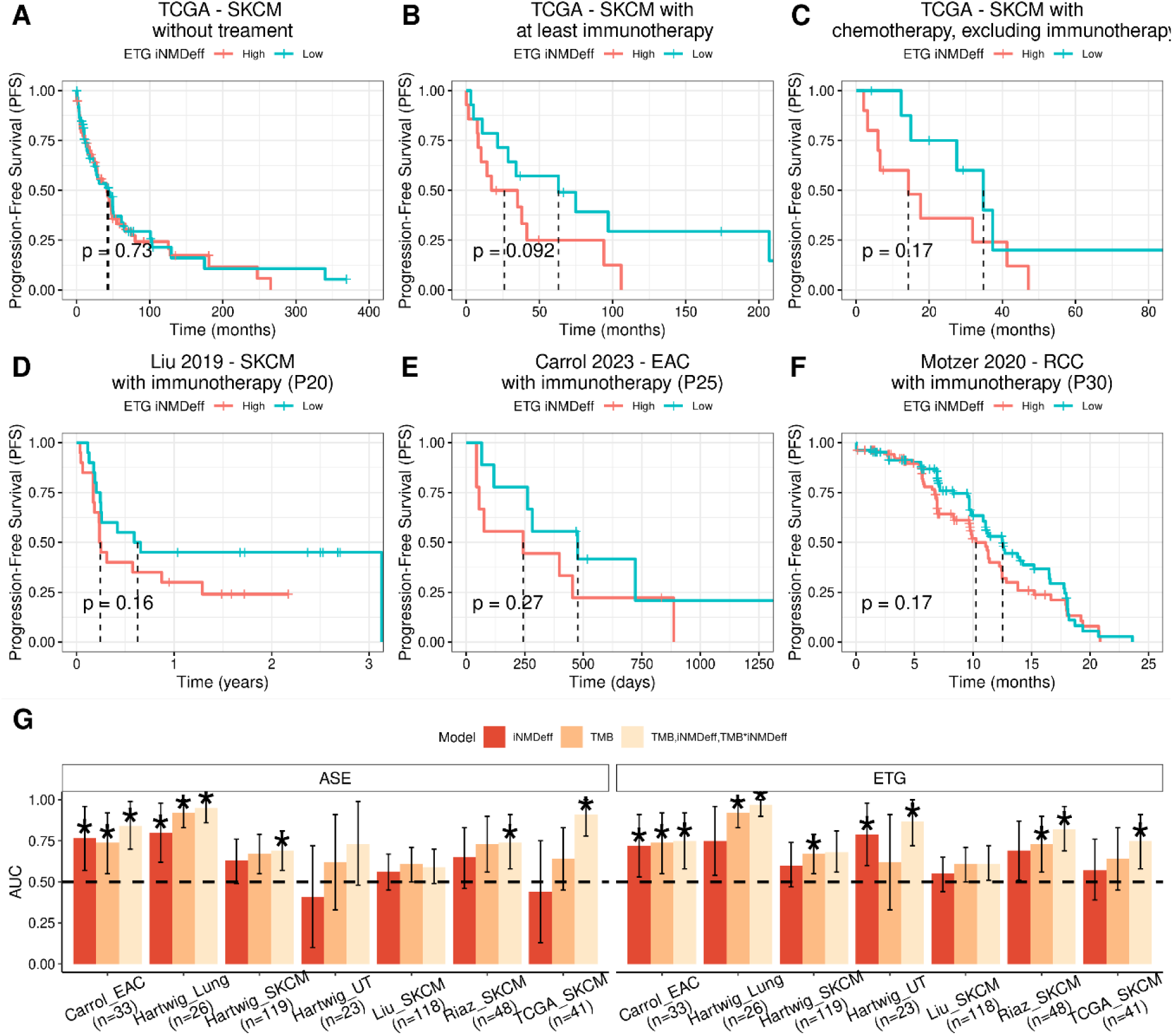
NMD efficiency impacts progression-free survival (PFS) and response to immunotherapy. **A-C,** Kaplan-Meier (KM) survival curves comparing progression-free survival (PFS) outcomes between groups with high versus low iNMDeff, as determined by the median of ETG iNMDeff in TCGA SKCM. The analysis is divided into three patient categories: untreated (A), those treated with immunotherapy at least once (B), and those treated exclusively with chemotherapy, excluding immunotherapy patients (C). Log-rank test p-values quantify the statistical significance of the survival differences. **D-F,** Validation of the KM curves for PFS in immunotherapy treated patient cohorts from Liu et al. 2019 (SKCM - skin cancer, melanoma, D), Carrol et al. 2023 (EAC - esophageal adenocarcinoma, E), and Motzer et al. 2020 (RCC - renal cell carcinoma, F). **G**, Performance of logistic regression models predicting immunotherapy response based on different variables: i) iNMDeff alone; ii) TMB alone; iii) a combination of iNMDeff, TMB, and their interaction term iNMDeff*TMB. iNMDeff was either from the ASE (left panel) or ETG (right panel) methods. Each model’s predictive power is indicated by the area under the curve (AUC) on the Y-axis, with the dataset and sample size (n) specified on the X-axis. A black dashed line at an AUC of 0.5 represents the threshold for random prediction accuracy and an asterisk denotes when the AUC is significantly different from it at a p-value < 0.05 via one-side DeLong test.

However, a difference emerged among patients who received immunotherapy (along with a maximum of one chemotherapy treatment; Fig. 7B and Supp. Fig. S21B for ASE), noting reduced PFS in the high iNMDeff group compared to the low iNMDeff group (ETG p = 9.2e-2 and ASE non-significant). The same trend was observed when discretizing at different thresholds (percentiles: 20 to 30, HR = 1.77 - 2.02, FDR = 32%). The borderline significance here might be attributed to the small number of SKCM patients treated with immunotherapy (n = 73). The TCGA pan-cancer analysis (includes all immunotherapy treated patients, n = 184) revealed a trend at the percentiles 20-25th (HR = 1.31 - 1.88, FDR = 19%) (Supp. Table S4).

Next, we asked if this effect of NMD efficiency on survival is particular to immunotherapy, or it extends also to other types of therapies. Among SKCM patients treated with chemotherapy, but not immunotherapy (Fig. 7C, Supp. Fig. S21C), a similar trend was observed (ETG p = 0.17 and ASE p = 0.16; ETG HR = 2.81, p = 5.1e-2). Thus, we cannot exclude that efficient NMD might be associated with poor response of melanoma not only to immunotherapy, but also to chemotherapies.

We next asked if other cancer types apart from SKCM show associations between iNMDeff and response to chemotherapy in terms of PFS. Again, from within the patients receiving chemotherapy, we compared the high iNMDeff against the low iNMDeff group. Indeed, we identified four additional cancer types following a similar trend at an FDR-adjusted meta-p < 10% (results for individual percentiles given in Supp. Table S5): pancreatic adenocarcinoma (PAAD, median HR = 2.16, median FDR = 3.8%), LGG (HR = 1.57, FDR = 4.5%), bladder cancer (BLCA_Basal_scc subtype, ETG: FDR = 3.4%, HR = 3.99) and ESCA_scc (ETG: FDR = 0.047%, HR = 2.5). No other cancer types were found with an opposite trend for the same FDR threshold. The positive HR, highlighting the worse PFS in chemotherapy-treated patients whose tumors have higher NMD efficiency extends to other cancer types. At a more relaxed FDR threshold of 25%, caution must be taken as we found additional cancer types, some with an opposite direction (better PFS, Supp. Table S6).

#### Association of iNMDeff with immunotherapy response in diverse cancer types

To validate the association of NMD efficiency with immunotherapy response (as PFS), we analyzed independent datasets that had both RNA-seq and WGS/WES available: i) Liu et al. 2019^77^, comprising 144 melanoma patients treated with anti-PD-1 immune checkpoint inhibitors (ICI); ii) Carroll et al. 2023^78^, involving 35 inoperable esophageal adenocarcinoma (EAC) patients who underwent first-line immunochemotherapy followed by chemotherapy; iii) Motzer et al. 2020^79^, consisting of 886 patients with advanced renal cell carcinoma (RCC) treated with anti-PD-L1 or anti-PD-1. We applied our above-mentioned iNMDeff proxy model based on gene-level expression (see Methods) to estimate iNMDeff for each patient in these cohorts, yielding iNMDeff estimates for 120 patients in Liu-SKCM dataset, 33 in Carroll-EAC dataset, and 353 in Motzer-RCC dataset.

The KM curves for PFS (Fig. 7D-F and Supp. Fig. S21D-F) demonstrated a consistent trend across the three immunotherapy studies for iNMDeff. Specifically, this trend was evident in melanoma (Fig. 7D and Supp. Fig. S21D; ETG p-value = 0.16 at 20th percentile, ASE p-value = 0.21, at 40th percentile), esophagus (Fig. 7E and Supp. Fig. S21E; discretized at percentile 25th, ETG p = 0.27 at 25th percentile, ASE p = 0.7), and renal cancer (Fig. 7F and Supp. Fig. S21F; ETG p = 0.17 at the 30th percentile, ASE p = 0.15 at the percentile 45th). The combined meta-analysis p-value (by Fisher’s method) from the 3 studies would be p = 0.13 and p = 0.27, for ETG and ASE, respectively. When analyzing the data using the Cox proportional hazards model, including covariates such as sex, age and TMB (see Methods) (Supp. Table S7), the HR from all three studies trended in the same direction for both NMD estimation methods in Liu-SKCM (ETG: HR = 1.68; ASE: HR = 1.37) and Motzer-RCC (1.29 and 1.23), and for the ETG method in Carroll-EAC (1.50; not supported in ASE).

Tumor mutation burden (TMB) is a well-established predictor of immunotherapy response, likely due to its association with neo-antigen appearance. We developed a predictive model that combines TMB and iNMDeff, aiming to classify immunotherapy responses into responders versus non-responders (see Methods). This model was applied across the previous datasets TCGA-SKCM, Carrol-EAC, Liu-SKCM, excluding Motzer-RCC due to missing TMB and treatment response data. Additionally we included new datasets of immunotherapy-treated patients from Hartwig Medical Foundation^80^ (urinary tract, n = 23, lung, n = 26 and melanoma, n = 119) and Riaz et al. (melanoma, n = 48)^81^, totaling seven datasets for this analysis.

The model’s predictive ability was significantly enhanced by including iNMDeff, showing a 7.4% mean increase (for both ASE and ETG) in discrimination capacity compared to a TMB-only model, averaging across the datasets (Fig. 7G and Supp. Table. S8). The area under the precision-recall curve (AUC) increased from 0.704 (TMB-only) to 0.778 (TMB + iNMDeff), averaged across the datasets. Notably, this improvement manifested variably across the different datasets: 1-9% in Riaz-SKCM, 11-27% in TCGA-SKCM, 1-2% in Hartwig-SKCM, 11-25% in Hartwig-UT, 3-5% in Hartwig-Lung and 1-10% in Carrol-EAC. Liu-SKCM did not exhibit any improvement. Regarding other performance metrics, averaged across datasets: recall (sensitivity) increased from 0.89 to a range of 0.85-0.91 (for ETG and ASE, respectively), and precision improved from 0.72 to a range of 0.74-0.78.

In conclusion, individual-level NMD efficiency can significantly impact cancer patient survival: there was a consistent trend of reduced PFS in immunotherapy-treated patients with high NMD efficiency, observed independently across 3 cancer types. Furthermore, iNMDeff can improve predictive models of immunotherapy response. We note there is a strong prior that a globally more efficient NMD in a patient would hinder immunotherapy, based on recent work associating NMD inactivity upon individual indel mutations in a tumor and immunotherapy response^11,44^. Plausibly, the underlying mechanism would be that in highly NMD-active tumors, a diminished expression of neoantigenic mutations produced by frameshifted and truncated mRNAs would occur. As an additional result, we also note high NMD efficiency might have adverse effects also in response to more conventional chemotherapies, with unclear underlying mechanisms.

## 3. Discussion

In our study, we quantified individual-level NMD efficiency in large-scale RNA-seq and germline WES data from normal tissues and cancers, totalling ∼27,000 samples. We considered both normal and mutated transcripts, to assess the variation in NMD efficiency across individuals and tissues.

We found a significant variability of iNMDeff across normal tissues and cancer types. This suggests tissue-specific regulation of the NMD pathway, which is broadly conserved between cancers and healthy tissues. Gastrointestinal tissues showed higher iNMDeff, and within them the hypermutated MSI colon and stomach cancers were among the ones with highest NMD efficiency. This is in agreement with a reported overexpression of NMD factors in MSI primary colon tumors, and that chemical inhibition of NMD caused a reduction in MSI tumor growth^40^. Together, this suggests that MSI tumors (speculatively, due to higher frameshift mutation burden) necessitate a higher NMD efficiency to survive, supporting that targeting the oncogenic activity of NMD could be a viable, personalized treatment strategy for some cancers^46^. Whether SNV/indel burden or other genomic features predict responses of an individual tumor to NMD inhibitiors remains to be determined in future work.

In contrast to the digestive tract, we observed that reproductive and nervous system tissues exhibited lower iNMDeff than other tissues, particularly so for neurons (as estimated from differential analysis of brain regions and glioma subtypes). A previous study in mice monitored the levels of *MEN1* gene^23^ and based on this reported NMD activity in brain, testis, ovary and heart, our comprehensive analysis across many human tissues suggests that considering multiple genes and mutations is necessary to reach robust conclusions on tissue specificity.

Aligning with our results, a recent study of ASE in GTex healthy tissues does rank the brain at the low end of NMD efficiency^53^. This might stem from the brain’s unique NMD regulation during development^82^, for instance through the role of miR-128’s targeting of *UPF1* to inhibit NMD, thus promoting neural differentiation, a mechanism persisting into adulthood^83^. In addition, other NMD-regulatory miRNA circuits have been discovered in the brain^84,85^. Together, ours and these previous findings indicate a reduced mRNA surveillance by NMD in mature neurons compared to other cell types. Extending this idea, developmental roles of NMD in other tissues such as blood^86^, muscle^87^, liver^88^, embryogenesis^89^, spermatozoa^90^ etc, might also contribute to the variability of NMD efficiency we observe across adult tissues. Lastly, we observed differences in NMD efficiency between some cancers and their matched normal tissues, suggesting a potential shift in NMD efficiency during transformation. For instance, while breast cancer tumors of the normal-like subtype, and their corresponding normal tissues align in NMD efficiency, the other breast cancer subtypes HER2, Luminal A, and Luminal B display higher iNMDeff; this is consistent with reports of higher *UPF1* expression in these subtypes^39^. This variation also underscores there may exist differences between cancer subtypes in NMD.

In addition to variation between tissues, variation in NMD efficiency between individuals is significant. Some of this is explained by association with somatic genetic variants in genes *TLX1* and *CDH1*; the latter is a known risk gene in hereditary diffuse gastric cancer and hereditary breast cancer^91–94^, and it is tempting to speculate that NMD has a mediating role in this^95^. We next extended the somatic variant analysis to CNAs, by a bespoke sparse-PCA-based method to control for genetic linkage in association studies, here applied to NMD efficiency variability. Three CNA-PCs reflecting large-scale gains were robustly associated with reduced NMD activity, most strongly CNA-PC3 and 86 located in regions 1q and 1q21-23.1. The 1q gain is found in ∼25% of cancers and it has been proposed to be selected through increasing dosage of the *MDM4* oncogene phenocopying *TP53* mutation^63^, or of *MCL-1* antiapoptotic factor^96^ or via upregulation of Notch genes^97^. This chromosome arm however also contains NMD factors *SMG5*, *SMG7*, the EJC gene *RBM8A*, and NMD-related genes *INTS3* and *SF3B4*, which we identified as associating with NMD efficiency upon CNA gain; additionally we identified 6 other genes involved in various RNA metabolic processes. A previous study identified significant enrichment of germline copy number variants of NMD genes, in neurodevelopmental disorder patients^98^, including *UPF2*, and also EJC genes including the chr 1q-located *RMB8A*. Interestingly, both gains and losses of the same genes, for example, *UPF2* and *RMB8A*, were linked to the phenotype, implying that NMD and EJC gene dosage imbalances impact NMD function and disease pathology. As for somatic variation, it was suggested that tumor evolution favors mutations and CNA co-occurring in NMD factor genes^20^. Interpreting the effects of NMD factor genetic alterations on NMD efficiency and downstream phenotypes (e.g. cancer patient survival) requires future work on modelling an extensive negative feedback regulatory network impacting NMD factors, and its dynamics^52^. This feedback mechanism, which varies by cell type and developmental stage, serves to mitigate disruptions in NMD factor genes, and our data suggests that this mechanism does not fully counterbalance the effects of chromosome gains or co-occurrence mutations that affect multiple NMD factors and/or NMD-related genes. In addition to 1q gains, further the chromosome 2q gains that we find associated with NMD efficiency are interesting to follow-up, since this arm encompasses candidate genes *CWC22* and *SF3B1* implicated in the splicing process, whose dysregulation may plausibly affect NMD. It has been shown that mutations in the spliceosome genes *SF3B1* and *U2AF1* render cancer cells more sensitive to NMD inhibition in a synthetic lethality fashion^30,99,100^ adding confidence that CNA of this same gene might disrupt NMD.

In addition to somatic variants in tumors, also the rare deleterious germline variants can affect NMD efficiency in at least the thyroid and brain. We identified 10 genes with diverse functions, potentially affecting NMD efficiency indirectly. For instance, *CRTC1*, primarily active in the brain subregions, acts as a transcriptional activator^101^. *NUP153* is involved in mRNA nuclear export and neural progenitor cell regulation^102^. *KDM6B*, a histone demethylase, is pivotal in gene expression regulation by removing the Polycomb repressive histone marks ^103,104^, and also linked to neurodevelopmental disorders^105^. We also identified genes whose mechanistic links to NMD are less obvious; replication in additional cohorts would be the sensible next step. A recent study identified genetic variants that influence NMD-targeted transcripts and their NMD decay efficiency across healthy GTex tissues^106^, with more than 50% of NMD-QTLs identified in the brain in particular. Interestingly, *SF3B4*, one of our identified candidates from somatic associations, emerged as the gene containing the highest number of germline NMD-QTLs.

Next, our analysis suggests various associations of differential NMD efficiency to tumor evolution: shaping selective pressures on tumor suppressor genes, with prominent effects on *SMAD4*, *TP53*, and *APC*. In a cancer-type specific analysis, we observed a similar trend in hypermutated tumors like colon, stomach MSI and also lung cancers, where these TSGs were more frequently mutated. In these individuals/tumors which are more adept at utilizing NMD mechanisms to ablate the activity of these tumor suppressor genes, applying NMD inhibition therapeutically -- alone or in combination with readthrough agents^107–109^ -- may be promising in restoring tumor suppression activity to treat cancer.

In support of the idea that manipulating NMD may treat some cancers, we observed that the naturally variable NMD efficiency is associated with cancer survival outcomes (in agreement with our recent population genomic study arguing that NMD modulates phenotype severity of various heritable genetic diseases^11^). We found that high NMD efficiency is associated with improved or worse outcomes depending on cancer type, consistent with a previously proposed complex, multifaceted roles of NMD influencing in cancer^30,74^. Our analysis particularly highlights the effects of the global NMD efficiency on responses to immunotherapy. Previous studies suggested that NMD-evading status of individual mutations^11,44^ does predict response to immunotherapy; here we present evidence of an additional role of the global, individual-level NMD efficiency in this, observed across various patient cohorts and cancer types. A well plausible mechanism is due to NMD’s role in silencing the presentation of neoantigens. Indeed this is consistent with our observation that predictive models containing interaction terms between TMB and iNMDeff have higher accuracy in predicting responses. We do note iNMDeff associations with efficiency of conventional chemotherapy, which may or may not be mediated by immune mechanisms. Speculatively, NMD may confer a generally increased tumor cell fitness in the context of occurrence of deleterious, mutated proteins that arose via somatic mutation (as shown before for one example, the HSP110DE9 chaperone mutant^40^), conferring additional ability to withstand various challenges to cancer cell fitness.

In summary, our study reveals extensive variation in NMD efficiency across individuals, tissues and tumors, resulting in part from genetic alterations, and underscores the potential of NMD efficiency estimates as a predictive biomarker for cancer treatment using immunotherapy and/or NMD inhibition.

## 4. Methods

### Sequencing data

We downloaded matched tumor and normal whole-exome sequencing (WES) *bam* files, along with tumor RNA-seq *fastq* files, from the TCGA consortium via the Genomic Data Commons (GDC) Data Portal^110^. This dataset encompassed 9,766 samples across 33 distinct cancer types. Copy number alteration (CNA) data, including both arm-level and focal-level information, was obtained from the Broad Institute’s GDAC Firehose portal^111^. From GTex, we directly obtained WES *VCF* files and tissue-specific transcript-level counts RNA-seq data, accessed through dbGaP (via AnVIL), v8 (dated 05-06-2017). This dataset included 979 individuals and spanned 56 different tissues, amounting to a total of 17,382 samples, with an average of 19 tissue types per individual. We also downloaded the ASE counts data for 838 individuals.

RNA-seq and CNA data of 1450 human cell lines from the Cancer cell line encyclopedia (CCLE)^62^ was also obtained.

### Clinical and other data sources

TCGA cancer cluster subtypes, determined based on mRNA non-matrix factorization (NMF), were sourced from the Broad Institute’s GDAC Firehose portal^111^. In addition, we acquired estimates of leukocyte fraction^112^ and tumor purity^113^ for all samples within the TCGA dataset. Patient clinical data and metadata were obtained directly from the GDC portal^111^. For progression-free survival (PFS) information, we relied on data available through cBioPortal^114^ [<u>REF</u>].

### TCGA cancer type stratification into subtypes

The classification of TCGA cancer molecular subtypes was meticulously gathered using functions from the *TCGABiolinks* R package, namely *PanCancerAtlas_subtypes* (only for BRCA) and *TCGAquery_subtype*, or directly from the referenced articles in certain cases. The classifications are as follows:

BRCA: Classified according to the PAM50 mRNA profiling into five subtypes: Basal, Normal-like, Luminal A, Luminal B, and HER2-enriched^115^.
SARC: Based on histological subtypes^116^, with specific subtypes such as Soft tissue leiomyosarcoma (STLMS) and Uterine leiomyosarcoma (ULMS) being aggregated into the ‘Muscle’ category, and Dedifferentiated liposarcoma (DDLPS) classified under ‘Fat’.
BLCA: Utilized mRNA clustering to divide samples into Basal squamous cell, Luminal papillary, Luminal infiltrated, and Luminal subtypes^117^.
HNSC: Categorized based on RNA-seq data into HPV positive or negative groups^118^.
COAD, STAD, and UCEC: These cancers were classified into MSI or MSS^119–121^. For UCEC, an additional distinction was made for samples with POLE mutations, predominantly within the MSS subgroup.
ESCA: Classified into esophageal squamous-cell carcinoma (SCC) and adenocarcinoma (AC)^122^.
GBM: Subdivided based on the Glioma CpG island methylator phenotype into High (G-CIMP-high) and Low (G-CIMP-low), alongside methylation cluster 6 (LGm6-GBM), Classic-like, and Mesenchymal-like^57^.
LGG (Lower Grade Glioma): Classified into G-CIMP-high, G-CIMP-low, Classic-like, Mesenchymal-like, PA-like, and 1p/19q codeletion (Codel)^57^.

### Alignment and quantification of RNA-seq and ASE

In adherence to GTex guidelines^123^, we processed TCGA RNA-seq data by aligning the reads to the human genome *GRCh38.d1.vd1*, utilizing *STAR* v2.5.3a^124^. Subsequent transcript-level quantification was conducted using *RSEM* v1.3.0^125^. For allele-specific expression (ASE) analysis, we retained RNA-seq reads with a minimum mapping quality of 255. To correct for allele-mapping bias, we employed *WASP* for adjustment^126^. Additionally, an unbiased approach for the removal of duplicate reads was implemented using a custom *WASP* script. This script randomly discards duplicate reads, ensuring that the removal process is independent of the read scores.

### Variant calling and annotations

Variant calling in TCGA was done using *Strelka2* v2.9.10^127^. For somatic WES, normal and tumor bam files were compared utilizing the *configureStrelkaSomaticWorkflow.py* script, employing the ‘--exome’, ‘--tumorBam’, and ‘--normalBam’ flags. In the case of germline WES, the analysis was conducted using only normal bam files, applying the *configureStrelkaGermlineWorkflow.py* script with ‘--exome’ and ‘--bam’ flags. For RNA-seq data, aimed at obtaining ASE counts, we again used normal bam files, this time with the *configureStrelkaGermlineWorkflow.py* script and the ‘--exome’, ‘--rna’, and ‘--forcedGT’ flags. The latter flag forces genotype calling to conform to previously identified genotypes from WES. Single nucleotide variants (SNVs) and insertions-deletions (INDELs) were called separately in all instances and subsequently merged. We annotated all resulting VCF files using ANNOVAR^128^ (2020-06-07 version), referencing GENCODE v26 (ENSEMBL v88), and incorporated the *gnomad211_exome* database to acquire population minor allele frequencies (MAF) from gnomAD^129^. In the case of GTex, as previously mentioned, the WES VCFs, RNA-seq counts data, and ASE counts were directly downloaded. The only additional step undertaken was a liftover to convert genome coordinates from GRCh37 to GRCh38. Regions with poor alignment between the genome versions were systematically identified and removed to ensure liftover accuracy^130^. Subsequently, the variants were annotated in the same manner as in TCGA, as described earlier.

### Obtaining NMD target gene sets from various studies

We created different gene sets of NMD targets from a range of studies employing both experimental and computational methods, to use in the estimation of ETG iNMDeff:

1) Tani H. et al. (2012)^6^ identified *UPF1* NMD targets categorized into three groups: A) 248 indirect targets with over 2-fold upregulation but stable decay rates; B) 709 direct targets with unaltered expression levels but over 2-fold increase in decay rates; and C) 76 bona-fide direct *UPF1* targets with both over 2-fold upregulation and extended decay rates. We excluded the genes from group B in subsequent analyses due to their lower confidence as NMD targets.
2) Colombo M. et al. (2017)^5^ provided *UPF1*, *SMG6*, and *SMG7* specific NMD target genes. For *UPF1* targets, we selected those with a ‘meta_pvalue’ < 0.05, resulting in 2725 genes. For *SMG6* targets, we applied a criteria of ‘UPF1_FDR’ < 0.05 and ‘meta_SMG6’ < 0.05, yielding 1780 genes. Likewise, for SMG7, we required ‘UPF1_FDR’ < 0.05 and ‘meta_SMG7’ < 0.05, leading to a list of 1047 genes.
3) Karousis E. D. et al. (2021)^13^ utilized nanopore sequencing and compiled a gene set pairing NMD target transcripts with controls, encompassing 1911 genes.
4) Courtney et al. (2020)^51^ identified a set of 2793 NMD target genes.
5) Schmidth S.A et al. (2015)^14^ listed 233 SMG6 specific target genes.
6) The ENSEMBL gene annotation file tags some genes as “nonsense_mediated_decay”.

With all this information, we created distinct NMD gene sets for each study, with the exception of Schmidt. These sets are named: “NMD Tani”, “NMD Colombo”, “NMD Karousis”, “NMD Courtney”, and “NMD Ensembl”. Additionally, we constructed specific gene sets for “SMG6” and “SMG7” NMD targets. For “SMG6”, we derived the set from the intersection of SMG6 NMD targets identified in both Colombo’s and Schmidt’s studies. For “SMG7”, we exclusively utilized the SMG7 NMD target genes from Colombo’s study. Additionally, we overlapped genes found in at least 2 studies to compile a “NMD Consensus” gene set, and the complete list of genes across all studies as “NMD All” gene set. Lastly, we derived negative control gene sets, by selecting genes with and without NMD-triggering features (see the following section) in their transcripts and excluding the NMD target gene sets from above: “RandomGenes with NMD features”, and “RandomGenes without NMD features”.

### Predicting NMD-triggering features for the selection of endogenous NMD targets and controls (ETG method)

For each gene within our NMD gene sets from the ETG method, we classified its transcripts into NMD targets and controls based on computationally predicted NMD-triggering features. This classification relied on two primary features identified in the literature: i) the presence of an upstream open-reading frame (uORF) in the 5’ untranslated region (UTR)^15,16^ that does not overlap the CDS, and ii) the existence of at least one splice site (or exon-junction complex, EJC) in the 3’ UTR, located more than 50 nt downstream of the termination codon^12,17^. We also considered the GC content of the 3’ UTR, known to influence transcript degradation by NMD^18^. To delineate NMD target-control pairs for each gene, our specific criteria is shown in Supp. Fig. 1B. In summary, the criteria for NMD target transcripts are as follows:

i) Presence of both a start codon and a stop codon.
ii) A median transcript expression (log TPM) across pan-cancer (TCGA) of >= 1.
iii) Presence of either a 3’UTR splice site or >= 2 uORFs (both having a minimum length of 30 nt).
iv) If multiple transcripts meet the above criteria, the one with the highest 3’ UTR GC content is selected.

For control transcripts, the requirements are:

i) Presence of both a start codon and a stop codon.
ii) A median transcript expression (log TPM) across pan-cancer (TCGA) of >= 3.
iii) Absence of any NMD-triggering features.

Lastly, for a transcript to be classified as an NMD target, and a transcript to be its paired control, the ratio of their expressions must be <= 0.9. This threshold ensures that the selected NMD target transcripts exhibit lower expression levels compared to their corresponding controls, aligning with the expected impact of NMD-triggering features on transcript stability.

To enhance the accuracy of this selection and mitigate the inclusion of potential false positives, we utilized data from cell lines with *UPF1* knockdown (KD) via an inhibitor, as reported in four different studies: HeLa cells^5,131^, HepG2^132^, K562^132^. For each selected transcript pair, we stipulated that the expression ratio in the four wild-type cell lines must average <= 0.9, reflecting the anticipation of high NMD efficiency in the control cell lines. To acquire the necessary RNA-seq count data, we conducted our own alignment and quantification using the same tools and versions previously outlined for the TCGA and GTex databases (as detailed in the section ‘Alignment and quantification of RNA-seq and ASE’).

It is important to note that not all selected transcript pairs will be utilized in our ETG iNMDeff, due to the filterings performed afterwards (see section “Filterings for the ETG method”).

### Using NMD rules for predicting NMD-triggering vs NMD-evading PTCs (ASE method)

For the ASE method, we determined whether a PTC, resulting from nonsense or indel mutations, is likely to undergo NMD based on its location within the coding sequence (CDS) of the transcript. For indels, we predicted the potential generation of a downstream PTC and identified its exact location. To categorize these PTCs as either NMD-triggering or NMD-evading, we relied on established NMD rules^8^, except the long-exon rule:

i) The 55nt-rule: PTCs positioned <= 55nt downstream from the last base of the penultimate exon are considered NMD-evading.
ii) The last exon rule: PTCs located within the last exon are deemed NMD-evading.
iii) The start-proximal rule: PTCs situated within the first 200 nucleotides of the transcript are classified as NMD-evading.

In scenarios where a transcript has a splice site in the 3’UTR, a PTC that would normally be classified as NMD-evading under the 55nt or last exon rule is instead considered NMD-triggering. This reclassification is due to the presence of an additional EJC located downstream of the PTC.

Using this classification, we generated three distinct NMD variant sets for each individual: ‘NMD-triggering PTCs’, ‘NMD-evading PTCs’, and ‘Synonymous’. These sets were specifically designed to accurately estimate the NMD efficiency for each individual, along with its two negative control sets.

### Filterings for the ETG method

To ensure an accurate estimation of ETG iNMDeff, all filtering processes were conducted on a per-individual basis and on all NMD gene sets separately. The following criteria were applied in both TCGA and GTex cohorts:

- We retained transcripts exhibiting a log2(raw counts) >= 1 in at least 50% of individuals within the specific cancer type of the individual being analyzed. If both transcripts in any NMD target-control pair failed to meet this threshold, the entire pair was excluded.
- Non-coding transcripts were excluded from the analysis.
- We discarded transcripts that overlapped with any type of somatic (only in TCGA) or germline truncating mutations (such as nonsense, start loss, nonstop mutations, inframe insertions and deletions, frameshift insertions and deletions, and splice site mutations) as well as those affected by CNAs (only on TCGA).
- A minimum of 2 pairs of NMD target-control transcripts was required for the estimation of an individual’s NMD efficiency.
- For all NMD gene sets, except from the “NMD Consensus” set, we randomly selected up to 50 pairs of NMD target-control transcripts. This approach was adopted to streamline the computational process.

### Filterings for the ASE method

Similar to the ETG method, in the ASE approach, all filterings were meticulously carried out on a per-individual basis and for each NMD germline variant set (“NMD-triggering PTCs”, “NMD-evading PTCs” and “Synonymous”). The following criteria were applied in both TCGA and GTex cohorts:

- We retained only those heterozygous variants that met the “PASS” threshold in the *VCF* file.
- A minimum total read coverage (sum of WT and MUT alleles count) requirement was set at 5 for nonsense and synonymous mutations, and at 2 for indels.
- Variants with a MAF > 20% were excluded to avoid confounding effects in the analysis. Common variants are known to exhibit lower NMD efficiency, likely due to evolutionary selection^8,10,11,53,133^.
- Variants located in non-coding transcripts or in transcripts consisting of a single exon were excluded.
- Variants in 331 genes identified as undergoing positive selection were removed^134^.
- For variants in genes experiencing negative selection, we employed the loss-of-function observed/expected upper bound fraction (LOEUF) estimate^135^. Genes in the lowest first percentile based on this score, representing the 10% most constrained genes, were excluded from the analysis.
- We excluded variants co-occurring with any type of somatic truncating mutation (such as nonsense, start loss, nonstop mutations, inframe insertions and deletions, frameshift insertions and deletions, and splice site mutations) in TCGA. This also included variants affected by CNAs in TCGA.
- Specifically for the ‘NMD-triggering PTCs’ variant set, further exclusions were made:

◦ NMD-evading PTCs (and vice versa).
◦ Frameshift variants that do not predict a downstream PTC.
- For the ‘Synonymous’ variant set:

◦ We excluded variants overlapping any gene from our ‘NMD All’ gene set to avoid confounding the analysis.
- In GTex only, if the variant only appeared in one tissue, then it was considered a somatic variant, and not germline, thus discarded.
- A minimum of 3 variant PTC pairs (WT-MUT) was required for the estimation of an individual’s NMD efficiency.
- Additionally, to optimize computational efficiency, we capped the number of variant pairs used in the analysis randomly sampling a maximum of 100.

### Quantification of individual NMD efficiency (iNMDeff) by a negative binomial regression

To estimate individual NMD efficiency (iNMDeff), we employed Bayesian generalized linear models, fitting a negative binomial distribution using “*Stan*”^136^. This modeling was implemented via the ‘stan_glm’ function from the *rstanarm* R package^137^, with ‘family = neg_binomial_2’ specified as the parameter.

For the ETG method, the model is applied, pooling all transcripts together within a sample, for each of the 11 NMD gene sets (includes the negative control) separately, as follows:

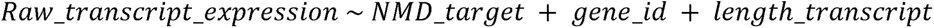

Where:

- *Raw_transcript_expression*: represents the raw count data from RNA-seq for each transcript.
- *NMD_target* : indicates whether the transcript is an NMD target (1) or the control (0) within the selected pair.
- *gene_id* : is the ENSEMBL gene ID included to adjust for between-gene differences.
- *length_transcript*: accounts for the total transcript length, calculated as the sum of base pairs of its exons. This adjustment is necessary to address the potential bias of more frequent read clustering in larger transcripts.

By comparing each NMD target transcript against its paired control from the same gene, we establish an internal control. This approach effectively accounts for potential confounders affecting trans-gene expression levels. For instance, CNAs or transcription factors might alter the expression of one transcript without affecting the other. Such discrepancies are particularly pertinent if comparing transcripts across different genes. Although we already exclude genes overlapping with CNAs, this internal control further ensures the robustness of our analysis against such confounding factors.

For the ASE method, the model is applied, pooling all PTCs together within a sample, for each of the 3 NMD variant sets (includes the negative control) separately, as follows:

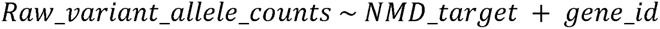

Where:

- *Raw_variant_allele_counts*: represents the allele specific expression raw counts of the germline PTC.
- *NMD_target* : indicates whether the allele is MUT (1), thus, NMD target, or WT (0), thus, control, within the selected pair.
- *gene_id* : is the ENSEMBL gene ID included to adjust for between-gene differences.

In both NMD methods, the coefficient assigned to the ‘NMD_target’ variable serves as our estimate of iNMDeff for a specific NMD variant or gene set, as well as its corresponding negative control. We reversed the direction of the raw coefficient values, so that now higher coefficients indicate greater iNMDeff, and lower coefficients indicate reduced efficiency. The final interpretation is that it is a negative log (base e) ratio of the raw expression levels of the NMD target transcripts (ETG) or MUT alleles (ASE) divided by the control transcripts (ETG) or WT alleles (ASE). For a more intuitive interpretation, one could exponentiate the coefficient to derive the ratio between NMD targets and controls. In this context, ratios above 1 would suggest lower NMD efficiency, while ratios below 1 would indicate higher NMD efficiency. It is important to note that for our analysis, we utilized the original log coefficients rather than these exponentiated ratios.

Interestingly, both iNMDeff methods suggest a possible differential NMD activity across tissues or individuals, and the differences in their NMD estimates might reflect differently active NMD subpathways or other types of biological or technical variation. The ETG method, based on transcript-level expression, offers a high data point count per individual, crucial for estimating iNMDeff accurately, but faces challenges like confounding factors such as CNAs and varying expression levels among transcripts of a gene, which we attempted to address with rigorous filtering and pairing NMD-target transcripts with appropriate controls. Conversely, the ASE method, while robust to confounders due to its internally-controlled quantification of the two alleles within the same transcript, is limited to more highly expressed genes and thus provides fewer data points, leading to potential noise issues. We used both methods as independent approaches, ensuring a more reliable analysis of NMD efficiency across individuals.

### Randomization tests for iNMDeff variability

#### Inter-Tissue iNMDeff variability deviation (ITNVD) test

This test consists in shuffling the data 2000 iterations, to see if the differences observed between tissues could have happened by chance (see also Fig. 3A). For each shuffle, we calculated the variability (standard deviation, SD) of the median iNMDeff across samples for each tissue type, to obtain a single value per iteration. We compared the median of the 2000 randomized variability scores to the observed variability in the original, unshuffled data, and termed the difference between them as the “Inter-tissue iNMDeff Variability Deviation (ITNVD)”, which serves as a measure of effect size. We then calculated a p-value to assess the statistical significance of this effect size. If there is no variability, the value should be close to 0.

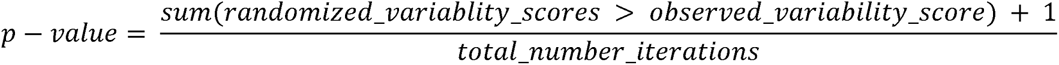

#### Tissue iNMDeff Deviation (TND)

This test compares the observed median iNMDeff for each tissue against a randomized median iNMDeff (note that these medians were essential in deriving the ITNVD score discussed earlier), as illustrated in Fig. 3A. The resulting difference between them is termed “Tissue iNMDeff Deviation (TND)”. We calculated this deviation score 2000 times, once for each reshuffling of the data, to create a comprehensive distribution of these deviations. Essentially, a positive value indicates that the tissue has a higher iNMDeff than expected by chance, and viceversa. As before, we then calculated a p-value to assess the statistical significance of this effect size.

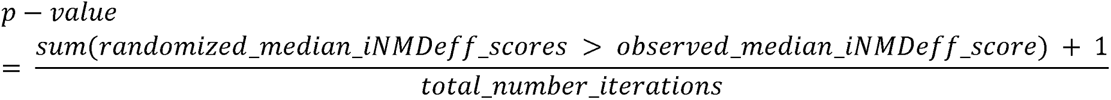

### Tissue variability of iNMDeff using linear models

As an alternative approach to assess the variability of iNMDeff across different tissues and cancer subtypes, we employed a linear modeling approach. This model aimed to predict iNMDeff values, based on a set of explanatory variables. These included demographic factors (age, sex), technical aspects (RNA-seq sample library size), and others (number of NMD targets as measured in ETG or number of PTCs as measured in ASE). For cancer samples from the TCGA, we further adjusted for other variables such as tumor purity, CNA burden, race, TMB, categories of POLE mutations, vital status, and all 86 pan-cancer CNA-PCs. In contrast, for the GTex, the model also accounted for the Death Group classification.

In constructing the model for each NMD method (ETG and ASE) and cohort (TCGA and GTex) separately, we adopted a rigorous approach. After establishing the initial model incorporating all variables, we systematically eliminated each variable in turn. This stepwise removal allowed us to evaluate the impact of each variable on the model’s overall explanatory power, specifically measuring the reduction in the total variance explained (R²) by the model (Supp. Table S1).

### Cellular deconvolution for TCGA brain tumors

In our study, we applied the UniCell: Deconvolve Base (*UCDBase*) method^56^ to perform bulk RNA-seq deconvolution across the entire TCGA cohort. For this particular analysis, pan-cancer RNA-seq was directly downloaded from XENA browser^138^, as the RNA-seq data underwent normalization to mitigate batch effects, followed by a log2 transformation of normalized values (log2(norm_value+1)), prior to deconvolution. This process resulted in a comprehensive matrix encompassing 10,460 samples, which spanned 708 distinct cell types across various hierarchical levels. Our analysis specifically targeted samples from GBM and LGG to assess cell type composition within these tumor types, specifically for: “astrocyte_of_the_cerebral_cortex”, “Bergmann_glial_cell”, “brain_pericyte”, “endothelial_cell”, “neuron”, “oligodendrocyte”, and “oligodendrocyte_precursor_cell”.

### Quantification of ASE PTC-NMD efficiency (pNMDeff) for inter-individual NMD efficiency variation

To assess the inter-individual variability of NMD efficiency, we quantified the NMD efficiency for each PTC arising from nonsense mutations or indels, a process we refer to as pNMDeff, following the approach previously described by our group^8^, applying ASE data instead of transcript-level counts. This methodology was utilized for evaluating germline ASE PTCs across both TCGA and GTex cohorts, and for somatic PTCs within TCGA. The analysis entailed comparing the expression (raw counts) of mutated (MUT) versus wild-type (WT) alleles at the PTC site:

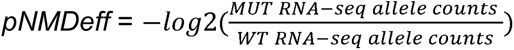

A pNMDeff score of 0 indicates no mRNA degradation, while a score of 1 indicates complete heterozygous mRNA degradation via NMD.

PTCs or transcripts were systematically excluded from our analysis based on several stringent conditions:

- Exclusion of indels without a downstream PTC prediction.
- Exclusion for additional germline or somatic indel, nonsense, or splice site disrupting variants (not applicable to somatic PTCs in TCGA), or overlap with somatic CNAs.
- Exclusion of PTC-bearing transcripts comprising a single exon without additional 3’ UTR splice sites.
- Exclusion based on the gene’s LOEUF score position in the first decile, indicating negative selection, or identification of the gene under positive selection.
- Exclusion of homozygous PTC variants.
- Exclusion of PTCs with a variant allele frequency > 20% (for somatic variants in TCGA).
- Exclusion of transcripts with median pan-tissue or pan-cancer expression level < 5, or significant expression variation among samples (coefficient of variation > 0.5).

Utilizing pNMDeff data from germline ASE in TCGA & GTex, and somatic ASE in TCGA, we assessed inter- and intra-individual variability of NMD efficiency, focusing on NMD-triggering PTCs as per canonical NMD rules^8^. We applied two metrics: one based on variance and another on correlations.

#### Intra-individual variability

we sampled two PTCs per individual, calculating the Spearman correlation between their pNMDeff scores across 1000 iterations.

#### Inter-individual variability

we compared two sampled individuals for the same common PTC, again calculating Spearman correlation between their pNMDeff scores, also across 1000 iterations.

Variance-based metrics involved calculating the variance of pNMDeff scores for either at least 3 PTCs within individuals or across individuals sharing the same PTC.

Each analysis was conducted separately for each cohort and was supplemented by a control analysis involving randomized pNMDeff values to establish a baseline for comparison.

### NMD-related genes classification

We integrated data from 112 NMD-related genes to investigate gene-level associations with somatic mutations and to evaluate CRISPR codependency scores for identifying causal genes. This selection of NMD genes was informed by two pivotal studies with experimental validations. From Baird et al., 2018^59^, we selected the top 76 genes identified via siRNA screening, focusing on those with a median seed-corrected Z-score above 1.5. Alexandrov et al., 2017^60^ contributed an additional 24 top genes. Our selection also encompasses core NMD factors identified through extensive literature review^3,7,61^, including *SMG1*, *SMG5*, *SMG6*, *SMG7*, *SMG8*, *SMG9*, *UPF1*, *UPF2*, *UPF3A*, and *UPF3B*, crucial for the NMD mechanism. We included core EJC factors such as *CASC3*, *EIF4A3*, *MAGOH*, and *RBM8A*, essential for RNA splicing and surveillance, alongside EJC-associated genes *ACIN1*, *ALYREF*, *AQR*, *CWC22*, *RNPS1*, *SRRM1*, which interact with or might be part of the EJC. The DECID complex, involving *DHX34*, *ETF1*, *GSTP1*, plays a role in distinguishing defective mRNAs, and genes within the NMD subpathway in the endoplasmic reticulum, *GNL2*, *NBAS*, *SEC13*, were also included in our analysis.

### Association analysis of somatic mutations

The association between iNMDeff and somatic mutations was investigated using general linear models (*‘glm’* function from the R package *“stats”*), applied to each gene within 33 cancer types and in a pan-cancer dataset. Covariates for individual cancer types included sample purity, the first 86 principal components of CNA-PCs, and cancer subtype based on mRNA NMF clustering. In the pan-cancer analysis, additional covariates such as CNA burden, RNA-seq sample library size, TMB, age, and sex were incorporated to address result inflation (Supp. Fig. S8). We tested all the genes but our focus was on: i) a set of 727 recognized cancer genes from the CGC^58^; ii) 112 genes related to the NMD pathway (see above); iii) the remaining 18,780 genes were considered as random genes for our analysis. Each gene was subjected to three separate analyses involving the following sets of genetic variants: (i) truncating mutations (including nonsense, indels, and splicing variants) and non-synonymous mutations, both tested within the same model unless one mutation type was absent, (ii) synonymous mutations, and (iii) somatic CNAs, where the variable indicated gene amplification (GISTIC CNA score > 0) or deletion (GISTIC CNA score < 0). In scenarios (i) and (ii), the variable was set to 1 if any mutation was present in the gene, otherwise 0.

### Somatic pan-cancer CNA signatures by sparse-PCA

We performed a sparse principal component analysis (sparse-PCA) on the 24,777 gene CNA arm-level data in 10,654 TCGA samples, using “*sparsepca”* R package^139^. Of note, we duplicated the rows by splitting genes into amplifications/neutral (keeping GISTIC CNA scores >= 0, negative scores were transformed to 0) and deletions/neutral (keeping GISTIC CNA scores <= 0, positive scores were transformed to 0), effectively obtaining a matrix of 49,554 x 10,654 dimensions.

In essence, sparse-PCA differs from standard PCA by performing implicit feature selection, focusing only on the most relevant genes that contribute significantly to data variation. This approach is particularly effective for high-dimensional data, as it efficiently manages sparsity, isolates key variables, reduces noise, and improves the interpretability of the results. In sparse-PCA, two parameters are pivotal: ‘*alpha’* and *’k’*. The *’alpha’* parameter governs the sparsity level of the principal components. Higher alpha values induce increased sparsity, thereby reducing the number of genes contributing significantly (non-zero weights) to each principal component (PC). In essence, “*alpha*” serves as a tool for regularization. On the other hand, “*k*” determines the number of sparse PCs to be calculated. The actual number of principal components with non-zero weights are what we refer to as effective PCs. To empirically optimize these parameters, we employed an autocorrelation scoring approach, focusing on the spatial relationship between neighboring genes. Autocorrelation in this context involves comparing each resultant PC, arranged according to genomic location, against a version of itself that is lagged by one gene position. High autocorrelation is anticipated when numerous genes, especially those in proximity, exhibit similar or identical weights due to being affected by the same arm-level change. Therefore, we expect the correlation between each original sparse-PC and its re-ordered version to approach a value near 1. This was quantitatively assessed by calculating the median autocorrelation of the lowest 1% percentile of effective PCs across various “*alpha*” and “*k*” values (as depicted in Supp. Fig. S9A). Our findings indicated that an “*alpha*” of 3e-04 and “*k*” of 100 yielded 86 effective pan-cancer PCs (with the remaining 14 PCs having gene weights of zero), while maintaining a median autocorrelation for the first 1% of PCs close to 1.

Each component in the CNA-PC might represent either a genomic amplification (Supp. Fig. S9B, left panel) or deletion (Supp. Fig. S9B, right panel), necessitating careful interpretation of the scores and their signs on a case-by-case basis. For example, a negative score in the CNA-PC does not automatically imply a deletion; it could be associated with an amplification, such as that of chromosome 2p. Therefore, the specific context and characteristics of each CNA-PC must be thoroughly examined to accurately determine whether the observed score reflects an amplification or a deletion in the genome.

### Association analysis of somatic pan-cancer CNA signatures

The association between individual weights of the pan-cancer CNA-PCs and our estimates of iNMDeff for both NMD methods was analyzed in 33 individual cancer types, and pan-cancer, from the TCGA. This analysis was conducted using general linear models with the ‘*glm*’ function from the R base package *‘stats’*. We incorporated several covariates into our models to control for potential confounding factors. These covariates included sample purity, cancer subtype based on mRNA NMF clustering, RNA-seq sample library size, age, and when applicable, sex. Effect size was determined by the beta coefficient of the CNA-PC variable. In the context of our model, a positive coefficient indicates that higher values of the specific CNA signature are correlated with increased NMD efficiency in individuals. It’s important to note that these CNA signatures could represent either genomic amplifications or deletions, hence the direction of the association should be interpreted accordingly.

In our analysis, ASE iNMDeff served as the initial discovery phase, leading to the identification of significant cancer-or pan-cancer-specific CNA-PC associations which were then validated using ETG iNMDeff, with FDR adjustments made only for these associations. This approach identified nine total CNA-PCs significantly associated with iNMDeff at an FDR below 10%. Of these, only three pan-cancer CNA-PCs (3, 52, and 86) exhibited consistent directions in their association with NMD efficiency, indicated by the same direction of effect sizes between ASE and ETG. In contrast, the remaining six CNA-PCs, which included three cancer-type specific (one in PCPG and two in UCEC) and three pan-cancer CNA-PCs, showed discrepancies in the direction of NMD effect and were therefore not considered in further analyses.

### Prediction of iNMDeff based on gene expression for validations

We developed LASSO or ridge regression models to predict iNMDeff based on gene-level expression data. The model was constructed separately for our two NMD methods, ETG and ASE, utilizing gene expression data from either TCGA or GTex. Prior to model development, genes with low expression (log2(TPM) >= 0.5 in fewer than 50% of pan-tissue samples) and those with low variance were excluded. The dataset was split into a training set (70% of the data) and a test set (30%). Additional preprocessing steps were taken before model construction to account for differences between cohorts, i.e. tumor (TCGA) or tissue samples (GTex) and the other dataset where we wanted to predict iNMDeff. This included correcting the RNA-seq counts matrix through quantile normalization and *comBat*^140^, following the methodology outlined in Salvadores et. al^141^.

These models were utilized in two distinct validation scenarios: i) validating chr1q amplification associated with reduced iNMDeff in the cell lines from the CCLE, and ii) validating the association of iNMDeff with PFS in external datasets from various studies. For scenario i), we predicted the cell line NMD efficiency (cNMDeff) for 942 cells. For scenario ii) we estimated the iNMDeff for 120 patients in Liu et al. melanoma dataset, 32 in Carroll et al. esophagus dataset, and 725 in Motzer et al. kidney dataset.

### Gene codependency CRISPR scores with NMD activity in cell line screens

To delineate NMD-associated genes, we leveraged genetic interaction data derived from gene-level CRISPR screening, as provided by the Achilles Project^66^. The initial dataset characterized negative scores as indicators of cellular growth inhibition or lethality post-gene knockout. To enhance the analytical robustness and infer gene functionality more accurately, we opted for a normalized, de-biased dataset employing the onion method and Robust PCA (RPCO), renowned for its efficacy in deducing gene roles. This adjustment yields a more intricate interpretation of the scores.

From the “Outputs generated by normalization pipelines - Normalized networks (ONION)” section^142^, we accessed the symmetrical matrix file “*snf_run_rpca_7_5_5.Rdata*”, encompassing data for 18,119 genes. Utilizing this matrix, we performed heatmap clustering (using ‘*ggcorrplot*’ function from the “*ggcorrplot*” R package) to visually represent the relationships among candidate genes implicated in chromosomal gains, alongside core NMD genes, and a set of randomly selected control genes for comparative analysis. For clarity in visualization, scores above 0.1 were capped at this value. Specifically, for chromosome 1q gains associated with CNA-PC3, we analyzed 6 out of 30 candidate genes, discarding those that did not cluster effectively for visualization. This subset included 23 NMD core genes—comprising 10 essential NMD factors, 4 EJC components, and 6 related genes, alongside 3 DECID complex members— and 18 control genes situated on chromosome 1q but outside the 1q21.1-23.1 region, excluding adjacent regions (1q23.2-1q31.3) for rigor. Similarly, for chromosome 2q gains linked to CNA-PC52, all 18 candidate genes were evaluated alongside the same set of 23 NMD core genes and 22 control genes outside the specific 2q31.1-2q36.3 region.

It’s important to note that the KO of some NMD factors, especially *UPF1*, may be lethal to cells, which could explain the lower similarity scores observed among NMD genes in CRISPR screens, which fully ablate gene activity.

Furthermore, we quantitatively assessed the NMD association by calculating the mean CRISPR codependency scores between each candidate gene and the 10 core NMD factor genes. This was juxtaposed with the mean CRISPR scores obtained between the same candidate gene and the set of random negative control genes, specific to the chromosomal regions under investigation (either 1q or 2q). Specifically, for the chromosome 1q region, linked to CNA-PC3, our assessment encompassed all 274 genes located within the 1q21.1-23.1 boundary, and were compared against 18 randomly selected control genes situated outside this specified region. Similarly, for the chromosome 2q region, associated with CNA-PC52, our analysis included all 303 genes found within the 2q31.1-2q36.3 area and were compared to 383 random control genes positioned outside the targeted region.

### Rare variant association studies (RVAS) via SKAT-O

We conducted a gene-based rare variant association study (RVAS) following the method from a previous study in our group^143^, utilizing the SKAT-O combined test^68^ (see schematic from Fig. 5A). Unlike burden testing, where variants are aggregated before regression against a phenotype, SKAT individually regresses variants within a gene against the phenotype and tests the variance of the distribution of individual variant score statistics. The SKAT-O test statistic is a weighted combination of the burden test (QB) and the SKAT variance test (QS) statistics, formulated as Qρ = ρ*QB + (1−ρ)QS. In this equation, Qρ represents the weighted mean statistic, and the parameter ρ determines the weighting of each test component. Notably, a gene burden test demonstrates greater power when all rare loss-of-function (pLoF) variants in a gene are causal, whereas SKAT exhibits higher power when some rare pLoF variants are not causal or when rare pLoF variants are causal but exert effects in opposing directions^68^.

We defined three stringency thresholds for determining putative pLoF for the rare germline variants i.e. those with a population allele frequency (MAF) of < 0.1%:

(i) pLoF: NMD-triggering PTC variants
(ii) pLoF_Missense_CADD15: NMD-triggering PTC variants + Missense with CADD >= 15
(iii) pLoF_Missense_CADD25: NMD-triggering PTC variants + Missense with CADD >= 25

The predicted deleterious missense variants were calculated using CADD scores^69^ at two different cutoffs: >= 25 (more stringent) and >= 15 (permissive).

Association testing was conducted separately for each cancer (33) or normal tissue type (56) and collectively for pan-cancer or pan-tissue analysis. We only considered individuals with European ancestry. A gene’s effect on iNMDeff was tested only if a rare pLoF variant in that gene was found in at least two individuals, across 18 matched cancer-normal tissues (Supp. Table S2). This resulted in varying numbers of genes being tested per cancer or tissue type. Significant associations are allowed to validate between GTex and TCGA cohorts across the different pLoF variant sets and NMD efficiency estimation method (ETG or ASE). In other words, as long as there is a significant association in matched tissues in both GTex and TCGA, regardless of the dataset or NMD method used, the association of the gene with NMD efficiency is considered replicated. The level of significance was determined after applying FDR correction using Benjamini-Hochberg within each tissue type separately in both cohorts (only correcting using the significant genes in the validation cohort). As a negative control, we randomized the iNMDeff values and re-tested the associations in the same way.

With this, we established an empirical FDR benchmark by contrasting the proportion of significant findings against those from randomized data. We calculated an empirical FDR of 30% at a nominal threshold of 5%, following this formula:

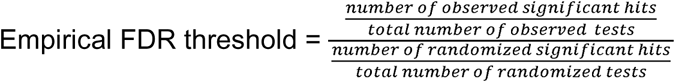

In mode detail, we performed SKAT-O testing using the R package “*SKAT v2.0.1*”. The initial step involved regressing covariates against iNMDeff using the function “*SKAT_Null_Model”.* When applicable, we controlled for age at diagnosis, sex, cancer (in TCGA) or tissue (in GTex) type, ancestry (first 6 PCs), RNA-seq sample library size, and tumor purity (in TCGA), as covariates. Categorical variables were encoded as dummy variables using “*fastDummies v1.6.3*”. Missing age data were imputed using the median value from the respective cohort. Post null model initialization, SKAT-O was executed with the “*SKAT*” function, setting the method to “*SKATO*”. This process involved running SKAT-O with 10 different p-values (ranging from 0 to 1) and identifying the p-value yielding the lowest p-value.

We only tested one type of inheritance model, encoding individual variants as follows:

- Additive model: 0 for no rare pLoF, 1 for rare pLoF, 2 for rare pLoF with somatic loss of heterozygosity (LOH, in TCGA) or biallelic rare pLoF.

### Gene-burden testing to estimate effect sizes for the significant RVAS genes

In addition to SKAT-O, we conducted gene-based burden testing to aggregate variants within the same gene, applying identical models as previously described. This was necessary because SKAT-O does not report effect sizes. The association testing was executed using linear regression with the *’glm’* function from that *‘stats’* base R package. The model was structured as follows:

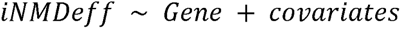

Here, the ‘Gene’ variable was treated as a binary categorical variable, representing the presence or absence of aggregated gene-level variants from the additive model. Covariates were incorporated as appropriate, including age at diagnosis, sex, cancer (in TCGA) or tissue type (in GTex), ancestry (first 6 PCs), RNA-seq sample library size, and tumor purity (in TCGA). Primarily, burden testing was employed to ascertain the effect sizes, i.e. the beta coefficients, of the 10 significant and replicated hits identified in our RVAS analysis. A positive effect size indicates that pLoF variants within the gene are associated with a higher iNMDeff and vice versa. *P*-values were adjusted by FDR within every gene and tissue separately, for every cohort.

### Redefinement of cancer gene list for estimating selection

Our initial selection of cancer-related genes originated from the 727 genes listed in the Cancer Gene Census (CGC)^58^. Within this list, we reclassified genes labeled as “Oncogene, Fusion” as Oncogenes (OGs), and similarly reclassified genes for tumor suppressor genes (TSGs). However, we excluded genes labeled as “oncogene, TSG” or “oncogene, TSG, fusion” due to the ambiguity in their classification. Additionally, we narrowed our focus to OGs with dominant inheritance patterns and excluded those specific to leukemia. To further refine our list, we cross-referenced it with two distinct gene sets:

1) The gene set identified by Solimini et al.^72^, who employed computational predictions and experimental validations through shRNA techniques. We intersected 1667 “STOP” genes with our 240 TSGs and 1249 “GO” genes with our 187 OGs.
2) The Mutpanning gene set^73^, where we focused on 41 genes with a cancer incidence higher than 3 and a significant q-value < 0.01, intersecting with our 240 TSGs. For our 187 OGs, we included all 217 genes with any cancer incidence and a significant q-value <0.01.

By combining the genes from these two intersections, we formed a more refined and specific gene set, which ultimately comprised 50 TSGs and 52 OGs. This integrative approach ensures a comprehensive and evidence-based selection of 102 key cancer genes.

### Estimation of selection in cancer genes via dNdS

The dNdS method is a state-of-the-art method to estimate selection, computing the dN/dS ratios for different types of mutations – missense, nonsense, and frameshift – at the gene level. A dN/dS ratio above 1 for a gene indicates positive selection, while a ratio below 1 signifies negative selection. dNdS was calculated using the “*dndscv*” function from the R package “*dndscv*” using the substitution model “*submod_192r_3w*“^71^.

We implemented this approach on all nonsense (stratifying NMD-triggering and NMD-evading PTCs) and missense mutations (serving as controls) found within the CGC set of genes in the pan-cancer TCGA cohort. We also applied this analysis for 33 cancer types separately, to identify cancer-specific genes. Individuals were grouped in iNMDeff high vs low, based on the median. In other words, our analysis was divided into various categories: i) comparison between NMD-triggering and NMD-evading PTCs; ii) comparison of individuals with high versus low iNMDeff; iii) analysis of the two types of somatic mutations, namely nonsense/PTCs and missense; iv) the two categories of cancer genes: TSG and OG. Consequently, each gene underwent dN/dS ratio calculation eight times, reflecting the above different contexts including mutation type, NMD region type, cancer gene type, and iNMDeff group sample type.

To enhance the accuracy of our classification between NMD-triggering and NMD-evading PTCs, we incorporated a new NMD-rule^8^ acknowledging that PTCs in longer exons might escape NMD. Accordingly, PTCs located in exons with a length >= 1000nt were categorized as NMD-evading. Conversely, PTCs considered NMD-triggering were restricted to exons with a maximum length of 500 nt.

### Survival analysis and cox regressions on OS and PFS

To investigate the relationship between individual NMD efficiency and cancer survival, we employed Cox proportional hazards regression models for each of the 47 cancer types stratified into subtypes and for the two NMD methods. These models were used to analyze two key survival outcomes: overall survival (OS) and progression-free survival (PFS). The structure of the model was as follows:

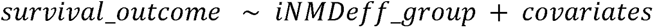

In this model, ‘*iNMDeff_group*’ was utilized as a binary categorical variable to distinguish between groups with high and low iNMDeff. This classification was based on a quantitative threshold determined by a specific percentile. In TCGA, we incorporated several covariates when applicable, which included sex, cancer type, RNA-seq sample library size, tumor purity, age (categorized into quartiles and treated as a discrete variable) and TMB. In the external validation datasets, and when applying Cox regressions using PFS as the outcome, the selection of covariates was adjusted based on data availability. For Carrol et. al and Motzer et. al datasets, we only could use age and sex. However, in the Liu et. al dataset, we were able to include sex and tumor purity as covariates. For Carrol et. al and Liu et. al, but not Motzer et. al, we additionally included TMB.

The Cox regression analysis was systematically repeated across a spectrum of percentiles (every 5th percentile up to the 50th), enabling an intricate evaluation of NMD efficiency’s impact on survival. For example, classifying individuals at the 20th percentile, is comparing the top 20% of individuals with the highest iNMDeff against the bottom 20% with the lowest iNMDeff. Meta-analysis p-values were calculated between the two NMD methods when possible, applying FDR correction per cancer type afterwards, within the range of percentiles and both NMD methods. For outlier associations, we set exclusion criteria for cases where confidence intervals exceeded 20, or the exponential hazard ratio coefficients were beyond the bounds of 15 and 0.001, or resulted in “Inf”. Additionally, associations were disqualified if regression analysis warnings indicated a failure to converge, specifically if the model “Ran out of iterations and did not converge” or a “Non-convergence warning detected” was issued. We performed a randomization of iNMDeff values and re-execution of Cox regressions to establish an empirical FDR benchmark by contrasting the proportion of significant findings against those from randomized data. With this, we calculated an empirical FDR of 16% at a nominal threshold of 5%, following this formula:

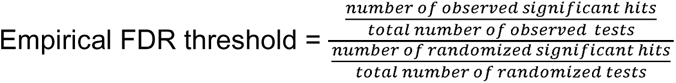

For our survival analysis, we utilized the *’surv’* and *’survfit’* functions from the *’survival’* R package. Kaplan-Meier curves were constructed using the *’ggsurvplot’* function from the *’survminer’* R package. Additionally, Cox proportional hazards regression models were executed employing the *’coxph’* function from the *’survival’* R package.

### Immunotherapy-treated patients from external datasets and filterings

To validate our PFS or immunotherapy response findings, we utilized external patient datasets from different studies, each focusing exclusively on individuals undergoing different immunotherapy treatments:

1) Liu et al. 2019^77^: 144 melanoma (SKCM) patients treated with anti-PD1 immune checkpoint inhibitors were examined. We normalized their RNA-seq gene-level TPM data using log2(TPM+1), excluding patients who received more than one previous therapy.
2) Carroll et al. 2023^78^: This research involved 35 patients with inoperable esophageal adenocarcinoma (EAC) who received first-line immunochemotherapy, initiating with ICI for four weeks followed by chemotherapy. RNA-seq gene-level raw counts were obtained, transformed to TPM using “*convertCounts*” function from “*DGEobj.utils*” R package, and normalized log2(TPM+1). Only pre-treatment (“PreTx”) RNA-seq samples were considered.
3) Motzer et al. 2020^79^: This dataset comprised 886 patients with advanced renal cell carcinoma (RCC), treated with the ICI avelumab plus axitinib or only sunitinib. The RNA-seq gene-level log2(TPM) counts were obtained for these patients. It is important to note that in the original data, values below 0.01 were set to 0.01, thus eliminating the need for pseudocount adjustments in our analysis. Samples receiving only sunitinib were omitted as it is a targeted therapy and not an immunotherapy.
4) Hartwig Medical Foundation^80^: this study included RNA-seq data from 3020 metastatic samples across 22 cancer types. We focused on 251 samples treated with immunotherapy (variable “consolidatedTreatmentType” == “immunotherapy”), normalizing their RNA-seq gene-level TPM counts using log2(TPM+1).
5) Riaz et al.^81^. Tumors from 68 patients with advanced melanoma (SKCM), who progressed on ipilimumab or were ipilimumab-naive, before and after nivolumab initiation. We accessed the RNA-seq gene-level raw counts (UCSC hg19) and transformed their TXIDs into Ensembl Gene IDs using “*biomaRt*”. After converting the raw counts to TPM and normalizing via log2(TPM+1), we retained only the samples under treatment (“On”), excluding pre-treatment (“Pre”) cases due to their scarcity.
6) TCGA: 469 melanoma (SKCM) samples, 73 were identified as having received at least one immunotherapy treatment.

The iNMDeff for these datasets was predicted using the obtained RNA-seq data, with TCGA or GTex cohorts serving as training sets. Averages of predictions from both cohorts were taken as the final iNMDeff for each individual, calculated separately for ETG and ASE methods. This approach yielded two iNMDeff values per individual. The final sample sizes, after filtering, were 120 for Liu-SKCM, 33 for Carroll-EAC, 353 for Motzer-SKCM, 251 for Hartwig, 48 for Riaz-SKCM, and for TCGA-SKCM, we utilized directly our estimated iNMDeff values for 36 and 73 ASE and ETG melanoma patients, respectively.

### Prediction model of immunotherapy response

To predict immunotherapy response, we designed a simple logistic regression model incorporating TMB and our predicted iNMDeff to classify immunotherapy outcomes as “Responders” (Complete Response, CR, and Partial Response, PR) or “Non-Responders” (Stable Disease, SD, and Progressive Disease, PD). This analysis spanned various datasets, excluding Motzer-SKCM due to unavailability of TMB and treatment response data, and included TCGA-SKCM, Carrol-EAC, Liu-SKCM, and Hartwig (categorized by urinary tract, lung, and melanoma), as well as Riaz-SKCM. The approach to TMB calculation and response categorization was dataset-specific and manually curated:

- TCGA SKCM (n = 41): Patients with at least one immunotherapy treatment were identified, with the best response recorded as the outcome. Patients with “NA” in response to treatment were excluded. Clinical PD or SD were tagged as “Non-Responders”, whereas PR or CR as “Responders”. TMB was computed as the total count of somatic coding mutations per individual, normalized by the exome size (36 Mb).
- Liu-SKCM (n = 118): Used the “BR” variable, excluding “MR” responses, and classified the remaining as per standard criteria. The “Total_muts” variable served as TMB.
- Carrol-EAC (n = 33): Applied “Response_binary” directly for classification into “Responders” and “Non-Responders”. Only pre-treatment (PreTx) iNMDeff-estimated samples were considered, with “PreTx.Tumor_Mutational_Load” as TMB.
- Riaz-SKCM (n = 48): Directly used the “Response” variable, including only patients treated “On” treatment, utilizing “Mutation.Load” for TMB.
- Hartwig (n = 251): “firstResponse” was filtered to exclude “ND” and “MR”, with “CR” and “PR” classified as Responders; “PD”, “Non-CR/Non-PD”, “Clinical progression”, “SD” as “Non-Responders”, stratified by cancer type for adequately sized samples (discarded if n < 20): Hartwig-UT (urinary tract, n = 23), Hartwig-lung (n = 26), and Hartwig-SKCM (melanoma, n = 119). “TOTAL_SNV” was taken as TMB.

Covariate adjustments were made based on dataset characteristics, incorporating age and sex generally, with Liu-SKCM excluding age but including cancer subtype due to significant TMB differences across melanoma subtypes, and Riaz-SKCM only including cancer subtype.

The “*glm*” function from R’s stats package, set with “binomial” family parameter, was employed for the logistic regression. Model efficacy was gauged using “AUC” and “roc” from the “pROC” R package, contrasting AUC values against a randomized baseline (∼0.5 AUC) via “roc.test” for significance evaluation. Moreover, sensitivity (recall), specificity, and precision were calculated using the “*caret*” R package, providing a comprehensive evaluation of the model’s performance across various metrics.

## 5. Acknowledgements

We are grateful to dr. Javier Lanillos for processing the TCGA RNASeq data in brain tumors using UniCell Deconvolve, and for pointing us to immunotherapy studies of interest. Further, we thank all members of the Genome Data Science lab for discussions. G.P.-M. was funded by an AGAUR FI fellowship, and F.S. was funded by the ICREA Research Professor program. Work in the lab of F.S. was supported by an ERC StG “HYPER-INSIGHT” (757700), Horizon2020 project “DECIDER” (965193), Spanish government project “REPAIRSCAPE”, CaixaResearch project “POTENT-IMMUNO” (HR22-00402), the SGR funding of the Catalan government, and the Severo Ochoa Centers of Excellence award of the Spanish government to the hosting institution. This publication and the underlying research are partly facilitated by Hartwig Medical Foundation and the Center for Personalized Cancer Treatment (CPCT) which have generated, analysed and made available data for this research. The results published here are in whole or part based on data generated by the TCGA Research Network (https://www.cancer.gov/tcga).

## Notes

### Competing Interest Statement

The authors have declared no competing interest.

