## Supplementary Figures S1-S21 and Supplementary Table S1-S8 legends for "Variable efficiency of nonsense-mediated mRNA decay across human tissues, tumors and individuals"

### Supplementary Material

**Supp. Fig. S1**

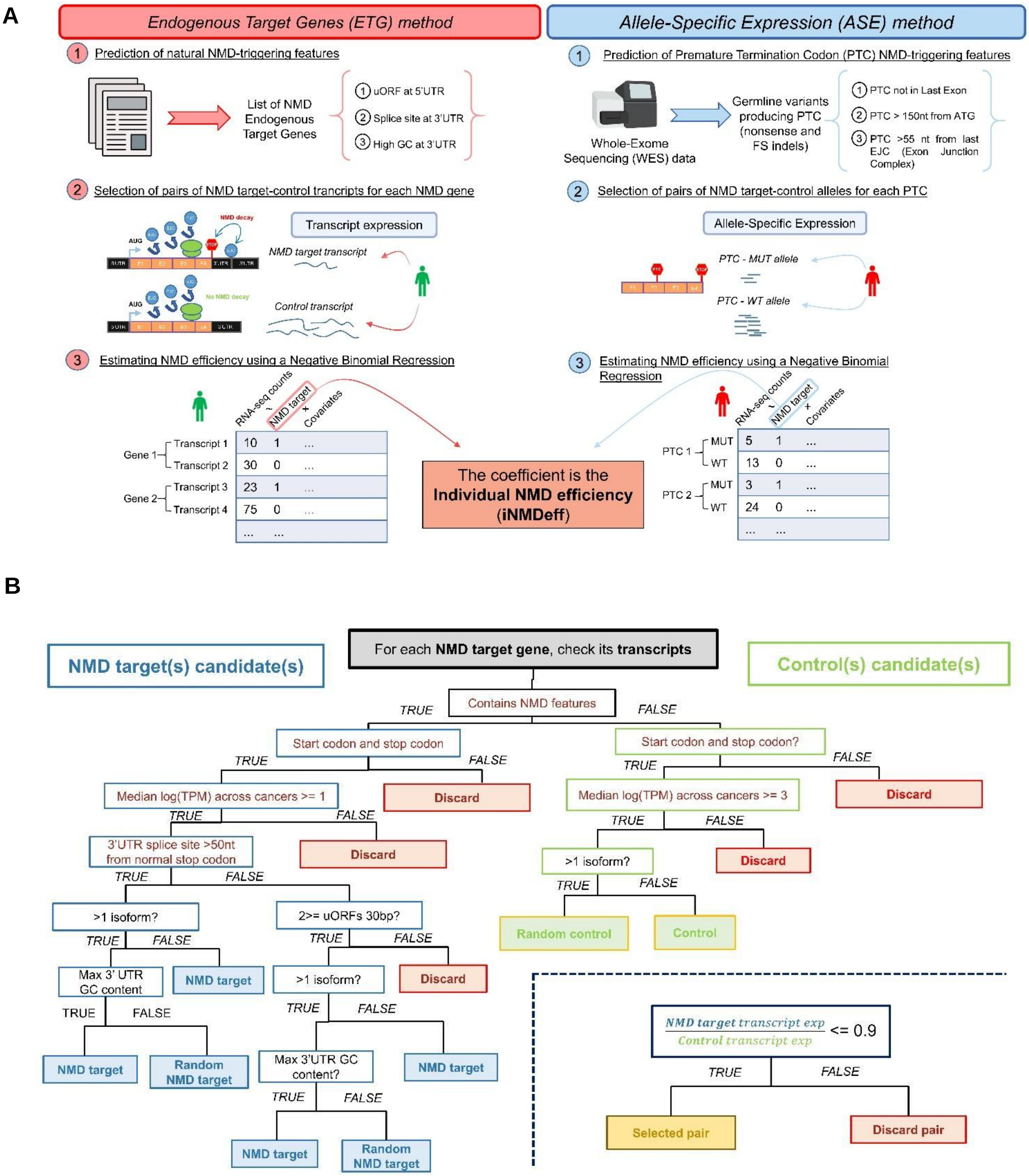

**Supp. Fig S1. Overview of NMD methods to estimate individual NMD efficiency (iNMDeff)**

**A,** Schematic illustration of two distinct methodologies employed to estimate individual NMD efficiency (iNMDeff): the Endogenous Target Gene (ETG) NMD method, on the left, and the Allele-Specific Expression (ASE) NMD method, on the right. Each method provides a unique approach to quantify the effectiveness of the NMD pathway in degrading mRNA transcripts for each individual, either using germline premature termination codons (PTCs) or endogenous transcripts with NMD-triggering features. **B**, Methodology for pairing NMD target and control transcripts for the ETG method. Decision tree outlining the process for selecting a pair of NMD target and control transcripts for each gene within the various NMD gene sets for the ETG method. The left branch of the tree delineates criteria for identifying NMD target candidates based on intrinsic NMD-triggering features: i) a splice site in the 3' untranslated region (UTR) situated at least 50 nt downstream from the normal stop codon; ii) an upstream open reading frame (uORF) within the 5' UTR; iii) high GC content within the 3'UTR. If multiple transcripts per gene exhibit these features, one is chosen at random. The right branch of the tree defines the selection of control candidates, which must exhibit some level of gene expression across cancer types without any NMD-triggering features. After pairing an NMD target with a control, the expression ratio must be less than 0.9 to be valid. If this criterion is not met, the selection process is reiterated to identify an alternative pair.

**Supp. Fig. S2**

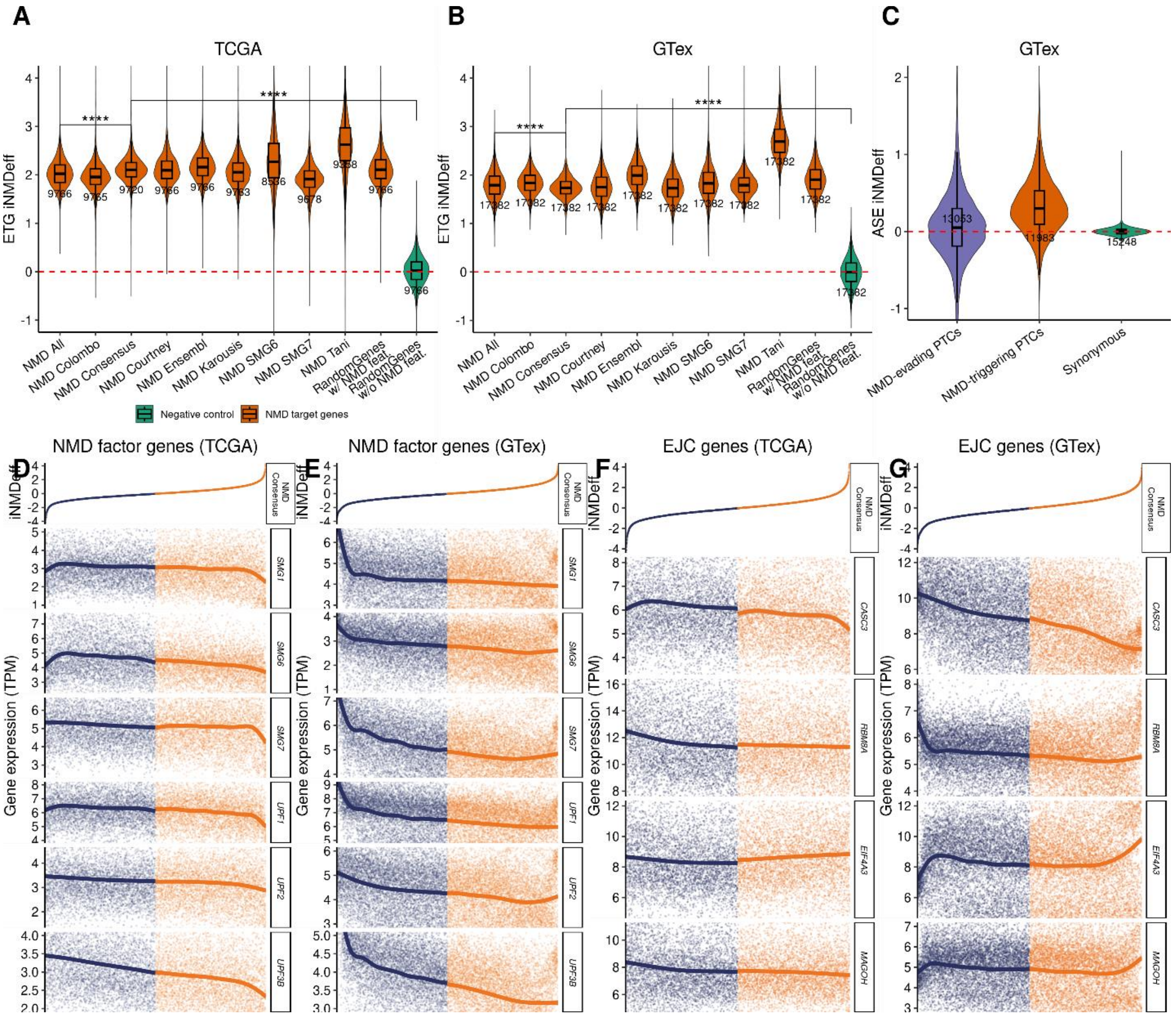


**Supp. Fig. S2. Individual-level quantification of NMD efficiency.**

**A-B**, Estimation of individual NMD efficiency, iNMDeff (Y-axis), using the ETG method across 9,766 TCGA samples (A) or 17,382 GTex samples (B), showcasing all NMD gene sets, one random gene set with NMD-triggering features, and one random gene set without NMD-triggering features as negative control. **C**, iNMDeff estimations using the ASE method, in GTex, for two NMD variant sets, alongside a non-NMD variant set as a control. ****$p < 0.0001$, by two-sided Mann–Whitney $U$ test for A and B. **D-E,** Gene expression levels (TPM) of 6 core NMD factors (*SMG1*, *SMG6*, *SMG7*, *UPF1*, *UPF3A*, *UPF3B*) compared against the ETG iNMDeff (using the “NMD Consensus” gene set) sorted from lowest to highest along the X-axis in TCGA (D) and GTex (E). Samples are stratified by the median ETG iNMDeff in high and low. **F-G**, Same as in D-E but for 4 core EJC genes (*CASC3*, *RBM8A*, *EIF4A3*, *MAGOH*) in TCGA (F) and GTex (G).

**Supp. Fig. S3**

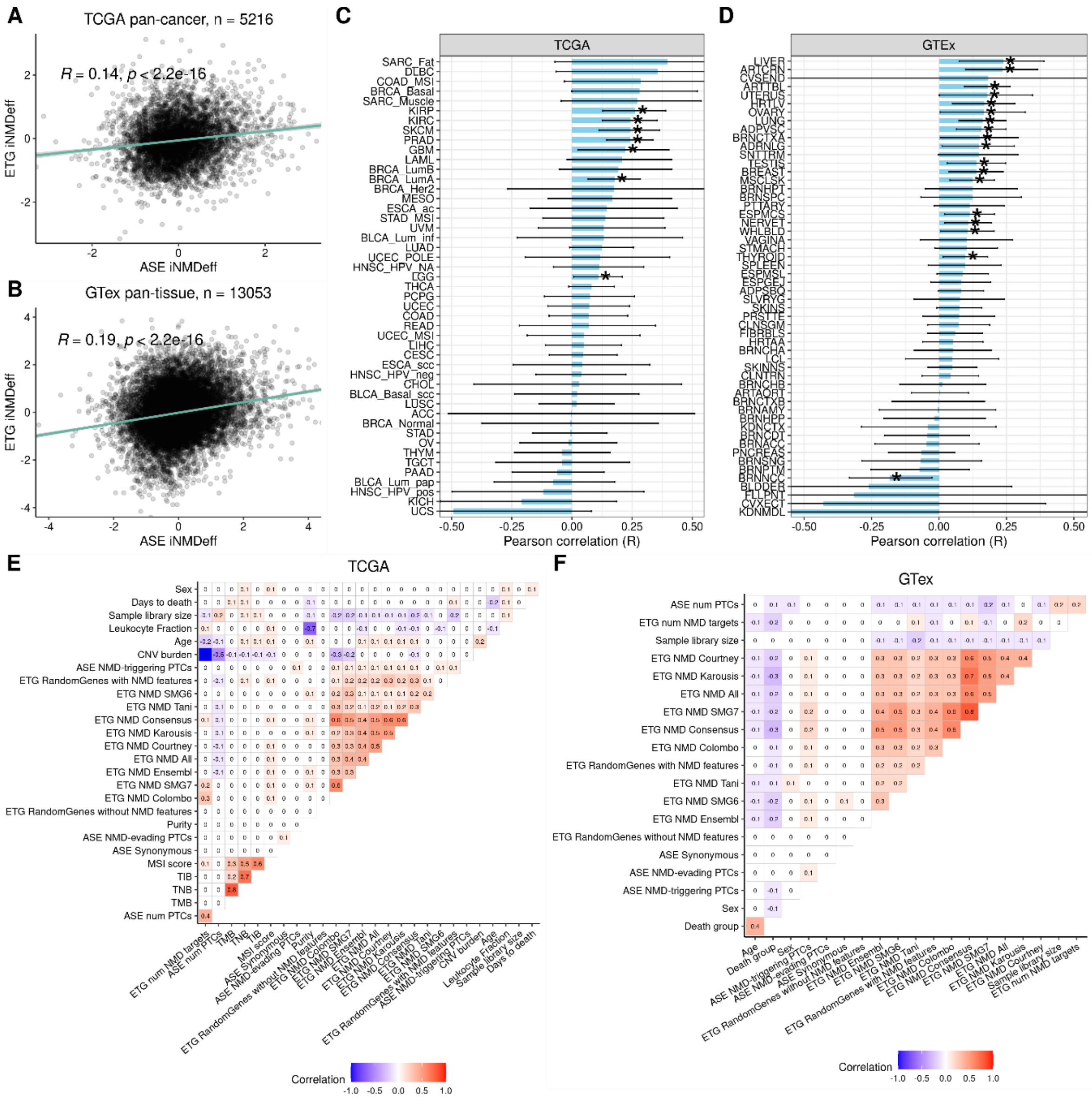


**Supp. Fig. S3. Agreement between the two methods to estimate NMD efficiency**

**A-B,** The correlation between ETG iNMDeff (Y-axis) and ASE iNMDeff (X-axis) is evaluated across pan-cancer samples in TCGA (A) and pan-tissue samples in GTex (B). Due to missing ASE data, the number of samples analyzed in TCGA is 5216 and 13053 in GTex. Each plot displays the Pearson correlation coefficient (R) and associated *p*-value. **C-D**, Similar to A-B, but the analysis is stratified for each cancer type (or subtype) in TCGA (C) and each normal tissue in GTex (D), plotted on the Y-axis. The X-axis shows the Pearson correlation coefficient (R), with the range limited to between -0.5 and 0.5 for clarity. Correlations deemed significant (FDR < 5%) are marked with an asterisk (*). **E-F**, Correlation matrix between various biological and technical variables and iNMDeff estimates from both ASE and ETG methods for TCGA (E) and GTex (F) cohorts. Biological variables include: sex, leukocyte fraction, days to death, death group, age, copy number alteration (CNA) burden, microsatellite-instability (MSI) score, tumor indel burden (TIB), tumor mutation burden (TMB), tumor nonsense burden (TNB), number of PTCs used for the ASE method (ASE num PTCs). Technical variables include: RNA-seq sample library size and tumor purity. ETG iNMDeff estimates from 11 NMD gene sets, including its 2 non-NMD controls, are: ETG NMD SMG6, ETG NMD SMG7, ETG NMD Tani, ETG NMD Consensus, ETG NMD Karousis, ETG NMD Courtney, ETG NMD All, ETG NMD Ensembl, ETG NMD Colombo, ETG NMD RandomGenes without NMD features, ETG NMD RandomGenes with NMD features. ASE iNMDeff estimates from 3 NMD variant sets, including its controls are: ASE NMD-triggering PTCs, ASE NMD-evading PTCs and ASE Synonymous. The Pearson correlation coefficient (R) is represented by a color spectrum ranging from -1 (blue) for a perfect negative correlation to 1 (red) for a perfect positive correlation with iNMDeff.

#### Supp. Fig. S4

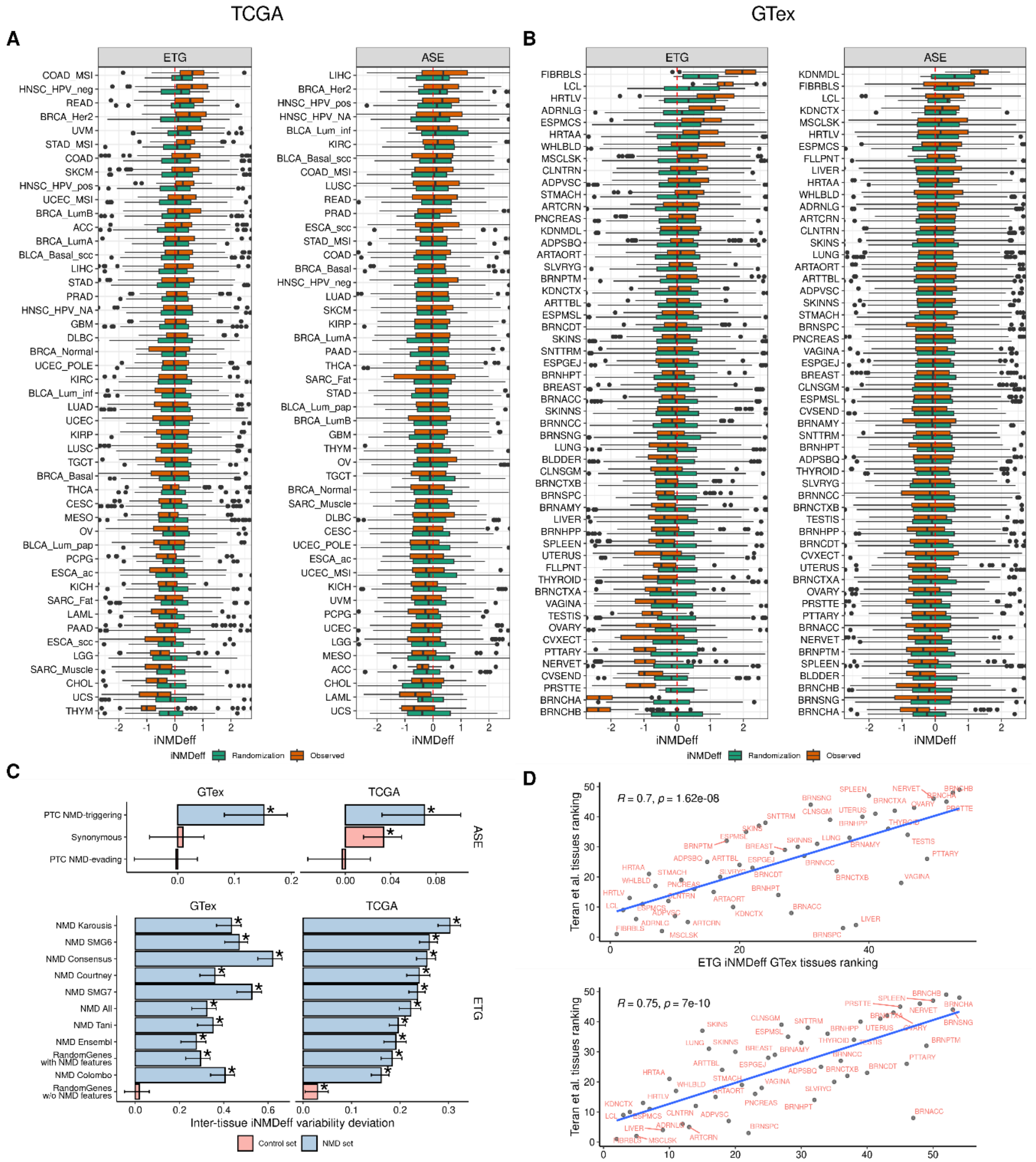


**Supp. Fig. S4. Inter-tissue variability of NMD efficiency**

**A-B**, Distribution of iNMDeff for both ETG (left panel) and ASE (right panel) methods across cancer types in TCGA (A) and normal tissues in GTex (B) are displayed on the X-axis as orange boxplots. Corresponding randomized iNMDeff values are illustrated alongside as green boxplots. **C**, The Inter-tissue iNMDeff Variability Deviation (ITNVD) test scores (X-axis) are shown for all evaluated NMD gene sets (11 for ETG method, 3 for ASE method), indicating the extent of variability in iNMDeff. For contrast, scores for non-NMD control gene sets are also presented.

Results from both GTex (left) and TCGA (right) cohorts are included. Positive ITNVD scores suggest significant variability in iNMDeff among different tissues or cancers, with statistical significance ($p \leq 0.05$) denoted by stars (*), as determined by the randomization test. **D**, Correlations of the ranking of tissues based on the median ETG iNMDeff (top panel) and ASE iNMDeff (bottom panel) from this study with the tissue ranking derived from Teran et al.'s ASE PTCs-NMDeff methodology. The analysis provides Pearson correlation coefficients (R) and corresponding *p*-values, quantifying the agreement between the two independent tissue ranking approaches.

#### Supp. Fig. S5

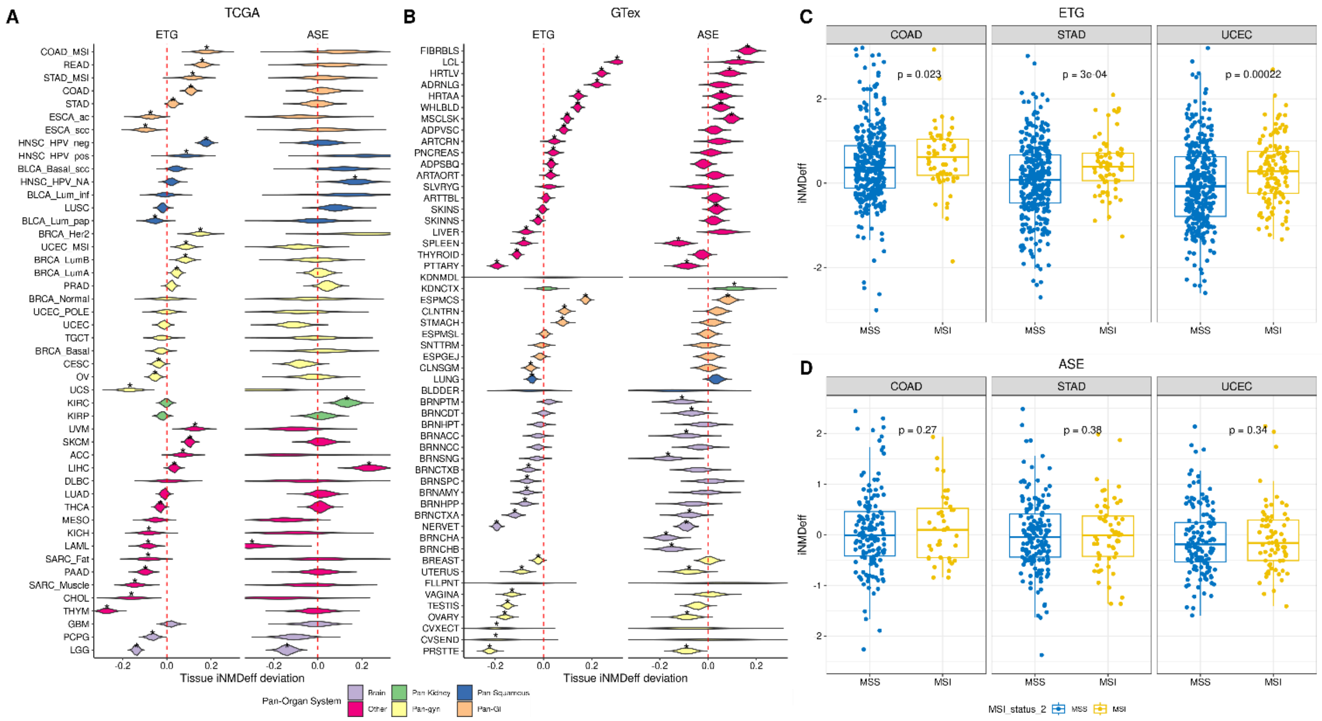


**Supp. Fig. S5. Intra-tissue variability of NMD efficiency**

**A-B**, Shows Tissue iNMDeff Deviation (TND) test scores (X-axis) across various cancer types in TCGA (A) and normal tissues in GTex (B), providing insight into the specific variations of NMD efficiency within tissues. Tissues are classified in primary groups: Nervous system-related tissues (Pan-nervous), Kidney-related tissues (Pan-kidney), Reproductive system tissues (Pan-reproductive), Gastrointestinal tissues (Pan-GI), those originating from Squamous cells (Pan-squamous), and the remaining tissues (Other). The groups of tissues were ordered based on the median TND scores, arranging them from the highest to the lowest median scores, top to bottom. **C-D**, Displays the iNMDeff on the Y-axis for both ETG (C) and ASE (D) methods in TCGA cancers: COAD, UCEC, and STAD, further dividing samples into MSI and microsatellite stable (MSS) groups. One-sided Mann-Whitney *U* tests were used to calculate *p*-values.

**Supp. Fig. S6**

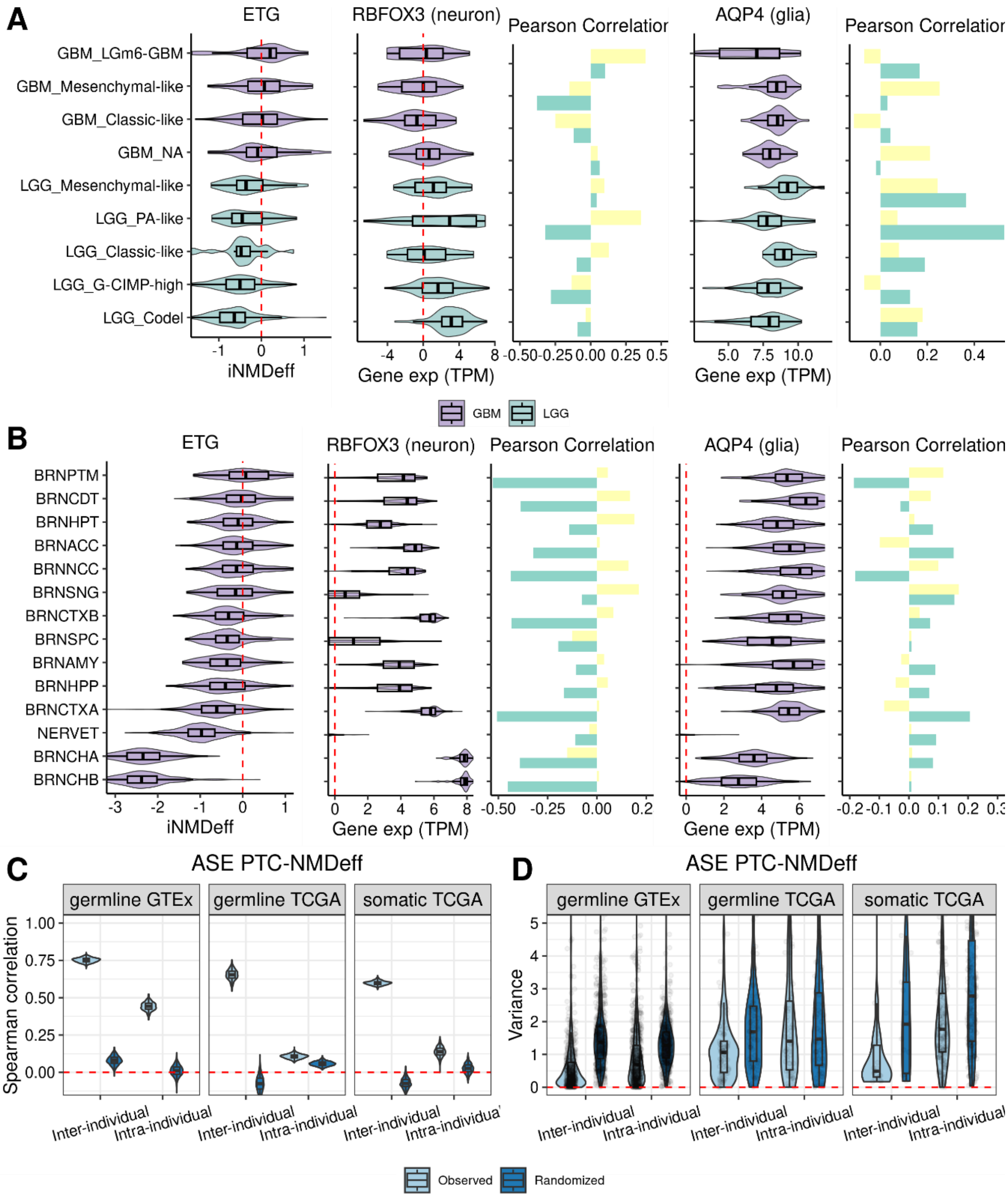


**Supp. Fig. S6. Lower NMD efficiency in the tissues of the nervous system and extensive inter-individual variation of NMD efficiency**

**A,** Scaled ETG iNMDeff (X-axis) for LGG and GBM within the TCGA cohort, segmented by genetic or histological subtypes. This section also shows the scaled gene expression (TPM) of the neural marker gene *RBFOX3* (second panel), along with its Pearson correlation with iNMDeff for both ASE and ETG methods (third panel). The analysis is extended to the glial gene marker *AQP4* in the fourth and fifth panels, illustrating expression levels and correlation with iNMDeff, respectively. **B,** Mirrors the analysis in A but focuses on GTex brain tissues, stratified by different subregions, showcasing the variability of NMD efficiency in normal brain tissue contexts and its correlation with neuron or glial cell type gene expression markers. **C-D**, Delve into the intra- and inter-individual variability of iNMDeff, leveraging ASE PTC-NMDeff. C assesses variability through the Spearman correlation of randomly selected PTC pairs within individuals (intra-individual variability) and between pairs of randomly chosen individuals sharing the same PTC (inter-individual variability). D repeats this comparative analysis using variance instead of correlation to measure variability. Light blue boxplots represent observed values, while dark blue boxplots serve as randomization controls, providing a baseline for comparison. These analyses are conducted within the TCGA cohort for both germline and somatic PTCs and extended to germline PTCs within the GTex cohort.

**Supp. Fig. S7**

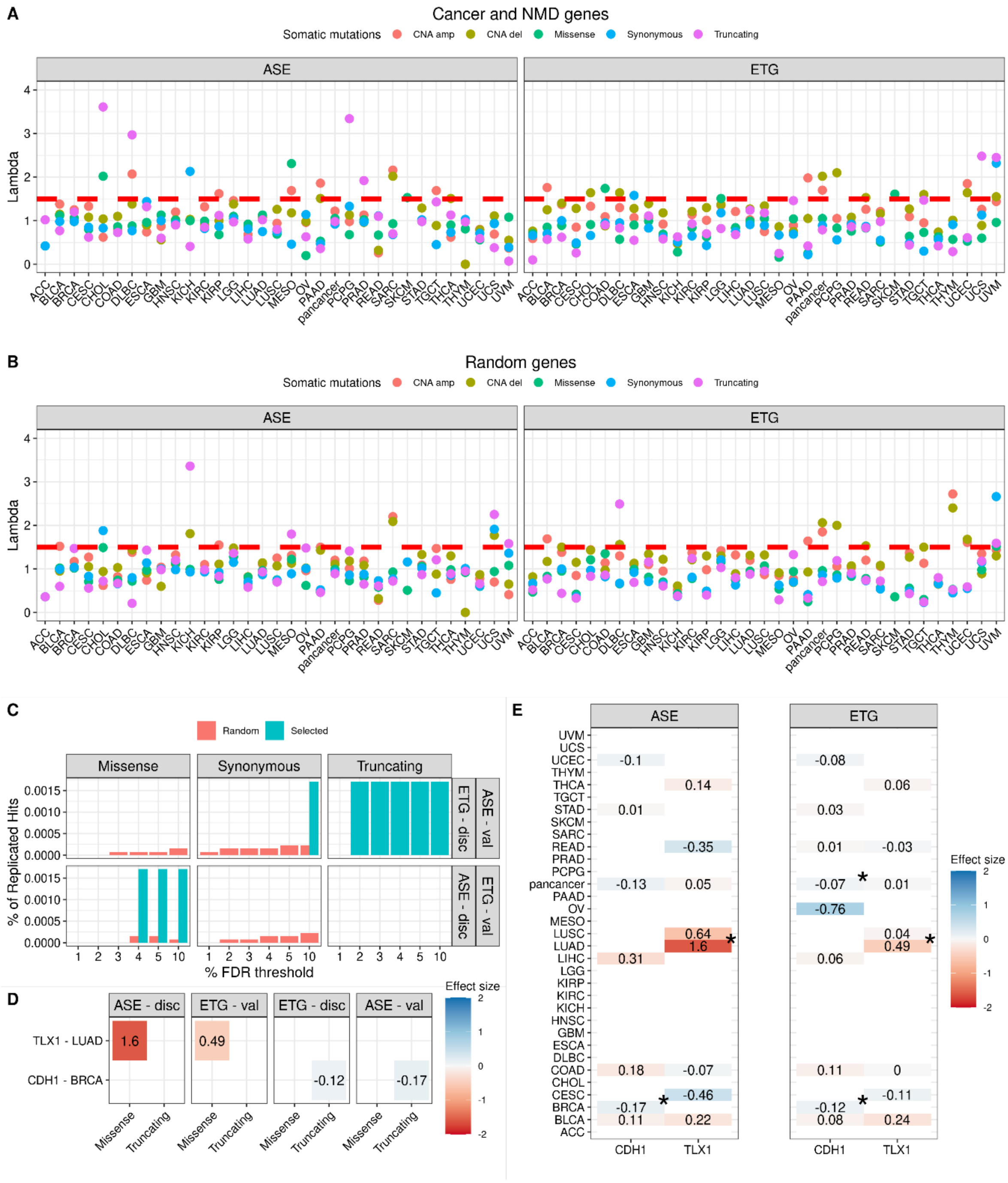


**Supp. Fig. S7. Associations analysis of somatic mutations with NMD efficiency.**
**A**, Visualizes inflation factor values, denoted as lambda (Y-axis), reflecting the degree of inflation in linear association studies between iNMDeff and various types of somatic mutations within cancer genes (TSGs and OGs) and NMD-related genes. Mutation types include Missense, Synonymous, Truncating, CNA amplifications, and CNA deletions. Analysis spans pan-cancer and individual cancer types (X-axis) for ASE (left panel) and ETG (right panel) methods. An

horizontal dashed-red line marks a lambda threshold (>= 1.5), beyond which associations are considered excessively inflated and thus excluded from further analysis. **B**, Replicates the lambda figure from A but focuses exclusively on the remaining genes, effectively filtering out cancer and NMD genes to assess inflation in a control random gene set. **C**, Proportion of replicated hits—calculated as the number of hits over the total number of tests—for each mutation type, depicted across different FDR thresholds from 1% to 10% (X-axis). The analysis alternates between ASE and ETG methods for discovery and validation within the same cancer type, illustrated in two panels: ETG discovery to ASE validation (top) and vice versa (bottom). Blue bars represent observed hits in cancer/NMD genes; red bars correspond to hits among the remaining genes. CNA amplifications/deletions were ultimately excluded due to significant inflation. **D**, Lists replicated and significant genes (Y-axis), categorized by somatic mutation type (X-axis), and stratified by NMD method (ASE or ETG) and direction of discovery/validation. Effect sizes, shown as beta coefficients from linear model associations with iNMDeff, are color-graded: negative values in red and positive in blue. **E**, Compiles effect sizes of associations for all cancer types in TCGA (Y-axis) for the two significant and replicated genes (X-axis), using the same color gradient as in D for effect size visualization. Cancer-type specific associations marked with significance ($p < 0.05$) are highlighted with an asterisk (*).

**Supp. Fig. S8**

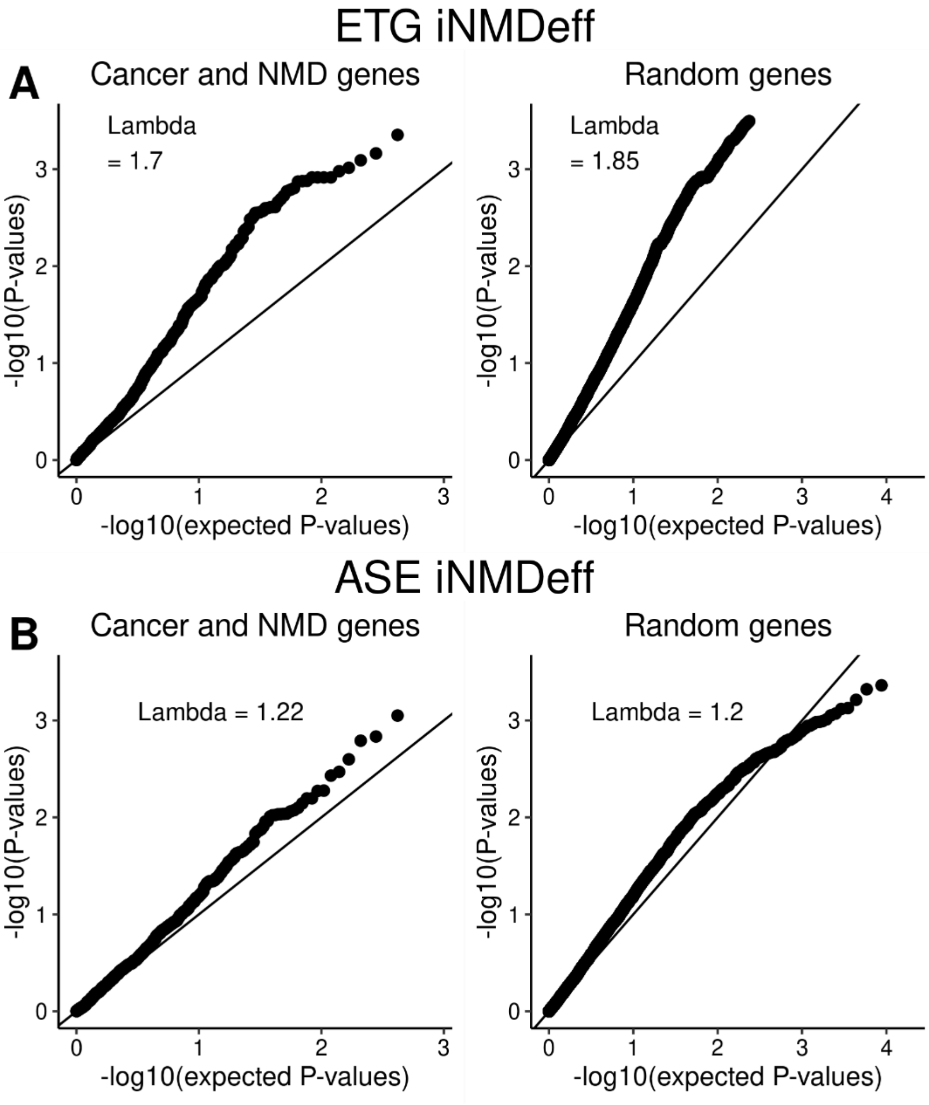


**Supp. Fig. S8. Inflation factor of the associations between somatic gene-level CNA amplifications and iNMDeff.**

**A-B,** QQ-plots examining the inflation in linear association studies between somatic gene-level CNA amplifications and iNMDeff, utilizing both ETG (A) and ASE (B) methods at the pan-cancer level. The analysis divides into associations within cancer and NMD genes (left panel) versus the remaining genes (right panel), with the Y-axis depicting the observed p-values and the X-axis the expected p-values, both in -log10 scale, to highlight deviation from expectation. Notably, inflation factor values, lambda, are provided, indicating significant inflation, particularly in the ETG method, which emphasizes the challenges in interpreting these associations.

**Supp. Fig. S9**

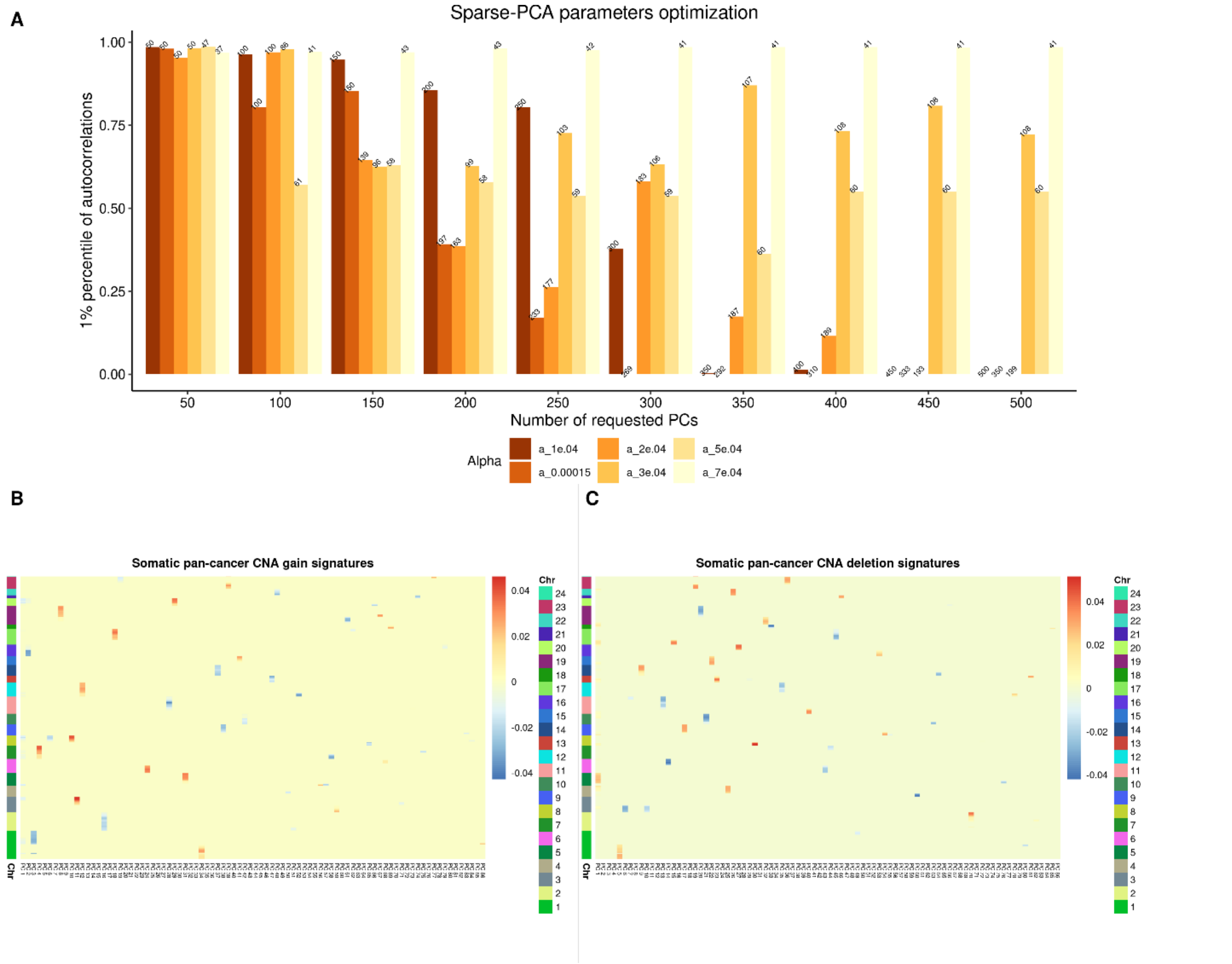


**Supp. Fig. S9. Refinement of sparse principal component analysis (sparse-PCA) parameters to effectively capture gene-level CNAs.**

**A**, Illustrates the the number of principal components (PCs) derived from sparse-PCA, capturing CNA gains or losses, against autocorrelation scores for the 1% of PCs with the lowest autocorrelation. The Y-axis tracks autocorrelation values from 0 to 1 for these PCs, while the X-axis varies the number of requested PCs in the *sparsepca* function, ranging from 50 to 500. Different alpha parameters (from 7e-04 in red to 1e-04 in light yellow) dictate the color gradient, indicating the degree of sparsity. Atop each bar, the count of "effective" PCs—those not entirely composed of zeros in gene weights—is noted. Optimal parameters were identified as requesting 100 PCs, yielding 86 non-zero effective PCs, with an average autocorrelation of 0.99 among the top 1% of PCs. **B-C**, Heatmaps displaying the gene weights across all 86 CNA-PCs, from -0.04 (blue) to 0.04 (red), sorted by genomic and chromosomal location, with each chromosome distinctly colored (legend provided). Chromosome 23 is marked as X, and Chromosome 24 as Y. In B (for CNA amplifications) and C (for CNA deletions), the initial 47 CNA-PCs predominantly indicate broad-scale alterations, akin to arm-level changes, while the remaining PCs highlight more localized genomic events.

**Supp. Fig. S10**

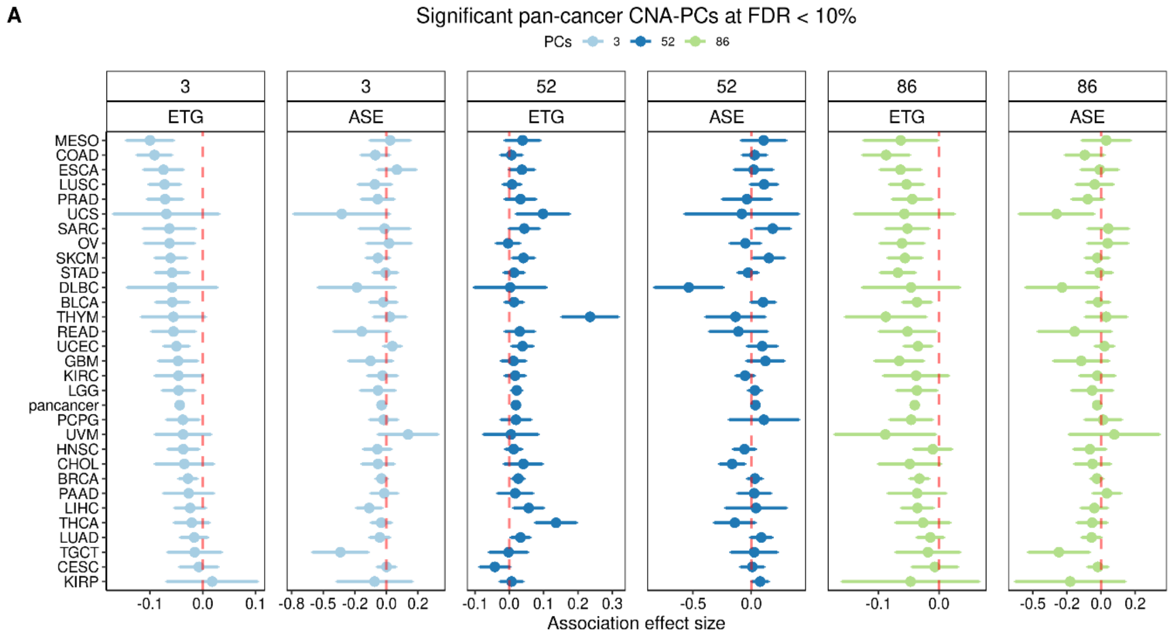


**Supp. Fig. S10. Significant associations between pan-cancer CNA-PCs and iNMDeff in TCGA at FDR < 10%**

Significant associations between three pan-cancer CNA-PCs and iNMDeff, employing both ETG (left panels) and ASE (right panels) methodologies, under a 10% FDR threshold. The implicated CNA-PCs are CNA-PC 3, CNA-PC 52, and CNA-PC 86, each demonstrating a noteworthy linkage with iNMDeff. The Y-axis categorizes the cancer types, including the pan-cancer level, while the X-axis quantifies the effect size, represented by the beta coefficient from linear model associations. The effect size means that higher values of a given CNA-PC signature correlate with increased iNMDeff, and vice versa.

**Supp. Fig. S11**

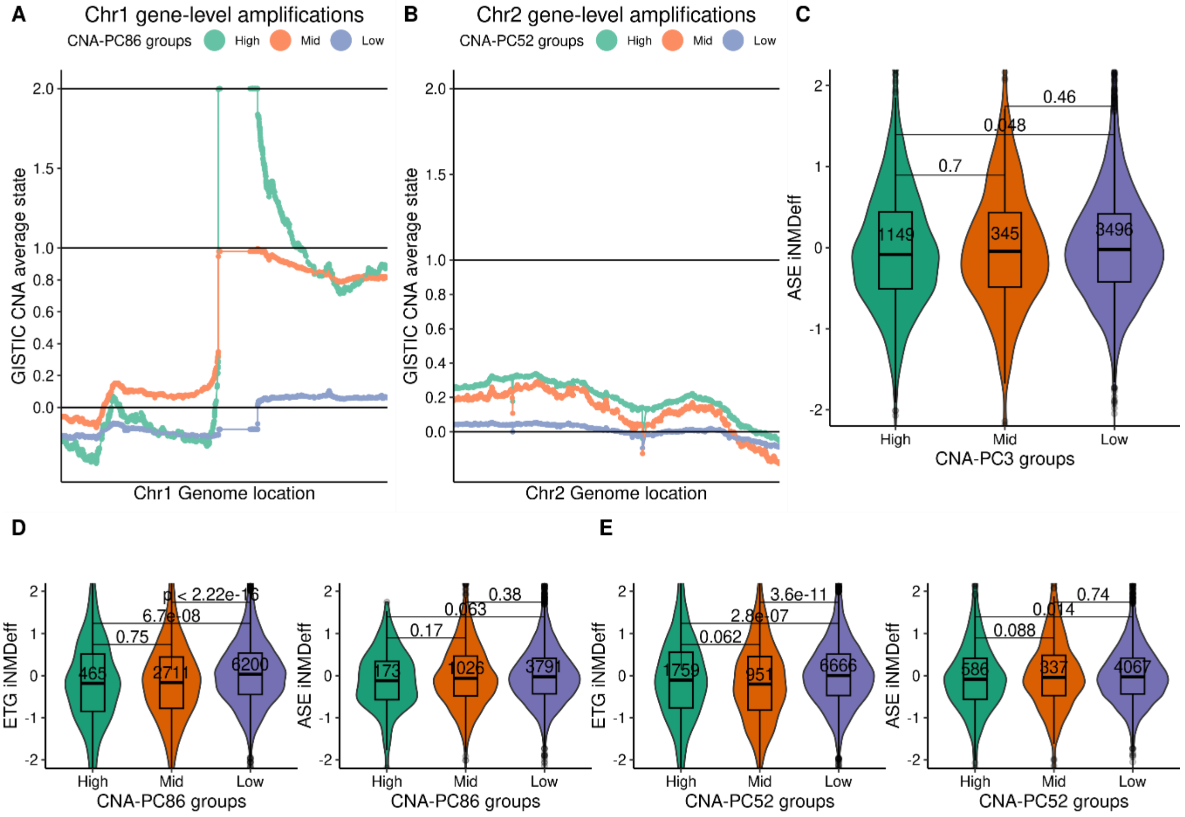


**Supp. Fig. S11. Somatic chromosomes 1q and 2q gains associate with reduced NMD efficiency**

**A-B**, Gene amplifications across chromosomes 1 (A) and 2 (B), ordered by genome location, are plotted along the X-axis, with amplifications assessed by averaging GISTIC CNA scores for each gene across TCGA participants (Y-axis). Samples are classified into three groups based on their pan-cancer CNA signature scores: "High", "Mid", and "Low" for both chromosomes, corresponding to CNA-PC3 for chr1 and CNA-PC52 for chr2. **C-E**, Stratification of scaled iNMDeff (Y-axis) according to these CNA-PC groupings: "High," "Mid," and "Low," across the X-axis for CNA-PC3 (C, specifically for ASE iNMDeff), CNA-PC86 (D, showcasing ETG iNMDeff on the left and ASE iNMDeff on the right), and CNA-PC52 (E, similarly displaying ETG and ASE iNMDeff). Statistical comparisons between groups utilize a two-sided Mann-Whitney *U* test to obtain the *p*-value.

**Supp. Fig. S12**

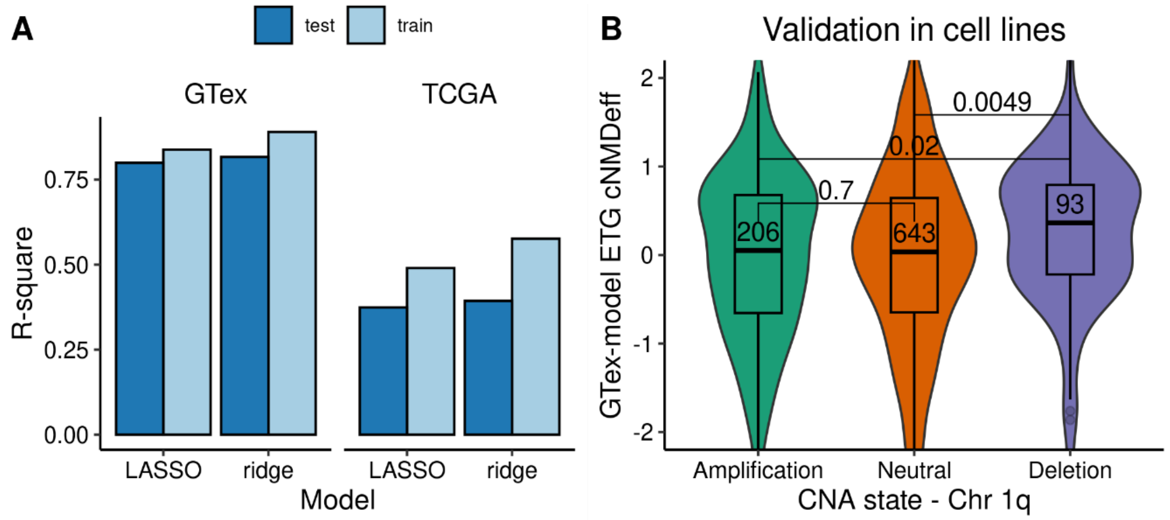


**Supp. Fig. S12. Proxy-model of ETG iNMDeff using global gene-level expression data for validations in external datasets.**

**A**, Performance of the models to predict ETG iNMDeff using gene expression data. Y-axis illustrates R-square values for LASSO and ridge regression models (X-axis). These models are separately trained and tested within both GTex (left panel) and TCGA (right panel) cohorts, demonstrating the models' predictive performance (R-square). **B**, Validation of the impact of chromosome 1q amplification on reducing iNMDeff. Here, a proxy-model, trained on GTex gene-level expression data, is applied to estimate cell line NMD efficiency (cNMDeff, Y-axis), categorized by their 1q arm status: amplified, neutral, or deleted. The groups' comparisons, conducted through a two-sided Mann-Whitney *U* test, underscore the statistical significance ($p < 0.05$) of 1q amplification's effect on diminishing NMD efficiency.

#### Supp. Fig. S13

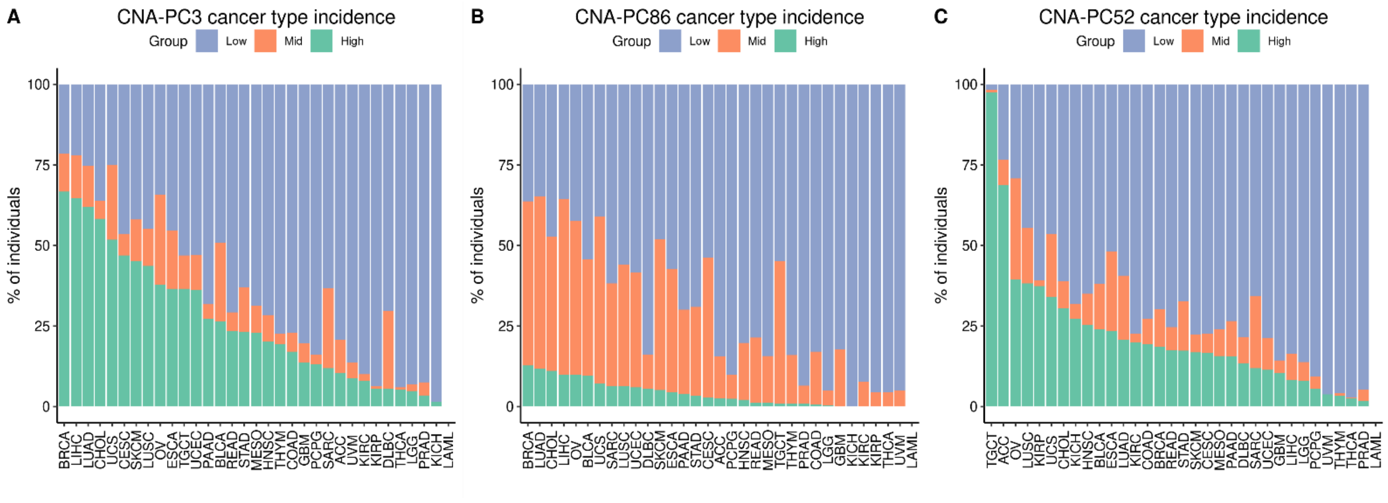


**Supp. Fig. S13. Incidence of CNA-PCs 3, 52 and 86 across cancer types.**
**A-C**, Percentage of individuals within each cancer type (X-axis), excluding LAML, categorized by their CNA-PC group status: “Low”, “Mid”, and “High”. Panels A to C respectively showcase the cancer incidence distributions for CNA-PC 3, 86, and 52.

#### Supp. Fig. S14

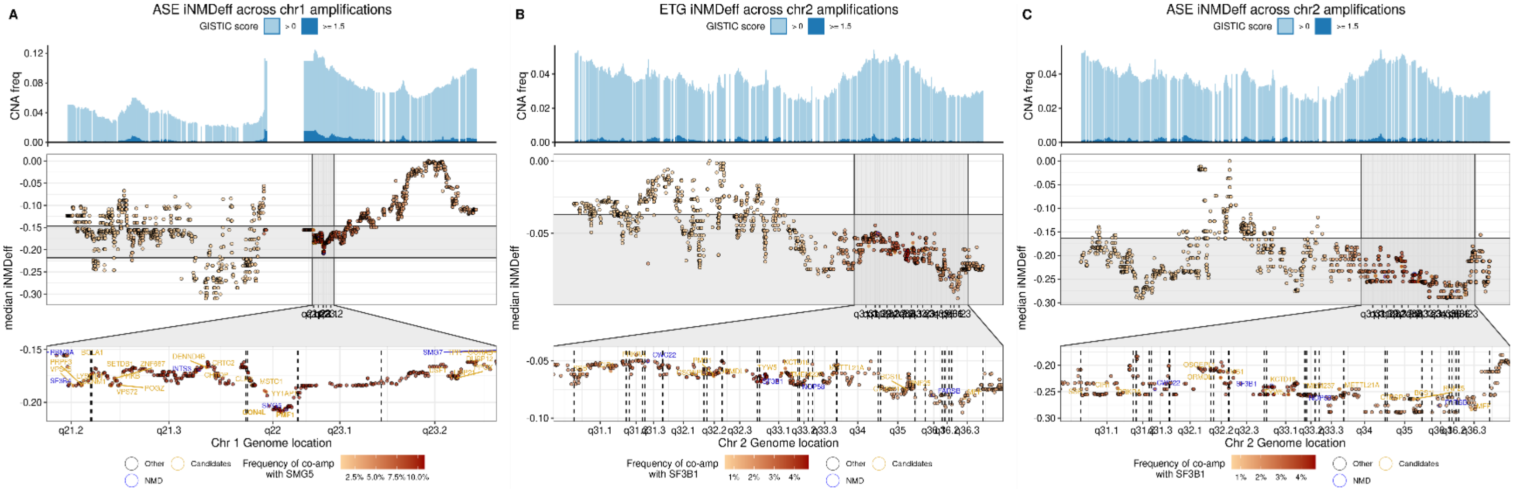


**Supp. Fig. S14. Gene-level iNMDeff showcases how focal CNAs along chromosomes 1q and 2q impact NMD efficiency.**

**A-C**, For chromosome 1 (A, focusing solely on ASE iNMDeff) and chromosome 2 (B and C for ETG iNMDeff and ASE iNMDeff, respectively), gene-level iNMDeff scores are determined by the median value across individuals presenting focal CNAs within each gene, plotted according to genomic location. The top section of each panel illustrates the pan-cancer CNA frequency per gene within TCGA, distinguishing between higher confidence CNAs (GISTIC scores ≥ 1.5) and lower confidence CNAs (scores > 0). The middle section visualizes individual gene-wise iNMDeff scores, where the depth of red shading correlates with the frequency of co-amplification alongside the NMD factor gene *SMG5* (for chromosome 1, A) or *SF3B1* (for chromosome 2, B-C). The bottom section provides a detailed view of specific regions of interest—1q21.1-23.1 for chromosome 1 and 2q31.1-2q36.3 for chromosome 2—underscoring a noticeable decrease in iNMDeff in cases of region amplification. “Candidate” causal genes, “NMD” factors or related genes, and the rest of the genes (“Other”) are color-highlighted.

#### Supp. Fig. S15

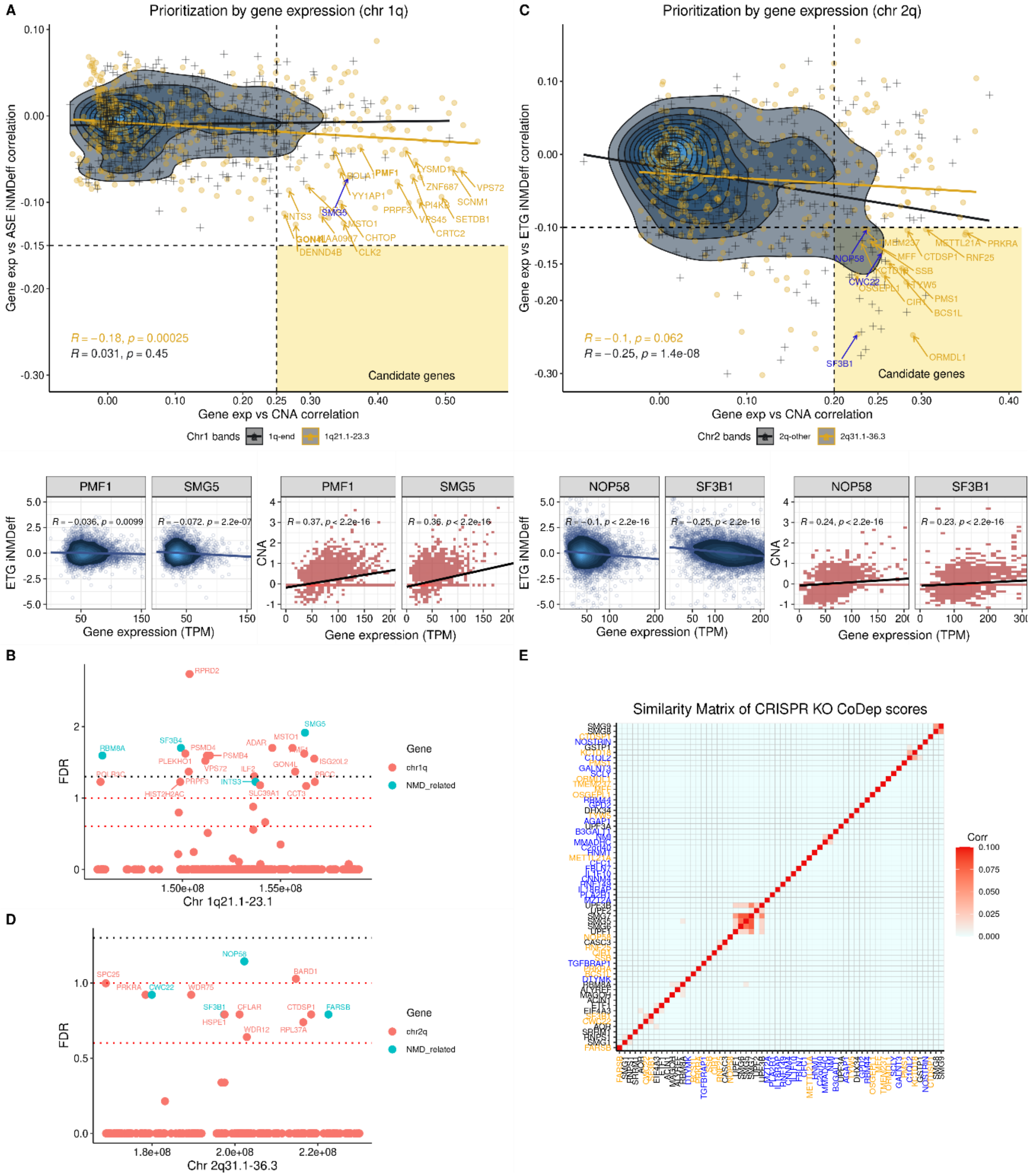


**Supp. Fig. S15. Prioritization of candidate genes from the chromosomal arms 1q and 2q, which exhibit CNA gains.**

**A and C**, Prioritization of candidate genes from chromosomes 1q and 2q, respectively, based on the correlation between gene expression and iNMDeff (Y-axis), measured by ASE in chr1q (A) and ETG in chr2q (C) methods, juxtaposed against CNA amplification (X-axis). The genes situated within the focal regions—1q21.1-23.1 for chr1 and 2q31.1-2q36.3 for chr2—are highlighted in orange, NMD-related genes are in blue, with the rest depicted in black. The anticipated candidates reside in the quadrant underscoring a negative correlation between gene expression and iNMDeff, alongside a positive correlation with CNA amplification, demarcated by vertical and horizontal dashed lines at set thresholds for the two scores respectively. This analytical approach is further illustrated with specific examples, including *PMF1* and NMD-factor *SMG5* for chr1q, and *NOP58* and NMD-related *SF3B1* for chr2q, showcasing their gene expression relative to iNMDeff (bottom left) and CNA amplification (bottom right). **B and D**, Test calculating the mean of CRISPR co-dependency scores, comparing candidate genes from chr1q (B) or 2q (D) with 10 core NMD factor genes versus random control genes from the same chromosomes. Genes with significant co-dependencies are highlighted, illustrated by adjusted *p*-values (Y-axis) across various FDR thresholds using horizontal dashed-lines: 5% (black), 10% and 25% (red). *P*-values are obtained through a one-sided Mann-Whitney *U* test. X-axis represents the genome location. **E**, Clustering heatmap of CRISPR co-dependency scores for 18 candidate genes (orange), 23 NMD-related genes (blue), and 22 random controls (black) from chr2q, particularly focusing on the region 2q31.1-2q36.3. The normalization method, RPCO (*onion*), transforms the scores into an affinity matrix, where higher scores indicate a stronger genetic association.

**Supp. Fig. S16**

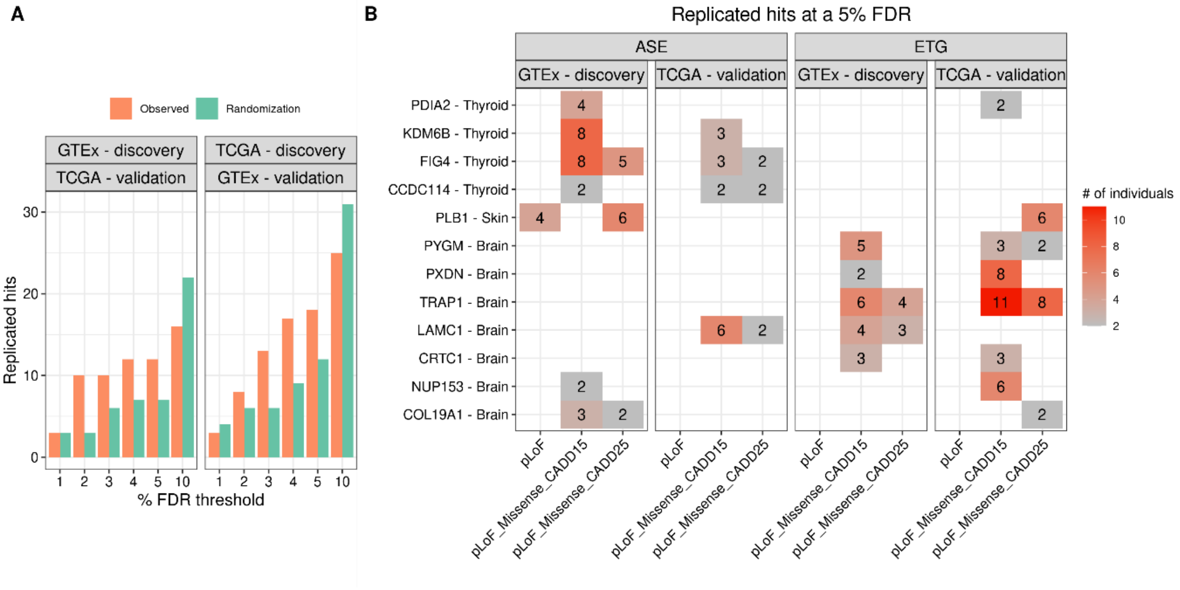


**Supp. Fig. S16. Rare deleterious germline variants are associated with NMD efficiency.**
**A,** Illustrates the number of replicated hits (Y-axis), differentiated by the cohorts TCGA or GTex and whether the direction is from discovery to validation or vice versa on top panels, across varying FDR thresholds from 1 to 10% (X-axis). The orange bars depict observed hits, while the green bars represent hits from randomizations of iNMDeff values, offering insights into the robustness of the associations against random expectations. **B,** Detailed view of replicated associations at 5% FDR. Shows replicated gene-tissue pairs (Y-axis) at a 5% FDR threshold, categorized by pLoF variant set (X-axis), NMD method, and the cohort where the association was identified (either discovery or validation). The values indicate the total count of individuals harboring a rare pLoF variant within the successfully replicated genes.

#### Supp. Fig. S17

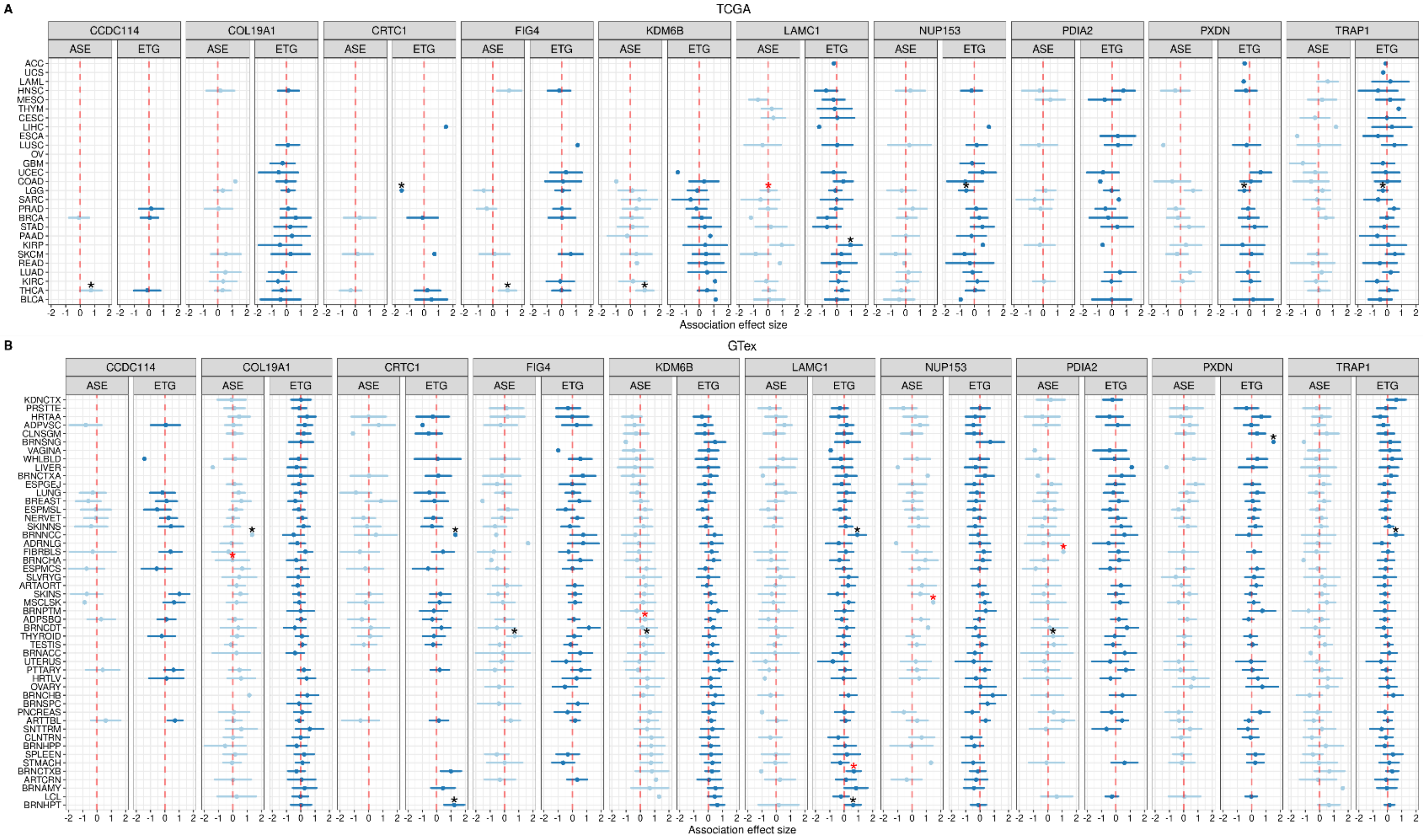


**Supp. Fig. S17. Gene burden test associations of rare germline pLoF variants with iNMDeff for 10 replicated hits**

**A-B**, Effect sizes (beta coefficients) are derived from linear model associations on the X-axis against specific cancer types (TCGA, A) and normal tissues (GTex, B) on the Y-axis, respectively. The analysis is splitted by the NMD method utilized, with ASE method results in left panels and ETG method in right panels. Displays 95% confidence intervals to indicate the precision of the effect size estimates, and an asterisk signifying FDR < 5%, which highlights statistically significant associations.

**Supp. Fig. S18**

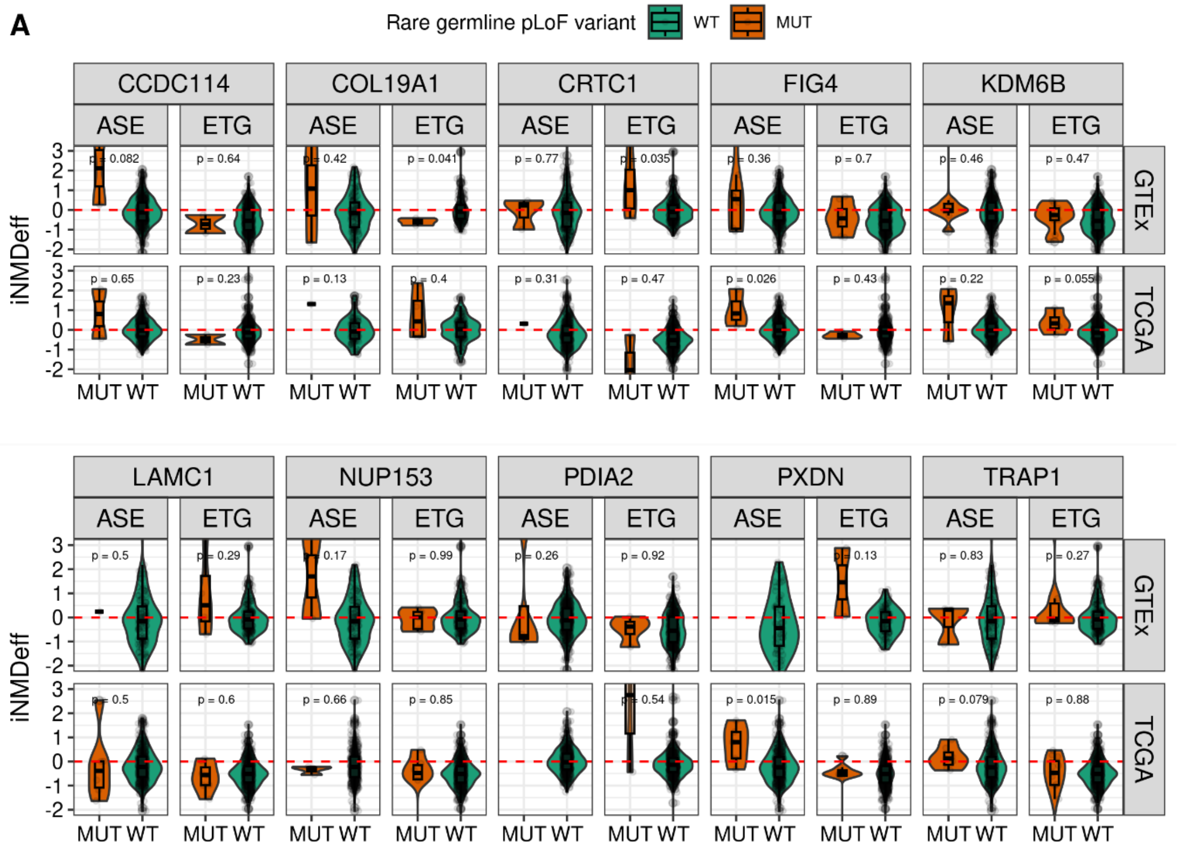


**Supp. Fig. S18. iNMDeff of individuals with and without rare pLoFs on the 10 replicated hits from the RVAS analysis**

**A,** iNMDeff (Y-axis) between individuals harboring wild-type (WT) alleles and those with rare pLoF mutations (MUT) within the 10 genes identified as significant in the RVAS analysis. These genes include *CCDC114*, *COL19A1*, *CRTC1*, *FIG4*, and *KDM6B* showcased in the top panels, and *LAMC1*, *NUP153*, *PDIA2*, *PXDN*, and *TRAP1* depicted in the bottom panels. The analysis spans both TCGA and the GTex cohorts and is further distinguished by the method of NMD efficiency estimation, either ASE or ETG. Statistical significance between the WT and MUT groups is assessed using two-sided Mann-Whitney *U* tests, when applicable.

**Supp. Fig. S19**

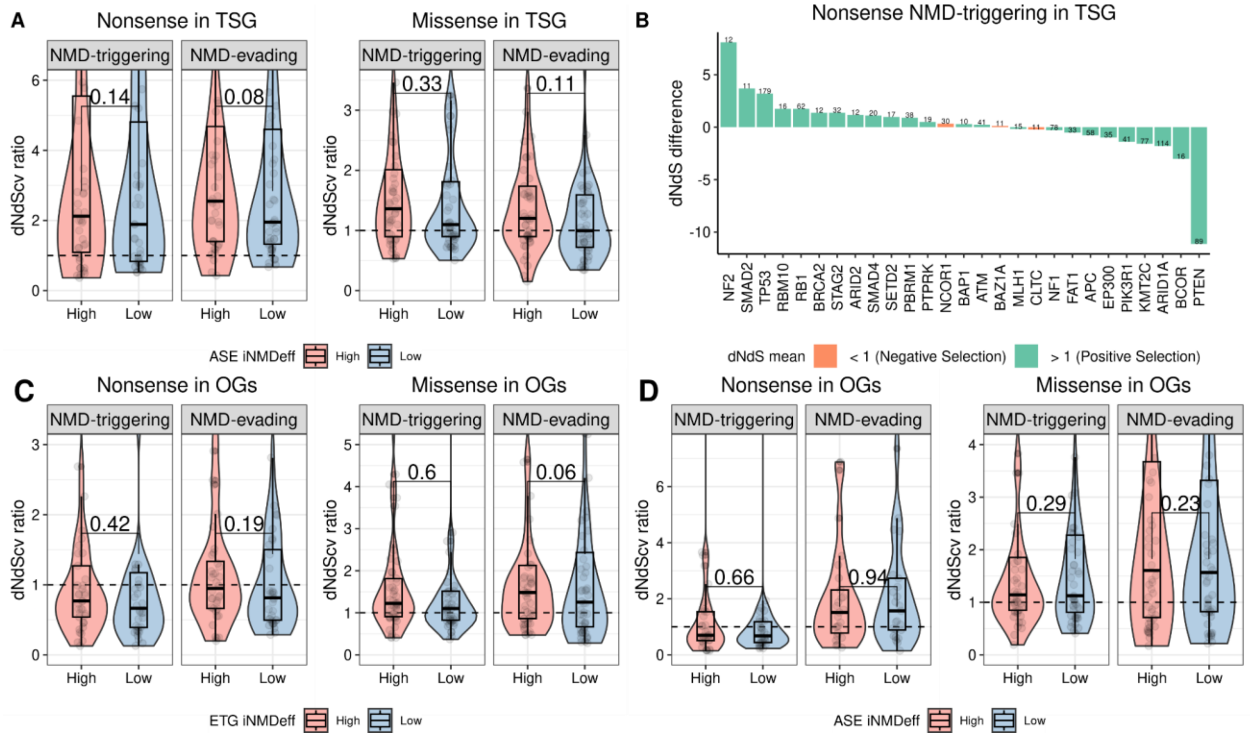


**Supp. Fig. S19. NMD efficiency modulates selection of somatic nonsense mutations.**
**A**, The dNdS ratios (Y-axis) of tumor suppressor genes (TSGs) for both NMD-triggering and NMD-evading nonsense (left panel) and missense (right panel) somatic mutations are compared between two groups of individuals with High (red) and Low (blue) iNMDeff, as determined by the median ASE iNMDeff. Statistical significance is assessed using one-sided Mann-Whitney tests on paired data. **B**, The differences in dNdS ratios (Y-axis) between High and Low iNMDeff groups are plotted for each gene (X-axis), specifically for NMD-triggering nonsense mutations within TSGs. Positive values indicate a stronger selection pressure in the high iNMDeff group compared to the low group, and vice versa. The number above each bar denotes the total count of nonsense mutations contributing to the dNdS calculation for both groups. Genes with less than 10 counts were removed. The barplot colors correspond to the mean dNdS ratios of the two iNMDeff groups (high and low): green indicates a mean ratio greater than 1, suggesting positive selection, while orange signifies a mean ratio less than 1, indicative of lack of positive selection. **C-D**, Same as A but for oncogenes (OGs) using either ETG (C) or ASE (D) iNMDeff individuals.

**Supp. Fig. S20**

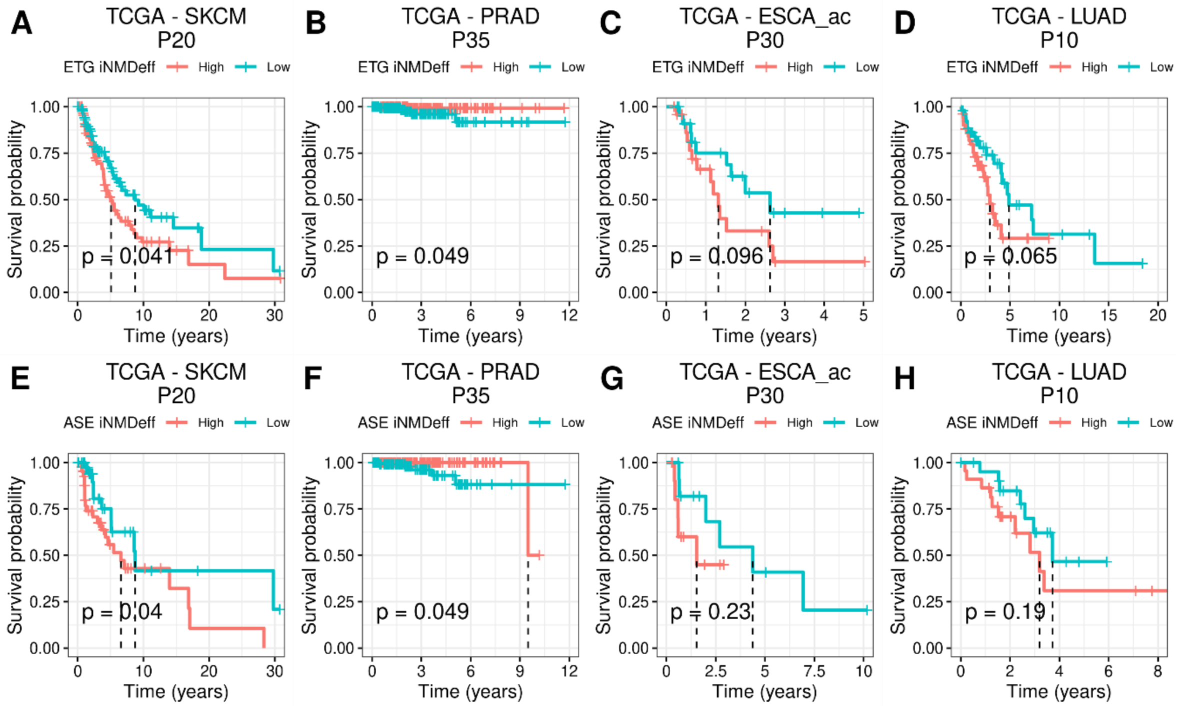


**Supp. Fig. S20. NMD efficiency impacts overall survival (OS) in diverse cancer types.**
**A-D,** Kaplan-Meier (KM) survival curves comparing overall survival (OS) outcomes between groups with High (red) versus Low (blue) iNMDeff, as determined by the median of ETG iNMDeff in TCGA cancer types: SKCM (percentile 20th), PRAD (percentile 35th), ESCA_acc (percentile 30th) and LUAD (percentile 10th). **E-H**, Same as A-D but the groups are determined by the median ASE iNMDeff. Log-rank test p-values quantify the statistical significance of the survival differences for all KM curves (A-H)**.**

**Supp. Fig. S21**

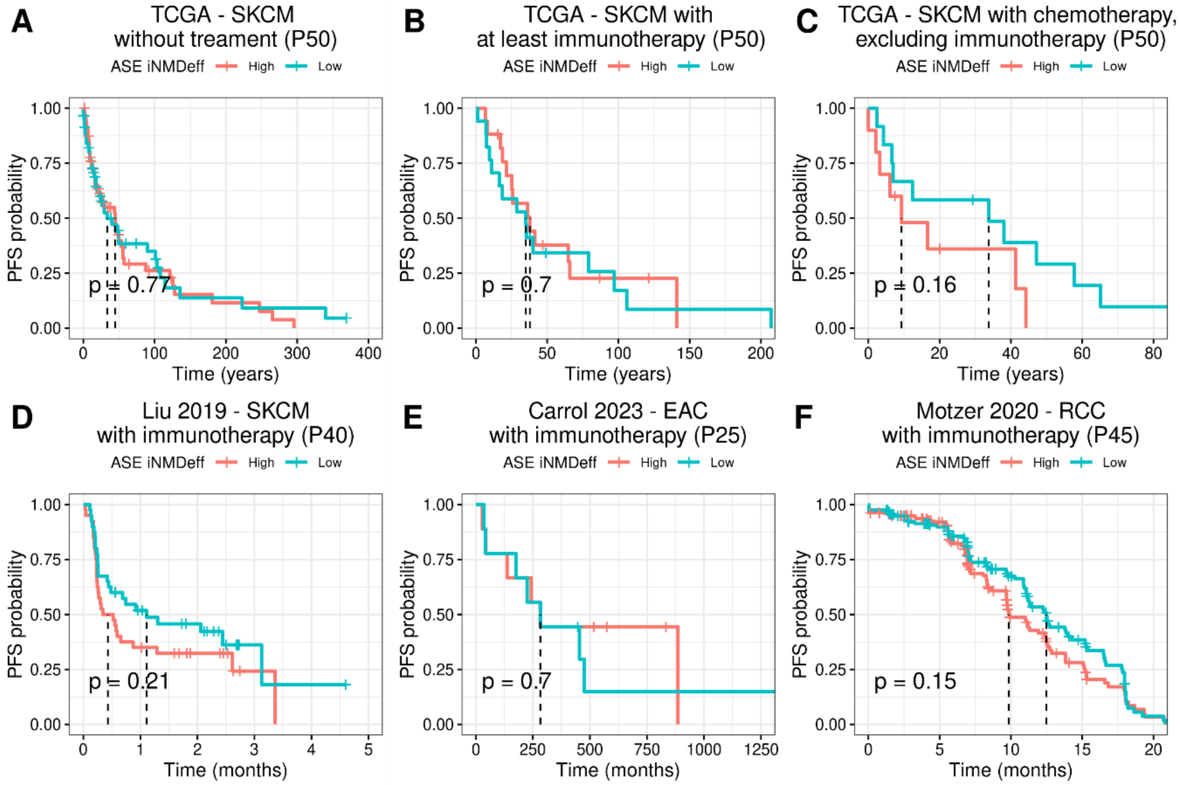


**Supp. Fig. S21. NMD efficiency impacts progression-free survival (PFS) and response to immunotherapy in treated patients**

**A-C,** Kaplan-Meier (KM) survival curves comparing progression-free survival (PFS) outcomes between groups with High (red) versus Low (blue) iNMDeff, as determined by the median of ASE iNMDeff in TCGA SKCM. The analysis is divided into three patient categories: untreated (A), those treated with immunotherapy at least once (B), and those treated exclusively with chemotherapy, excluding immunotherapy patients (C). Log-rank test p-values quantify the statistical significance of the survival differences. **D-F,** Validation of the KM curves for PFS in external patient cohorts from Liu et al. 2019 (SKCM, D) at the 40th percentile, Carrol et al. 2023 (EAC, E) at the 25th percentile, and Motzer et al. 2020 (RCC, F) at the 45th percentile. Patients treated with Sunitinib were excluded from this analysis.

### Supplementary Tables

**Supp. Table S1. Performance of linear regression models to predict iNMDeff.**
A stepwise removal procedure was used to evaluate the impact of each variable on the model's overall explanatory power, specifically measuring the reduction in the total variance explained ($R^2$) by the model.

**Supp. Table S2.** Complete names and acronyms for cancer types stratified into subtypes (TCGA) and normal tissues (GTex).

**Supp. Table S3.** Impact of iNMDeff on overall survival: significant findings from Cox proportional hazards regression models across cancer types

**Supp. Table S4.** Impact of iNMDeff on progression-free survival in immunotherapy-treated patients: significant findings (FDR < 10%) from Cox proportional hazards regression models across cancer types.

**Supp. Table S5.** Impact of iNMDeff on progression-free survival in chemotherapy-treated patients: significant findings (FDR < 10%) from Cox proportional hazards regression models across cancer types.

**Supp. Table S6.** Impact of iNMDeff on progression-free survival in chemotherapy-treated patients: significant findings (FDR < 25%) from Cox proportional hazards regression models across cancer types.

**Supp. Table S7.** Impact of iNMDeff on progression-free survival in immunotherapy-treated patients for the external validation datasets applying Cox proportional hazards regression models.

**Supp. Table S8.** Performance improvement of logistic regression models for predicting immunotherapy response across cancer types and validation datasets by including iNMDeff.
